# Targeting Fibrotic Scars with Extracellular Vesicles Extracted from Mature *Aloe vera* Gel: Enhanced Antioxidant, Anti-inflammatory, and Antifibrotic Activity through M2 Macrophage Polarization and Inhibition of Myofibroblast Differentiation

**DOI:** 10.1101/2025.06.17.660207

**Authors:** M. Camila Ceballos-Santa, Janitri Venkatachala-Babu, Thomas R. Gaborski, Stefan Schulze, Karin Wuertz-Kozak

## Abstract

**Background:** Abnormal scarring and fibrotic skin disorders arise from dysregulated wound healing. While *Aloe vera* is widely recognized for its therapeutic properties, the potential of its extracellular vesicles (Av-EVs) remains underexplored.

**Objective:** To isolate and characterize Av-EVs and evaluate their antioxidant, anti-inflammatory, and antifibrotic properties *in vitro*, focusing on extraction method and plant maturity.

**Methods:** Av-EVs were isolated from the gel of mature and young *Aloe vera* leaves using manual (NB) or blender-based (B) homogenization and characterized by nanoparticle tracking analysis, transmission electron microscopy, protein quantification, and immunoblotting of EV-associated markers. Antioxidant and cytotoxicity assays (DPPH, alamarBlue) were followed by functional anti-inflammatory and antifibrotic analyses in LPS-stimulated THP-1 macrophages and TGF-β1/vitamin C-activated human dermal fibroblasts (RT-qPCR, immunofluorescence). Proteomic analysis was performed to investigate maturity-dependent differences in Av-EV cargo.

**Results:** Mature NB-derived EVs exhibited the strongest bioactivity, with superior antioxidant capacity, suppression of pro-inflammatory mediators, enhanced M2 macrophage polarization, and significant downregulation of COL1A1 and α-SMA. Immunoblotting further supported Av-EV identity and sample purity. B-derived and young EVs showed reduced bioactivity, with the latter failing to inhibit fibrotic markers.

**Conclusion:** Manually extracted Av-EVs from mature leaves demonstrate superior multifunctional bioactivity, supporting their potential as plant-derived nanotherapeutics for fibrotic scar modulation.

## 1. Introduction

Disruptions in the tightly regulated wound healing process – hemostasis, inflammation, proliferation, and remodeling – can lead to pathological outcomes, ranging from chronic nonhealing wounds to excessive fibrotic scarring. The latter is characterized by aberrant collagen accumulation, disorganized extracellular matrix (ECM) remodeling, and persistent myofibroblast activation driven by abnormal fibroblast proliferation and transdifferentiation.^[1]^ Among the principal causes of abnormal wound healing are chronic inflammation and oxidative stress, which induce an imbalance of oxygen- and nitrogen-based free radical production and the cellular antioxidant defense system.^[2]^ Excess reactive oxygen species (ROS) and reactive nitrogen species (RNS) not only damage cellular macromolecules such as lipids, proteins, and DNA, but also perpetuate pro-inflammatory signaling and impair tissue regeneration.^[3]^ In this dysregulated microenvironment, ROS/RNS activate latent transforming growth factor-beta (TGF-β), which in turn promotes fibroblast-to-myofibroblast differentiation and excessive secretion of collagen and α-smooth muscle actin (α-SMA) – a critical protein for wound contraction. These molecular events ultimately result in the formation of fibrotic scar tissue, clinically recognized as hypertrophic scars and keloids.^[4, 5]^

To mitigate such outcomes, *Aloe vera* has been extensively studied for its diverse pharmacological properties and high concentration of bioactive constituents.^[6–8]^ This succulent plant harbors over 200 compounds and 75 nutrients, including soluble sugars, lipids, proteins, polysaccharides and lignin, polyphenols, enzymes, vitamins (e.g., A, C, E, B_1_, B_2_, B_12_) and minerals (e.g., calcium, potassium, sodium, magnesium),^[9]^ which collectively confer its wound healing, anti-inflammatory, antioxidant, antimicrobial, immunomodulatory, analgesic, and moisturizing benefits.^[10, 11]^

Historically, *Aloe vera* research has centered on topical and oral formulations. A large body of evidence, including *in vitro* studies, preclinical models, and human clinical trials, has consistently demonstrated the therapeutic efficacy of *Aloe vera*-based products.^[12–29]^ However, a promising new frontier in this research has emerged with the exploration of plant-derived extracellular vesicles (PDEVs) – nanoscale membranous particles secreted by plant cells that not only retain the therapeutic attributes of their source tissue but also offer target delivery capabilities. By delivering a diverse cargo of lipids, nucleic acids, proteins, metabolites, and even pharmacological compounds, PDEVs can mediate intracellular and intercellular communication, maintain tissue homeostasis, and support regenerative processes.^[30]^

Compared to their mammalian counterparts, PDEVs offer multiple advantages beyond their typical mechanism of action, such as being ethically uncontroversial, cost-effective and scalable, and evidence thus far suggests that they do not pose risk of carcinogenesis, immunogenicity, or tissue rejection. Additionally, they overcome key limitations of synthetic and mammalian EV-based drug delivery systems, including low yield, short bioavailability, poor loading efficiency, and limited tissue targeting.^[30–34]^ For these reasons, PDEVs are now considered as a new class of cell-free therapeutic platforms with high potential for clinical translation and preventive healthcare.

Among PDEVs, *Aloe*-derived EVs have recently moved into the spotlight for their ability to resemble the plant’s therapeutic properties while acting as efficient, biocompatible nanocarriers. Preliminary studies suggest that they can regulate inflammatory signaling, activate antioxidant defenses, and prevent the differentiation and contractile activity of myofibroblasts.^[35–37]^ Their versatility is further exemplified by their applications in indocyanine green encapsulation for targeted melanoma therapy.^[38]^ However, there remains a significant lack in the literature regarding the influence of extraction methodology and plant maturity on the biological efficacy of *Aloe*-derived EVs, and no *in vitro* study has yet evaluated their concurrent ability to modulate oxidative stress, inflammatory responses, and fibrotic remodeling within the context of abnormal skin scarring.

Therefore, in this study, we report a comprehensive biological characterization of *Aloe vera* gel-derived extracellular vesicles (Av-EVs), with a focus on their capacity to modulate key pathological mechanisms underlying abnormal scar formation. Specifically, we investigated the ability of Av-EVs to attenuate pro-inflammatory cytokine expression in M1-polarized macrophages while promoting M2-like phenotype, suppress myofibroblast differentiation in human dermal fibroblasts, and reduce free radical levels in an antioxidant assay – three interrelated processes central to fibrotic remodeling. To deepen our understanding of their therapeutic value, we systematically compared Av-EVs isolated via manual extraction (no blender, NB) versus shear-force homogenization (blender, B) and evaluated the influence of leaf maturity by comparing EVs derived from mature *Aloe vera* leaves (commercially-grade) and younger leaves from nursery-grown plants. This comparative framework provides critical insights into how source material and processing techniques affect the physicochemical and functional properties of Av-EVs, and positions them as promising, plant-based nanotherapeutics for the prevention and treatment of fibrosis-related skin disorders.

## 2. Materials and Approach

### 2.1 Av-EVs Isolation and Characterization

Comprehensive details regarding Av-EV isolation and characterization are available in our peer-reviewed published protocol called “A Step-By-Step Protocol for the Isolation of *Aloe vera*-Derived Extracellular Vesicles via Manual and Shear-Force Homogenization”.^[39]^

#### 2.1.1 Av-EVs Isolation from Aloe vera Gel

Fresh *Aloe vera barbadensis miller* leaves from mature plants (Melissa’s/World Variety Produce, Inc., USA) with similar appearance and dimensions (∼67 cm long x ∼9 cm wide x ∼3 cm thick) were purchased from a local market. These mature leaves were cut and rinsed with distilled water until the mucilage was removed. Subsequently, the pointed ends and the anterior leaf skin were chopped with a sterilized knife to separate it from the transparent inner gel. After peeling, the extraction and homogenization of the *Aloe vera* gel was carried out employing a manual method (no blender, NB Av-EVs) and a shear force-based (blender, B Av-EVs) method. For the manual method, the gel was meticulously scooped out using a sterilized spoon, avoiding any inclusion of the yellow latex. Approximately 400 mL of the extracted gel was transferred into a clean glass container where it was mixed with Dulbecco’s phosphate-buffered saline (DPBS, Cytiva, USA, SH30028.02) in a 1:1 v/v ratio and homogenized using an orbital shaker (Chemglass CLS-4021-100 Versa-Orb, USA) at 100 rpm and 4 °C for 1 h. Then, the remaining solid residues were separated with a sterilized fine-mesh sieve. On the other hand, the entire transparent inner portion was removed from the posterior leaf skin and homogenized with 1:1 v/v ratio of DPBS using a commercial blender for the shear force-based method to obtain approximately 800 mL of *Aloe vera* juice. In both cases, the soluble fractions were sequentially centrifuged (Centrifuge 5810, Eppendorf, USA) at 1,000 g for 10 min, 2,000 g for 20 min, 3,000 g for 30 min, and 15,000 g for 35 min at 4 °C to remove debris and larger particles. The supernatants were then ultracentrifuged (Optima^TM^ MAX-XP Ultracentrifuge, Beckman Coulter^®^, USA) at 100,000 g for 60 min at 4 °C and the resulting 6 pellets were resuspended and pooled in 10 mL of DPBS. The final suspensions containing the Av-EVs were filtered through a 0.22 µm filter (GVS, USA, FJ25BSCCA002AL01) and stored in aliquots of 1 mL at −80 °C until further use. The entire process was performed ∼4 – 5 times for each method, processing a total of 400 mL of *Aloe vera* gel and yielding four 10 mL batches of purified Av-EVs per ultracentrifugation cycle.

#### 2.1.2 Nanoparticle Tracking Analysis

Particle concentration and size distribution were measured using nanoparticle tracking analysis (NTA, NanoSight NS300, Malvern Panalytical, UK). The samples were taken from the Av-EV suspension aliquots and subsequently diluted 1:100 with DPBS to reach an optimal concentration for instrument linearity. Three videos of 30 s were captured with a camera level of 14 and measured with a detection threshold of 5. Data was processed with the NanoSight NTA 3.4 software.

#### 2.1.3 Transmission Electron Microscopy

Transmission electron microscopy (TEM) was employed to verify vesicle structure, for which 3 μL of the Av-EVs preparation were spotted on formvar/carbon-coated copper grids with a 200-mesh size (Electron Microscopy Science, USA) following glow discharge for 30 s at 30 mA and incubation for 30 s. The excess sample was blotted, and grids were rinsed sequentially with molecular grade water and counterstained with 0.75% uranyl formate solution for 1 min. Grids were allowed to air dry before imaging them on a Talos L120C TEM equipped with a 16 MP CETA camera at 8,500 and 22 k magnification using the TIA software (Thermo Fisher Scientific, USA).

#### 2.1.4 Protein Quantification

Pierce^TM^ BCA protein assay kit (Thermo Scientific, USA, 23225) was used to quantify protein concentration in Av-EV samples following the manufacturer’s instructions for the standard protocol and microplate procedure. For each protein quantification assay, a single EV pellet – obtained from one of the six ultracentrifugation pellets produced during one Av-EV isolation cycle – was individually resuspended in 300 µL of Pierce^TM^ RIPA lysis buffer (Thermo Scientific, USA, 89901) supplemented with 1× Halt™ protease and phosphatase inhibitor cocktail (Thermo Scientific, USA, 78444) and incubated on ice for 30 min. The lysate was then centrifuged at 14,000 g for 15 min at 4 °C (Centrifuge 5424 R, Eppendorf, USA) and the resulting supernatant was carefully collected and used for subsequent analysis. No pooling of multiple pellets was performed.

Briefly, the diluted bovine serum albumin (BSA, 23209) standards were prepared using 0 – 400 µL of RIPA for a working range of 2,000 – 25 µg/mL. The working reagent (WR) was prepared by mixing reagent A (23228) and B (1859078) at a 50:1 ratio until obtaining a clear, green WR for a total of 9 standards, 6 Av-EV independent samples (n=6) and 2 replicates for both isolation methods. 10 µL of each standard and Av-EVs samples were pipetted into a clear 96-well plate and 200 µL of the WR were added to each well. The plate was thoroughly mixed on an orbital shaker for 30 s, covered with aluminum foil to protect it from direct light, and incubated at 37 °C for 30 min. Afterward, the absorbance was read at 562 nm on a SpectraMax iD3^®^ spectrophotometer equipped with the Softmax Pro 7 software (Molecular Devices, USA). Results were normalized to the blank standard measurement of DPBS, and the protein concentration of each Av-EV sample was determined based on the BSA standard curve. Statistical difference was assessed with unpaired student’s t-test.

#### 2.1.5 Immunoblot Analysis of Plant-specific Markers

Western blot analysis was performed to assess the presence of EV-associated and non-EV protein markers in NB Av-EV preparations. Briefly, protein concentration from six pooled pellets generated during an ultracentrifugation cycle was determined using the Pierce™ BCA protein assay kit, as described in section

#### 2.1.4. Equal amounts of total protein (17 µg) were loaded per sample

Proteins were separated by SDS–PAGE using 4–20% precast polyacrylamide gels (Bio-Rad, USA, 4561094) and subsequently transferred onto nitrocellulose membranes (Bio-Rad, USA, 1704158). Membranes were blocked for 1 h at room temperature (RT) in 5% BSA (Thermo Scientific Chemicals, USA, AAJ6410009) prepared in Tris-buffered saline containing 0.1% Tween-20 (TBS-T), followed by overnight incubation at 4 °C with primary antibodies against the EV-associated markers TET8 (1:1000; PhytoAB, USA, PHY1491A) and SYP121/PEN1 (1:1000; Agrisera, USA, AS22 4827), as well as the negative marker Calnexin (1:100; Boca Scientific, USA, AS17 4164).

Following primary antibody incubation, membranes were washed with TBS-T and incubated for 1 h at RT in the dark with species-appropriate IRDye-conjugated secondary antibodies (LI-COR Biosciences, USA): IRDye 680RD goat anti-rabbit IgG (H+L) (1:10,000; 926-68071) for TET8 and SYP121, and IRDye 800CW goat anti-mouse IgG1 (1:10,000; 926-32350) for Calnexin. Protein bands were visualized using an Odyssey XF Imaging System (LI-COR Biosciences; acquisition software version 1.1) with near-infrared fluorescence detection.

### 2.2 Cell Cultures

#### 2.2.1 THP-1 Culture and Differentiation into M0 Macrophages

Human leukemia monocytic type 1 (THP-1) cells (The American Type Culture Collection (ATCC), USA) were cultured in Roswell park memorial institute medium (RPMI 1640, Cytiva, USA, SH30255.01) supplemented with 10% fetal bovine serum (FBS, Cytiva, USA, SH30070.03), 1% anti-anti (A/A, GibcoTM, USA, SV30079.01), and 1% sodium pyruvate (Gibco^TM^, CN, 11360-070) at 2 × 10^5^ cells/mL concentration under standard conditions (37 °C with 5% CO_2_). The cells were allowed to grow for two weeks with medium change every 2 – 3 days. Differentiation to macrophages (M0) was done by seeding the THP-1 cells into 24-well plates at 2 × 10^5^ cells per well and adding 100 ng/mL of phorbol 24-myristate 13-acetate (PMA, Cayman Chemical, USA, 16561-29-8) with standard medium for 24 h until the cells were attached and attained a macrophage-like morphology. Following adhesion, the corresponding treatment was added, and the specific assay was performed.

#### 2.2.2 hDFs Culture

Primary human dermal fibroblasts (hDFs) (ATCC, USA) were cultured in Dulbecco’s modified eagle medium (DMEM, Cytiva, USA, SH30023.01) supplemented with 10% FBS and 1% A/A under standard conditions. To support cell proliferation and maintenance, 5 ng/mL fibroblast growth factor (FGF, PeproTech, USA, 100-18B) was freshly added to the culture medium with each media change. Cells were fed every 2 – 3 days and passaged using 1X Trypsin-EDTA (Gibco™, CA, 15400-054) upon reaching approximately 80% confluence, and only passages ≤ 5 were used for experiments. For all assays, 2.5 × 10⁴ cells per well were seeded into 24-well plates and allowed to adhere for 24 hours prior to treatment administration.

For the gene expression analysis of cell responses to Av-EVs isolated from mature *Aloe vera* leaves, two independent hDF donors (lot numbers 80405992 and 80124232) were included. For each donor, all treatments were evaluated across three independent experimental replicates, with each replicate performed on a separate day and plate using an independently prepared Av-EV isolation sample. Thus, the main analyses are presented using the combined measurements from both donors (n = 6 per treatment group) to facilitate direct visualization and comparison of the effects of EV dose and isolation method. Importantly, donor-specific responses were also evaluated separately to account for potential inter-donor variability. As the donor-resolved analysis supported the same overall biological response, particularly the greater antifibrotic activity of NB Av-EVs at the intermediate and high doses, the combined dataset was retained in the main manuscript for clarity, while the complete donor-specific analysis is provided in the Supplementary Information [Figure 1S].

For the cytotoxicity assessment, cellular uptake of Av-EVs, α-SMA immunofluorescence, and gene expression analyses of responses to NB Av-EVs isolated from young *Aloe vera* leaves, hDFs from lot number 80405992 were used. For collagen I immunofluorescence analysis, hDFs from lot number 80124232 were used.

### 2.3 Biological Effects of Av-EVs Cargo

#### 2.3.1 Antioxidant Capacity

The antioxidant activity of Av-EVs was evaluated by measuring their ability to scavenge the free 2,2-diphenyl-1-picrylhydrazyl (DPPH) radical, using a DPPH antioxidant assay kit (Dojindo, USA, D678) following the manufacturer’s protocol. Identical to the protein quantification assay, for each antioxidant analysis, a single EV pellet – obtained from one of the six ultracentrifugation pellets produced during one Av-EV isolation cycle – was individually resuspended in 300 µL of DPBS. No pooling of multiple pellets was performed.

Briefly, 20 µL of three (n=3) NB and B Av-EV samples obtained from separate isolation experiments under the same conditions were prepared at different concentrations (5 × 10⁸, 1 × 10⁹, 5 × 10⁹, and 2 × 10¹⁰ EVs/mL) and incubated at RT in the dark with 80 µL of assay buffer and 100 µL of DPPH working solution (WS) diluted in 99.5% molecular grade ethanol (Sigma-Aldrich, USA, SHBS1884). Additionally, DPPH + ethanol was used as negative control, and EVs in DPBS as sample blank. At time intervals of 1, 24, 48, and 72 h, absorbance was measured at 517 nm using the SpectraMax iD3® spectrophotometer. The DPPH radical scavenging activity (inhibition ratio %) of the samples was calculated using the following equation:

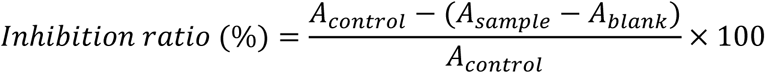

Statistical difference was determined using two-way analysis of variance (ANOVA) followed by Geisser-Greenhouse’s epsilon correction test, with concentrations and time points as fixed factors.

#### 2.3.2 Cytotoxicity Assessment

Cell viability was assessed using alamarBlueTM cell viability reagent (InvitrogenTM by Thermo Fisher Scientific, USA, DAL1025). THP-1 and hDFs were seeded in their respective standard medium and left to adhere for 24 h, as described in sections 2.2.1 and 2.2.2. Following attachment, cells were stimulated according to the protocols detailed in sections 2.3.3 and 2.3.4. to induce either M1 macrophage polarization (via LPS) or fibroblast to myofibroblast differentiation (via TGF-ꞵ1/vitamin C), respectively. Then, 350 µL of freshly made working alamarBlue solution diluted at 10% in each corresponding standard medium was added to each well at the indicated time points and allowed to incubate under standard conditions for 3 h. The plate was wrapped with aluminum foil to protect it from direct light. Next, 200 µL of the supernatant from each well was transferred to a 96-well black plate with clear bottom (Corning Incorporated, USA, 3603) and fluorescence levels of the reduced resazurin were read on a SpectraMax iD3® spectrophotometer equipped with the Softmax Pro 7 software (Molecular Devices, USA). The excitation/emission wavelengths were set at 560/590, respectively. Results were normalized with a blank of 10% alamarBlue without cells and the statistical analysis was performed using the respective untreated conditions (i.e., –LPS/–EVs for macrophages or –TGFꞵ1–VitC/–EVs for fibroblasts) as controls. Statistical difference was determined using one-way ANOVA followed by Tukey’s multiple comparison test.

#### 2.3.3 Anti-inflammatory Response (Macrophages)

Six replicates (n=6) of THP-1 were expanded and seeded as described in section 2.2.1. Once the cells were attached to the wells and differentiated, M0 macrophages were polarized to M1 macrophages through lipopolysaccharide (LPS, Sigma-Aldrich, USA, L6529-1MG) treatment. Fresh NB or B Av-EVs were added to serum-free medium simultaneously with 500 ng/mL of LPS and the treatment continued for 24 h, 72 h or 7 days. The experimental setup is specified in **Table 1**. D indicates different doses of EVs, where D1, D2, and D3 correspond to 1,000, 2,500, 5,000 Av-EVs per target cell, respectively. The term “EVs” is used throughout to describe particle concentrations measured by nanoparticle tracking analysis (NTA); however, it is important to note that not all particles quantified by NTA are necessarily extracellular vesicles. These measurements may include co-isolated nanoparticles such as protein aggregates or other vesicle-like structures. C-X-C motif chemokine ligand 10 (CXCL10) also known as Interferon gamma-induced protein 10 (IP-10), C-C motif chemokine ligand 2 (CCL2) also known as monocyte chemoattractant protein-1 (MCP-1), interleukin-1β (IL-1β), tumor necrosis factor alpha (TNF-α), interleukin-6 (IL-6), interleukin-10 (IL-10), and arginase 1 (ARG1) expression was quantified by RT-qPCR to analyze the anti-inflammatory potential of Av-EVs and their capacity to modulate macrophage polarization from M1 to M2 phenotype (see details in 2.4.1).

**Table 1.** Experimental setup for the anti-inflammatory response.

| Condition | Explanation |
| --- | --- |
| –LPS / –EVs | Untreated control |
| +LPS / –EVs | Pro-inflammatory positive control |
| –LPS / +NBEVs | NB Av-EVs control |
| –LPS / +BEVs | B Av-EVs control |
| +LPS / +NBEVs D1 | LPS and 1,000 NB AV-EVs / cell |
| +LPS / +NBEVs D2 | LPS and 2,500 NB AV-EVs / cell |
| +LPS / +NBEVs D3 | LPS and 5,000 NB AV-EVs / cell |
| +LPS / +BEVs D1 | LPS and 1,000 B AV-EVs / cell |
| +LPS / +BEVs D2 | LPS and 2,500 B AV-EVs / cell |
| +LPS / +BEVs D3 | LPS and 5,000 B AV-EVs / cell |

#### 2.3.4 Antifibrotic Response (Dermal Fibroblasts)

TGF-ꞵ1 (PreproTech, USA, 100-21) and vitamin C (L-Ascorbic acid, Sigma-Aldrich, USA, A8960-5G) treatment was used to induce hDFs transdifferentiation of fibroblasts to myofibroblasts, a recognized indicator of fibrosis. In brief, hDFs were expanded and seeded as described in section 2.2.2. Once the cells were attached to the wells, the standard medium was replaced with fresh medium with 10 ng/ml TGFꞵ1 + 100 µM vitamin C. Fresh NB or B Av-EVs were added simultaneously and the treatment continued for 72 h. The experimental setup is specified in **Table 2**. Identical to macrophages, D1, D2 and D3 were 1,000, 2,500, 5,000 NB or B Av-EVs per target cell, respectively. Unlike macrophages, hDFs continue proliferating during culture; therefore, EV doses were calculated based on the expected cell number after 72 h (approximately 5 × 10⁴ cells per well). The expression of collagen type I (COL1A1) and α-SMA, also known as actin alpha 2 (ACTA2) was quantified by RT-qPCR to analyze the antifibrotic potential of Av-EVs and their capacity to modulate myofibroblast differentiation. Additionally, immunofluorescence microscopy was used for α-SMA protein visualization (see details in 2.4.1 and 2.4.2).

**Table 2.** Experimental setup for the antifibrotic response.

| Condition | Explanation |
| --- | --- |
| –TGFβ1–VitC / –EVs | Untreated control |
| +TGFβ1+VitC / –EVs | Fibrotic positive control |
| –TGFβ1–VitC / +NBEVs | NB Av-EVs control |
| –TGFβ1–VitC / +BEVs | B Av-EVs control |
| +TGFβ1+VitC / +NBEVs D1 | TGFβ1/VitC and 1,000 NB AV-EVs / cell |
| +TGFβ1+VitC / +NBEVs D2 | TGFβ1/VitC and 2,500 NB AV-EVs / cell |
| +TGFβ1+VitC / +NBEVs D3 | TGFβ1/VitC and 5,000 NB AV-EVs / cell |
| +TGFβ1+VitC / +BEVs D1 | TGFβ1/VitC and 1,000 B AV-EVs / cell |
| +TGFβ1+VitC / +BEVs D2 | TGFβ1/VitC and 2,500 B AV-EVs / cell |
| +TGFβ1+VitC / +BEVs D3 | TGFβ1/VitC and 5,000 B AV-EVs / cell |

#### 2.3.5 Cellular Uptake of Av-EVs

To visualize the cellular uptake of Av-EVs, vesicles were labeled using the PKH26 red fluorescent membrane kit (Sigma–Aldrich, USA, 4103849473) (panel: red). After isolation, 10 mL of purified Av-EVs in DPBS were stored in aliquots at −80 °C as previously described in section 2.1.1. For PKH26 labeling, one 1 mL aliquot was thawed and ultracentrifuged at 100,000 g and 4 °C for 60 min. The pellet was then resuspended in 1 mL of Diluent C. Separately, 4 µL of PKH26 ethanolic dye solution was mixed in 1 mL of Diluent C until complete dispersion. Then, the Av-EVs suspension was mixed in the dye solution and incubated at RT for 10 min. An equal volume (2 mL) of 10% FBS in DMEM was added to stop the labeling reaction. Finally, the solution was ultracentrifuged, and the resulting pellet was washed with DPBS and ultracentrifuged one more time to remove free dye aggregates. The final pellet containing the labeled Av-EVs was resuspended in the initial volume of DPBS (1 mL), wrapped in aluminum foil to protect from light and stored at −20 °C until further use.

THP-1 and hDFs were seeded in their respective standard medium and left to adhere for 24 h, as described in sections 2.2.1 and 2.2.2. Following attachment, cells were stimulated according to the protocols detailed in sections 2.3.3 and 2.3.4. to induce either M1 macrophage polarization (via LPS) or fibroblast to myofibroblast differentiation (via TGF-ꞵ1/vitamin C), respectively. Simultaneously, cells were treated with PKH26-labeled Av-EVs at the highest dose (D3: 5,000 EVs/cell) and incubated for an additional 24 h to allow for vesicle uptake. Subsequently, the cells were fixed with 4% paraformaldehyde solution (Thermo Scientific, USA, J19943-K2) at RT for 15 min, permeabilized with 10X PBS-Triton X-100 (Alfa Aesar, USA, J63521) at RT for 15 min, and blocked with 4% BSA in DPBS at 4 °C for 2 h. From this point on, all procedures were carried out under low-light conditions to protect the fluorescent dyes. The Alexa Fluor^TM^ 488 phalloidin (Invitrogen, USA, A12379) was diluted 1:400 in 4% BSA and added to the cells to stain the actin filaments (F-actin) at RT for 30 min (panel: green), followed by Hoechst 33342 (Thermo Scientific, USA, H3570) staining to label cell nuclei at a dilution of 1:3,000 in DPBS at RT for 5 min (panel: blue). Finally, the cells were washed and visualized under fluorescence microscopy (Olympus IX81, USA) to confirm the cellular uptake of PKH26-labeled Av-EVs and evaluate cytoskeletal and nuclear features. The images were captured at 20X, 40X or 60X magnifications and processed using FIJI (Image J) software.

### 2.4 Analysis of Cell Responses to Av-EVs from Mature *Aloe vera* Leaves

#### 2.4.1 Gene Expression

Reverse transcription-quantitative polymerase chain reaction (RT-qPCR) was employed to investigate the anti-inflammatory potential of Av-EVs using THP-1 macrophages stimulated with LPS by analyzing the gene expression of human pro-inflammatory (CXCL10/IP-10, CCL2/MCP-1, IL-1β, TNF-α, IL-6) and anti-inflammatory (IL-10) cytokine, as well as the expression of ARG1, an enzyme highly expressed in M2 macrophages. Additionally, RT-qPCR served to investigate the hDFs transdifferentiation into myofibroblast after treatment with TGF-β1 and vitamin C by analyzing the gene expression of COL1A1 and α-SMA/ACTA2. The ID of each primer can be found in **Table 3**.

**Table 3.**
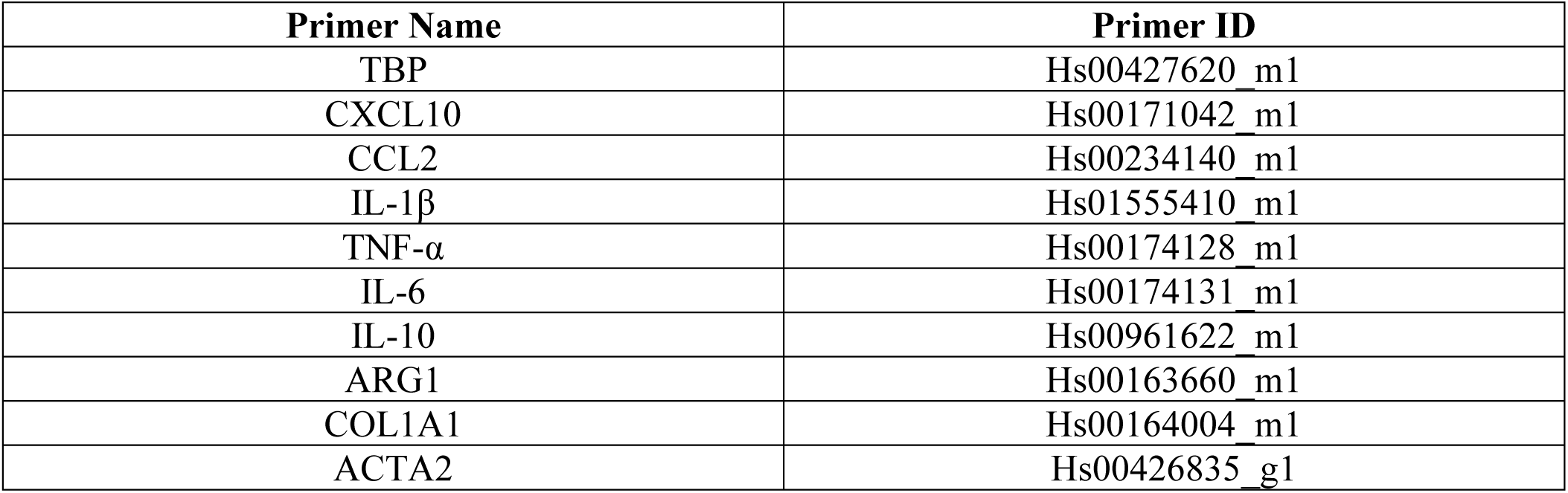
TaqMan^TM^ gene expression assays used for RT-qPCR analysis.

Total RNA was extracted from THP-1-derived macrophages and hDFs using the RNeasy mini kit (Qiagen, DE, 74106) according to the manufacturer’s protocol. RNA concentration and purity were assessed spectrophotometrically using a NanoPhotometer® N50 (Implen, DE), and yields were expressed in ng/μL. Subsequently, RNA was reverse transcribed into complementary DNA (cDNA) using the high-capacity cDNA reverse transcription kit with RNase inhibitor (Applied Biosystems™ by Thermo Fisher Scientific, USA, 4374967) on a T100™ thermal cycler (Bio-Rad, USA). Quantitative real-time PCR (qPCR) was then performed using TaqMan™ fast advanced master mix (Applied Biosystems™, USA, 4444557) with gene-specific primers (Applied Biosystems™, USA, 4331182) on a QuantStudio™ 3 Real-Time PCR System (Applied Biosystems™, USA). Gene expression levels were quantified by the ΔΔCt method represented in terms of fold change (2^−ΔΔCt^). The TATA-binding protein (TBP) gene served as the endogenous reference. Experimental conditions were normalized to their respective stimulated controls: +LPS/–EVs for pro-inflammatory assays and +TGF-β1+VitC/–EVs for fibrotic assays. All qPCR reactions were conducted in duplicate for each gene and treatment group.

Statistical significance was determined using one-way ANOVA followed by Tukey’s multiple comparison test for both THP-1 and hDF experiments, as presented in the main manuscript.

For hDF gene expression experiments evaluating Av-EVs isolated from mature *Aloe vera* leaves, data from the two biological donors were additionally analyzed using a mixed-effects model (REML) to account for the hierarchical structure of the experimental design and potential inter-donor variability. In this analysis, treatment was specified as a fixed effect and donor as a random effect, with three independent experimental replicates nested within each donor. The complete mixed-effects analysis, including donor-specific responses across treatment conditions, is provided in the Supplementary Information [Figure 1S].

#### 2.4.2 Collagen type I and Smooth Muscle Actin Immunofluorescence Analysis

To analyze collagen type I and α-SMA protein expression in fibrotic hDFs treated with the intermediate dose (2,500 EVs/cell) of either NB-or B-derived Av-EVs (as described in section 2.3.4), immunofluorescence staining was performed. The cells were seeded, differentiated, fixed, permeabilized and blocked as described in section 2.3.5. Then, the cells were incubated separately with collagen I antibody (Novus Biologicals, USA, NB600-408) at a dilution of 1:200 or with alpha-smooth muscle actin monoclonal antibody (1A4), eBioscience™ (Invitrogen, USA, 14-9760-82) at 1 µg/mL in 0.1% BSA at 4 °C overnight. The next date, the cells were labeled with goat anti-rabbit IgG (H+L) superclonal™ secondary antibody, Alexa Fluor 488^TM^ conjugate (Invitrogen, USA, A27034) or with goat anti-mouse IgG (H+L) superclonal™ secondary antibody, Alexa Fluor 488^TM^ conjugate (Invitrogen, USA, A28175), respectively, both at a dilution of 1:200 at RT for 45 min (panel: green). Afterward, F-actin was stained with Alexa Fluor^TM^ 594 phalloidin (Invitrogen, USA, A12381) diluted 1:400 in 4% BSA at RT for 30 min (panel: red), and nuclei were stained with Hoechst 33342 at RT for 3 min (panel: blue). For each condition, three independent images were acquired at 20X magnification under identical exposure settings to enable objective comparison of fluorescence intensity. From each image, two regions of interest were analyzed, yielding a total of six collagen I or α-SMA, and six background pixel intensity measurements per group. Quantification was performed using FIJI (ImageJ) software by calculating net collagen I or α-SMA signal intensity as the difference between each measurement and the average background value. To enhance visualization while preserving quantitative accuracy, uniform brightness and contrast adjustments were applied post-capture across all images. Statistical significance was determined using one-way ANOVA followed by Tukey’s multiple comparison test.

### 2.5 Analysis of Cell Responses to NB Av-EVs from Young *Aloe vera* Leaves

#### 2.5.1 Anti-inflammatory (Macrophages) and Antifibrotic (Dermal Fibroblasts) Responses

To assess the influence of plant maturity on the biological activity and protein composition of Av-EVs, we extended our investigation to include EVs manually extracted from young *Aloe vera barbadensis miller* leaves. Six leaves (n=6) were harvested from live potted plants purchased at a local plant shop and the EVs were isolated using the same manual protocol previously described in section 2.1.1. Following isolation, the anti-inflammatory potential of young NB Av-EVs was evaluated in THP-1-derived macrophages after 24 hours of co-treatment, and the antifibrotic response was assessed in hDFs (see details in 2.3.3 and 2.3.4, respectively). Gene expression of CXCL10/IP-10, CCL2/MCP-1, IL-1β, TNF-α, IL-6, IL-10, COL1A1 and α-SMA/ACTA2 was quantified by RT-qPCR (see details in 2.4.1), with all reactions performed in duplicate for each gene and treatment group. The ID of each primer can be found in **Table 3**. Statistical significance was determined using one-way ANOVA followed by Tukey’s multiple comparison test.

#### 2.5.2 Proteomic Characterization via Label-Free Data-Dependent Acquisition Mass Spectrometry

To investigate molecular differences in vesicle cargo, label-free data-dependent acquisition (DDA)-based proteomic profiling was carried out on Av-EVs derived from both mature and young *Aloe vera* leaves. Each group consisted of three biological replicates (n=3), where the mature EV samples were isolated from three independent leaves, and the young EV samples were obtained from three separate potted plants. Following ultracentrifugation, EV pellets were directly resuspended in Pierce™ RIPA lysis buffer (Thermo Scientific, USA, 89901), and protein concentration was determined using the BCA assay as described in section 2.1.4.

Samples were submitted to MtoZ Biolabs (USA) for protein extraction, LC-MS/MS analysis, and downstream bioinformatics employing a transcriptome-informed *Aloe vera* database (∼77,000 protein sequences)^[40]^. In their facilities, proteins were reduced with 10 mM dithiothreitol at 56 °C for 1 h, alkylated with 55 mM iodoacetamide at RT in the dark for 1 h, and digested overnight with sequencing-grade trypsin (Promega) in 50 mM NH₄HCO₃ at 37 °C, using the previously described SP3 workflow.^[41]^ Peptides were desalted using C18 columns, vacuum-dried at 45 °C, and resuspended in 0.1% formic acid. Peptide separation was carried out using an Easy-nLC 1200 system coupled to an Orbitrap Exploris 480 (Thermo Fisher Scientific) with a 66-minute gradient on a C18 column (150 μm i.d. × 170 mm, 1.9 μm, 100 Å). Mass spectrometry was performed in positive ion mode using a top-20 data-dependent acquisition method with a full scan range of m/z 400–1200 at 60,000 resolutions. Peptide fragmentation was achieved using high-energy collision dissociation at a normalized collision energy of 30%. Raw MS/MS data were analyzed against *Aloe vera* specific database that was generated based on the transcriptomics data described by Choudhri et al.^[40]^ Briefly, unigenes were downloaded from Bioproject PRJNA359629, and coding sequences were predicted using TransDecoder^[42]^ implemented in Galaxy,^[43]^ resulting in 76,958 protein sequences. Search parameters included up to two missed cleavages, fixed carbamidomethylation of cysteines, and variable modifications including methionine oxidation and N-terminal acetylation. A 1% false discovery rate was applied at both peptide and protein levels. Quantitative data were median-normalized and missing values imputed using the K-nearest neighbor (KNN) algorithm. Bioinformatics analyses included differential protein expression, principal component analysis (PCA), hierarchical clustering, gene ontology (GO), Kyoto encyclopedia of genes and genomes (KEGG) pathway enrichment, and protein–protein interaction (PPI) network mapping.

### 2.6 Statistical Analysis

All experiments involving Av-EVs derived from either mature or young *Aloe vera* leaves were performed using EV preparations obtained from independent leaves and isolation cycles. Protein quantification (n=6) was analyzed using an unpaired two-tailed Student’s t-test, whereas antioxidant capacity assays (n=3) were evaluated using two-way ANOVA with concentration and time as factors, followed by Geisser– Greenhouse correction.

For THP-1-based assays, n=6 represents experimental replicates initiated from separate frozen cell stocks, with cells independently thawed, expanded, seeded, and treated. Cytotoxicity and gene expression data were analyzed using one-way ANOVA followed by Tukey’s multiple comparison test.

For primary hDF-based assays, including cytotoxicity assessment, collagen I and α-SMA immunofluorescence, and gene expression analysis of responses to NB Av-EVs isolated from young *Aloe vera* leaves, one biological donor was used, and statistical significance was determined using one-way ANOVA followed by Tukey’s multiple comparison test. Gene expression analysis of responses to Av-EVs isolated from mature *Aloe vera* leaves included two biological donors, with three independent experimental replicates per donor for each treatment condition. Each experimental replicate was performed on a separate day and plate using an independently prepared Av-EV isolation sample.

For the main manuscript, data from both donors are presented together and were analyzed using one-way ANOVA followed by Tukey’s multiple comparisons test to facilitate direct comparison among treatment conditions, EV doses, and isolation methods. To additionally account for inter-donor variability and the hierarchical structure of the experimental design, the data were analyzed using a mixed-effects model with restricted maximum likelihood (REML), specifying treatment as a fixed effect and donor as a random effect, with independent experimental replicates nested within donor. The donor-resolved data and corresponding mixed-effects analysis are provided in the Supplementary Information [Figure 1S].

Normality was evaluated using the Shapiro-Wilk test. Data are presented as mean ± standard deviation (SD), and p < 0.05 was considered statistically significant. All statistical analyses were performed using GraphPad Prism version 11.1.0 (GraphPad Software, USA).

## 3. Results and Discussion

### 3.1 Av-EVs Isolation and Characterization

Extracellular vesicles from *Aloe vera* gel (Av-EVs) were isolated using two protocols: a gentle manual homogenization approach (NB Av-EVs) and a shear force-based method using a commercial blender (B Av-EVs), as illustrated in **Figure 1A**. All vesicle preparations were characterized in accordance with the MISEV2023 guidelines established by the International Society for Extracellular Vesicles (ISEV).^[44]^ Characterization was conducted on four independent replicates for each method.

**Figure 1.**
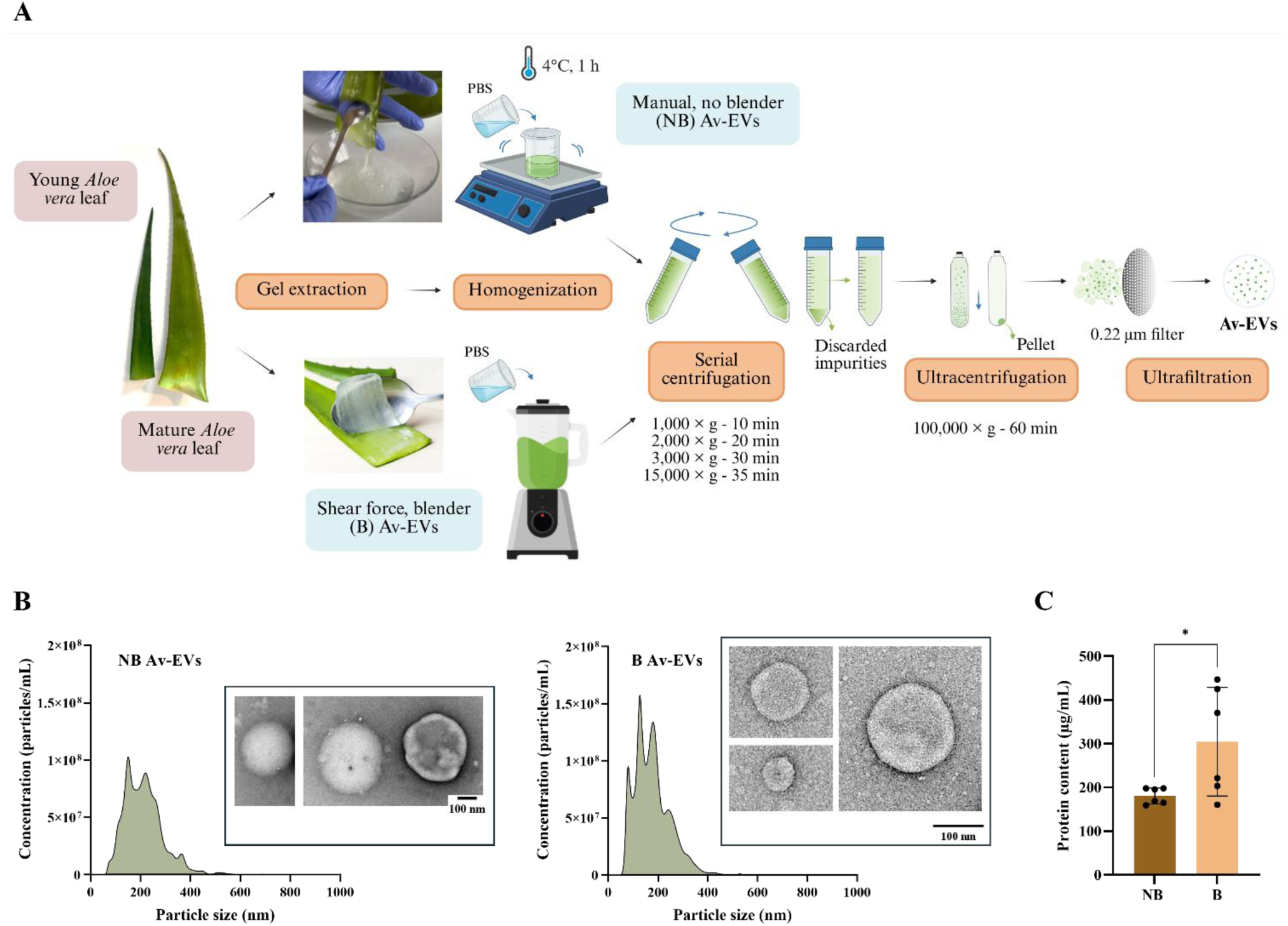
**(A)** Schematic illustration of NB and B Av-EV isolation procedures from Aloe vera barbadensis miller leaves. **(B)** NTA data showing size distribution of typical NB and B Av-EV samples with their respective TEM images. **(C)** Protein content data represents the mean ± SD with n=6 and p < 0.05 *.

Nanoparticle tracking analysis (NTA) established that all final suspension samples consistently yielded concentrations ranging from 1.4 to 2.0 × 10¹⁰ particles/mL, for an average total of 5.6 to 8 × 10^12^ particles per leaf. Furthermore, they displayed heterogeneous size distributions with mean diameters between 150 – 250 nm [**Figure 1B**], consistent with previous reports on *Aloe*-derived EVs.^[35–37, 45]^ While this overall range was reproducible across both methods, NB EVs typically clustered between 1.4 to 1.6 × 10¹⁰ particles/mL, whereas B-derived EVs approached 1.8 to 2.0 × 10¹⁰ particles/mL. The disparity may reflect the influence of shear stress introduced by blender homogenization, which could not only liberate vesicles already present in the inner parenchyma but also mobilize vesicles from within vacuoles or induce vesiculation through mechanical disruption. This hypothesis is supported by literature describing how laminar or turbulent shear flow can destabilize lipid membranes, promoting membrane budding or fragmentation. Such mechanical stress can cause intracellular content leakage and drive the spontaneous formation of EV-like vesicles, often diverse in size and composition.^[46]^

Transmission electron microscopy (TEM) verified the presence of rounded, membrane-bound vesicles with well-defined lipid bilayers – characteristic of EVs. The morphology observed in both NB and B Av-EVs groups supports the preservation of vesicle integrity following isolation, regardless of the extraction method used. Nonetheless, notable differences in size distribution were evident as expected: B-derived Av-EVs displayed a broader range of diameters (approximately 70 to 170 nm), whereas NB-derived EVs showed a more uniform morphology, with most imaged vesicles falling near the upper size range observed by NTA (∼250 nm). While some vesicles measured closer to 300 nm in TEM, this discrepancy may be attributed to swelling or flattening during sample preparation and imaging, a known artifact in conventional TEM workflows. These observations align with NTA results and further support the idea that blender-induced shear stress promotes vesicle heterogeneity, potentially generating a mixture of natural and mechanically derived vesicles.

Protein quantification results provide further evidence of the impact of extraction methods [**Figure 1C**]. NB-derived EVs exhibited consistent protein concentrations across independently processed leaves, ranging from 159 to 198 µg/mL, corresponding to a total protein yield of 3,816 to 4,752 µg per leaf. In contrast, B-derived EVs showed substantial inter-replicate variability, with concentrations spanning 160 to 447 µg/mL and a total protein yield of 3,840 to 10,728 µg per leaf. Statistical analysis confirmed a significant difference in total protein levels between the NB and B groups (p < 0.05), suggesting that the shear-based method may produce a more heterogeneous population of vesicles, potentially including non-physiological particles or protein-rich artifacts. However, when protein levels were normalized to the total EV yield per leaf, both methods exhibited relatively consistent protein-to-particle ratio. NB-derived EVs ranged from 6.81 to 7.43 × 10⁻⁹ µg/EV, while B-derived EVs ranged from 5.33 to 13.41 × 10⁻⁹ µg/EV. Despite the higher variability observed in the raw protein concentration of B samples, the differences between the two groups became less pronounced after normalization, indicating that the protein content per EV remains relatively similar across both EV types.

In summary, both isolation protocols were successful in yielding Av-EVs that meet established EV characterization criteria in terms of size, concentration, and morphology. However, the manual method yielded more uniform vesicle populations with slightly lower protein variability, suggesting greater preservation of native EV characteristics and reduced likelihood of shear-induced artifacts.

### Confirmation of Plant-specific Markers in Av-EVs

PDEVs, similar to mammalian EVs, carry diverse biomolecular cargo capable of modulating recipient-cell function. However, PDEV composition remains comparatively less characterized, and unlike mammalian EVs, a universally accepted set of protein markers has not yet been established across plant species and vesicle subpopulations. Moreover, the vesicle composition and therapeutic performance appear to vary with the physiological and environmental conditions under which vesicles are synthesized and released, as later outlined in section 3.3.3. Nevertheless, several protein families have been recurrently identified in plant EV preparations, including annexins, heat-shock proteins (HSPs), aquaporins, proteins involved in cell-wall remodeling, actins, patellins, syntaxins, clathrin heavy chain, and RAS-related proteins, as well as lipids such as phosphatidic acid (PA), phosphatidylethanolamine (PE), phosphatidylcholine (PC), and plant sphingolipids (GIPCs). Finally, micro RNAs (miRNAs) and metabolites can also be informative.^[47, 48]^

Consistent with this framework, label-free LC–MS/MS analysis of Av-EVs identified multiple proteins previously associated with plant EVs, including HSPs, actins, clathrin heavy chain, small GTPases, syntaxins, annexins, and proteins involved in cell-wall remodeling. These findings provide molecular evidence supporting the plant’s vesicular nature of the isolated particles. A detailed characterization of the Av-EV proteomic profile and maturity-dependent differences in protein composition is presented in section 3.3.4.

To complement the proteomic characterization, immunoblotting was performed for the plant EV-associated proteins TET8 and SYP121/PEN1,[49] together with calnexin as a non-EV/endoplasmic reticulum-associated marker [**Figure 2**]. A SYP121/PEN1-associated signal was detected predominantly within the 35–40 kDa molecular-weight region, consistent with its predicted molecular weight of ∼37.9 kDa. In contrast, no discrete calnexin signal was observed at its expected molecular weight (∼65 kDa), consistent with limited detectable contamination from endoplasmic reticulum (ER)-derived material in the Av-EV preparation. TET8 was not detected under the experimental conditions evaluated.

**Figure 2.**
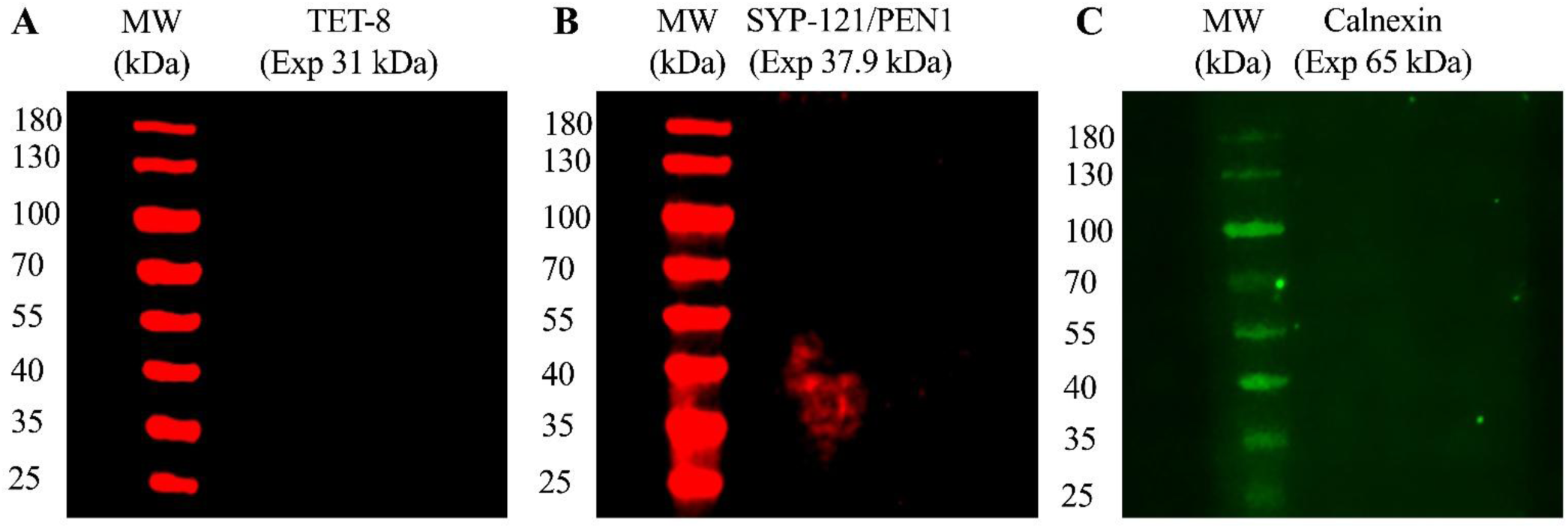
Immunoblot analysis of plant EV-associated and non-EV markers in NB Av-EV preparations (17 µg total protein per lane). EV-associated markers: **(A)** TET8 and **(B)** SYP121/PEN1. Non-EV marker: **(C)** Calnexin. Expected molecular weights were approximately 31, 37.9, and 65 kDa, respectively.

The absence of detectable TET8 likely reflects the heterogeneity of PDEV populations and the condition-dependent abundance of individual markers across plant species, tissue types, physiological states, and vesicle subpopulations. TET8 has been characterized primarily in *Arabidopsis* preparations, particularly in the context of stress and immune responses,^[49–51]^ whereas its expression and enrichment in EVs derived from *Aloe vera* gel remain poorly defined. In this study, Av-EVs identity was therefore confirmed through a complementary, integrated characterization strategy rather than reliance on a single marker. The detection of SYP121/PEN1, the absence of a discrete calnexin signal, and the LC–MS/MS identification of multiple proteins recurrently associated with plant EVs together provide convergent molecular evidence supporting the successful isolation of a plant EV-enriched preparation from *Aloe vera* gel. Collectively, these findings establish a reliable molecular framework for Av-EV characterization and provide a foundation for future studies aimed at defining tissue-specific marker profiles, EV subpopulations, and standardized characterization criteria for *Aloe vera* gel-derived vesicles.

### 3.2 Biological Effects of Av-EVs Cargo from Mature *Aloe vera* Leaves

#### 3.2.1 Antioxidant Capacity

The antioxidant potential of Av-EVs was evaluated using the DPPH radical scavenging assay, a widely accepted method that quantifies a compound’s ability to neutralize free radicals via electron or hydrogen atom donation. In this assay, decreased DPPH absorbance correlates with increased antioxidant activity, expressed as the DPPH inhibition ratio (%). Both NB and B Av-EVs were tested at four concentrations (5×10⁸, 1×10⁹, 5×10⁹, and 2×10¹⁰ EVs/mL) and assessed at four time points (1, 24, 48, and 72 hours) to evaluate both dose-and time-dependent antioxidant dynamics.

As shown in **Figure 3**, all Av-EV treatment groups demonstrated a clear time-dependent increase in DPPH inhibition, indicative of progressive and sustained antioxidant activity. During the first hour, all samples – regardless of dose or method – exhibited relatively uniform inhibition values (∼20 – 40%), suggesting similar initial redox interactions. By 24 hours, B-derived EVs maintained modest inhibition levels (20 – 40%), whereas some NB-derived EV replicates began to display more pronounced scavenging effects, reaching inhibition values of up to ∼80%. Notably, greater variability emerged across replicates between the 24 and 48 h window, potentially reflecting differences in cargo release kinetics, vesicle stability, or vesicle–radical interaction. Despite this fluctuation, the antioxidant response became more consistent by 72 hours, with nearly all treatment conditions – especially those involving NB EVs – stabilizing at high inhibition levels (> 90%).

**Figure 3.**
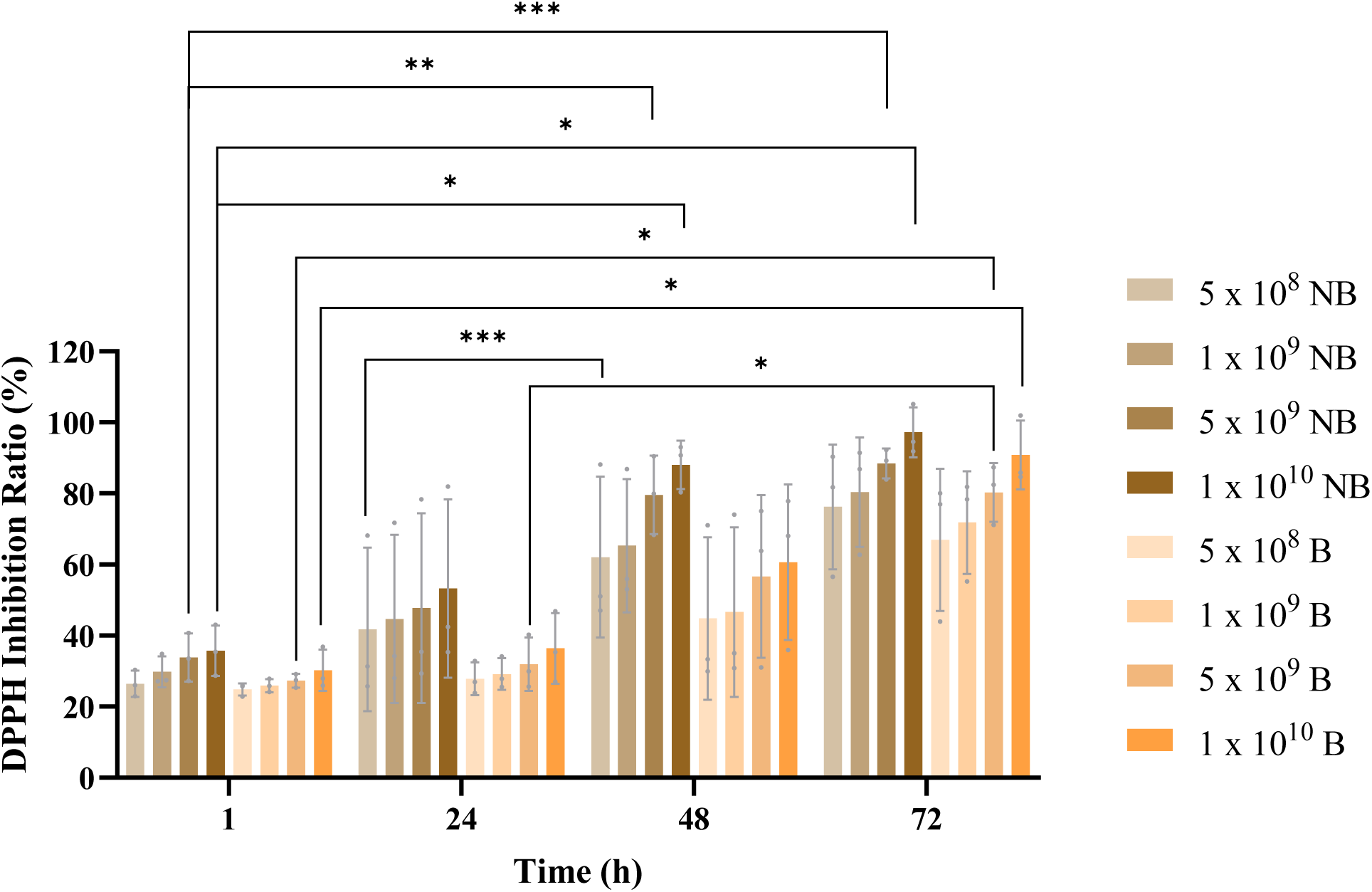
DPPH inhibition ratio (%) over time for NB and B Av-EVs at four concentrations (5×10⁸, 1×10⁹, 5×10⁹, and 2×10¹⁰ EVs/mL). Inhibition data represents the mean ± SD with n=3 and p < 0.05 *, < 0.01 **, < 0.001 ***.

Statistical analysis confirmed a significant increase in inhibition from 1 to 48 hours for NB EVs at both 5×10⁹ and 2×10¹⁰ EVs/mL (p < 0.01 and p < 0.05, respectively), and from 1 to 72 hours (p < 0.001 and p < 0.05, respectively). In contrast, B-derived EVs at the same concentrations reached significance only between 1 and 72 hours (p < 0.05). Furthermore, even the lowest NB EV dose (5×10⁸ EVs/mL) showed a significant increase between 24 and 48 hours (p < 0.001). These findings underscore the sustained redox-modulatory capacity of Av-EVs and suggest that NB-derived vesicles may deliver more timely and effective antioxidant action.

Although statistical significance was not observed across all comparisons, a consistent dose-dependent trend was observed: the highest concentration (2×10¹⁰ EVs/mL) yielded the strongest DPPH inhibition at each time point, whereas the lowest concentration (5×10⁸ EVs/mL) produced the weakest response. This pattern reinforces the premise that the antioxidant efficacy of Av-EVs is proportional to the concentration of vesicle-delivered bioactive compounds. Interestingly, across all tested doses, NB-derived EVs achieved greater inhibition ratios than their B-derived counterparts. This superior performance, even at lower concentrations, highlights the importance of gentle manual extraction methods in preserving vesicle integrity and maximizing functional antioxidant potential.

This antioxidant efficacy aligns with the well-documented phytochemical profile of *Aloe vera*, which includes a variety of redox-active molecules such as α-tocopherol (vitamin E), ascorbic acid (vitamin C), carotenoids, polyphenols (flavonoids and tannins), indoles, alkaloids, and anthraquinones (e.g., aloin and *Aloe*-emodin).^[22, 52–54]^ The plant also contains enzymatic antioxidants (e.g., superoxide dismutase (SOD) and glutathione peroxidase (GPx)) that contribute to ROS detoxification.^[22]^ Prior studies show *Aloe* gel scavenges DPPH, ABTS, and NO radicals in a concentration-dependent manner, limits lipid peroxidation, and protects against oxidative damage.^[23]^ It can further modulate oxidative stress pathways, including inhibition of the NOX4/ROS/p38 axis,^[55]^ preventing activation of latent TGF-β1 and the fibrogenic transition of fibroblasts and endothelial cells into α-SMA–positive myofibroblasts. Consistent with these reports, *Aloe*-derived EVs have been shown to delay skin photoaging via Nrf2/ARE activation, with reductions in H₂O₂,^[37]^ senescence-associated secretory phenotypes (SASP) factors, and malondialdehyde (MDA) *in vitro* and *in vivo*.^[45]^

Importantly, our proteomic profiling (section 3.3.4) corroborates the presence of a vesicle-encoded antioxidant program in mature Av-EVs. These vesicles exhibited enrichment of proteins involved in ether-and sphingolipid-metabolism pathways together with higher aldehyde dehydrogenase (NAD⁺) activity, consistent with enhanced detoxification of lipid-peroxidation products such as 4-HNE. Collectively, these proteomic features provide molecular support for the sustained, dose-responsive reduction in free radical levels observed in our antioxidant capacity assay. Moreover, they help explain how the ability of Av-EVs to mitigate oxidative damage likely disrupts the ROS↔TGF-β1 positive-feedback loop, thereby facilitating the resolution of inflammation and preventing pathological scar formation (e.g., hypertrophic scars, keloids).

#### 3.2.2 Cytotoxicity Assessment

Cell viability, defined as the ability of cells to maintain both functional integrity and proliferative potential, was assessed using the alamarBlue assay, which measures metabolic activity through the reduction of resazurin to fluorescent resorufin by intracellular oxidoreductases in metabolically active cells. This assay provides an integrative measure of cell viability by reflecting plasma membrane integrity, glycolytic flux, mitochondrial respiratory chain activity, and biosynthetic capacity. To evaluate the biocompatibility and potential cytotoxicity of NB and B Av-EVs, the alamarBlue assay was applied to both THP-1-derived macrophages and hDFs exposed to inflammatory or fibrotic stimuli, respectively. Untreated cells served as controls for statistical analysis (–LPS/–EVs or –TGF-β1–VitC/–EVs), and Av-EVs were administered at increasing doses (1,000, 2,500, and 5,000 EVs/cell) to assess their effects on cellular metabolism and viability across both the anti-inflammatory and antifibrotic experimental timelines.

THP-1-derived macrophages were activated using LPS, the major outer membrane component of Gram-negative bacteria, with or without Av-EV co-treatment. As shown in **Figure 4**, LPS alone significantly increased macrophage metabolic activity at 24 h, 72 h, and 7 days compared to the untreated control. Moreover, cotreatment with LPS and Av-EVs further enhanced metabolic activity at 24 and 72 hours across all doses and isolation methods, a period during which cells are expected to exhibit full M1 polarization and heightened inflammatory function. These findings confirm that Av-EVs do not exert cytotoxic effects and that macrophages remain metabolically viable under all treatment conditions.

**Figure 4.**
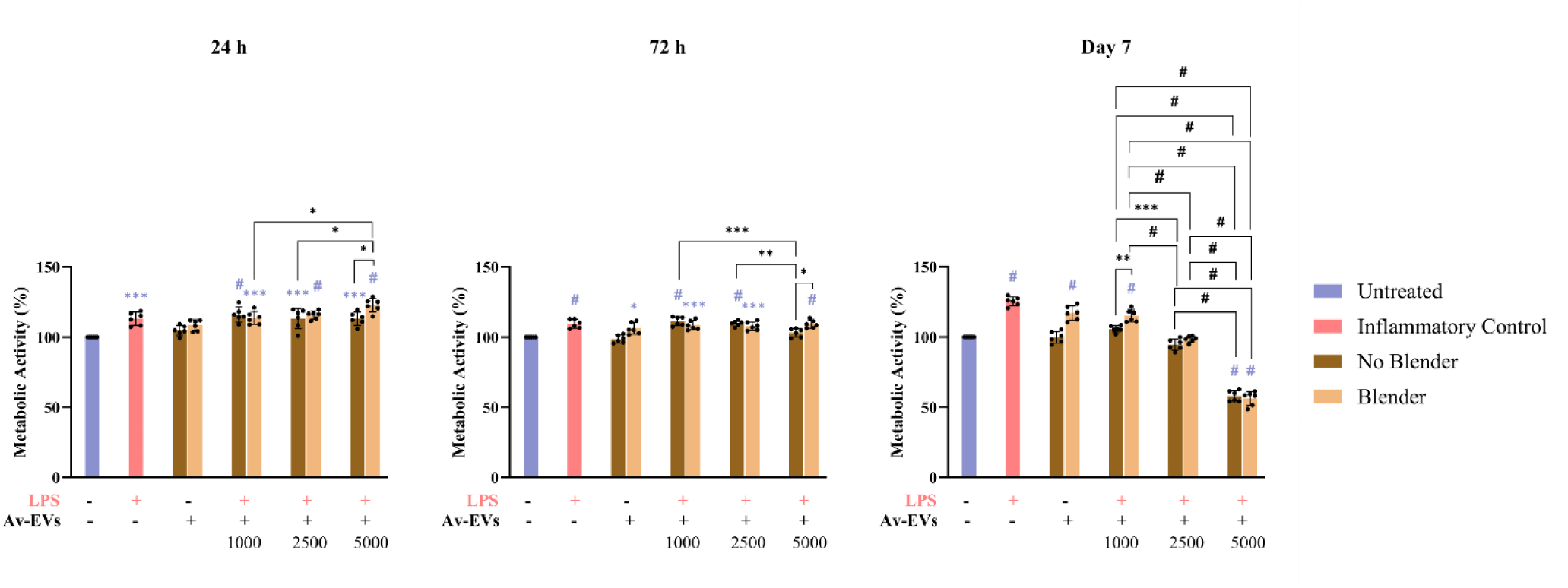
Metabolic activity percentage in LPS-induced macrophages after 24 h, 72 h, and 7 days of co-treatment with No Blender or Blender Av-EVs at increasing doses (1,000, 2,500, 5,000 EVs/cell). Fluorescent data represents the mean ± SD with n=6 and p < 0.05 *, < 0.01 **, < 0.001 ***, < 0.0001 #. Statistical differences relative to untreated control (-LPS/-EVs) are shown in purple, while statistical differences between the isolation methods and doses are shown in black.

The moderate decline in metabolic activity observed between 72 hours and day 7 suggests that Av-EVs facilitate a phenotypic transition from M1 to M2 macrophages, consistent with the expected resolution phase of inflammation as confirmed with the gene expression data later (see section 3.2.4). This phenotypic shift from a pro-inflammatory to a reparative state is not only defined by cytokine expression profiles but also by fundamental metabolic reprogramming. M1 macrophages depend on aerobic glycolysis for rapid ATP production and biosynthesis to support inflammatory signaling, ROS generation, and cytokine secretion.^[56–59]^ In contrast, M2 macrophages rely on oxidative phosphorylation (OXPHOS) and fatty acid oxidation (FAO), to support their anti-inflammatory and tissue-remodeling functions, which are energetically more efficient but generate fewer reduced equivalents.^[56–59]^ Since the alamarBlue assay detects metabolic activity via NADH/NADPH-linked reduction of resazurin,^[60]^ glycolytic M1 cells typically yield stronger signals than OXPHOS-driven M2 cells. Therefore, a decrease in alamarBlue signal at later time points under M2-polarizing conditions, such as Av-EV treatment, likely reflects a metabolic adaptation rather than cytotoxicity. Importantly, the dose-dependent nature of this effect – particularly the notable reduction at 5,000 EVs/cell – suggests that higher concentrations of Av-EVs may more strongly influence macrophage polarization and promote the initiation of tissue repair.

It is also noteworthy that *in vitro*, THP-1-derived macrophages naturally undergo reduced viability over time due to nutrient depletion, lack of stromal survival factors, and activation of intrinsic apoptotic pathways.^[61]^ Similarly, *in vivo* M1 macrophages are generally short-lived, undergoing apoptosis or efferocytosis during the resolution of inflammation, while M2-polarized macrophages tend to persist longer due to their reparative and homeostatic roles.^[62]^

To evaluate the metabolic impact and biocompatibility of Av-EVs on hDFs under fibrotic stimulation, the alamarBlue assay was performed following a 72-hour exposure to TGF-β1 and vitamin C, with or without Av-EV co-treatment. **Figure 5** shows that all groups exposed – including the fibrotic control and the Av- EV co-treated conditions – had a significant increase in metabolic activity compared to the untreated control. This increase reflects the transition from a quiescent fibroblast state to an activated phenotype. In this regard, resting hDFs exhibit low metabolic demand, relying mainly on basal mitochondrial respiration to support ECM maintenance and structural stability. Upon stimulation, activated fibroblasts display increased proliferation, migration, and biosynthesis – particularly of collagen and fibronectin – accompanied by enhanced glycolysis and mitochondrial oxidative metabolism.^[63, 64]^ These shifts led to a measurable rise in cellular redox activity, as detected by the assay.

**Figure 5.**
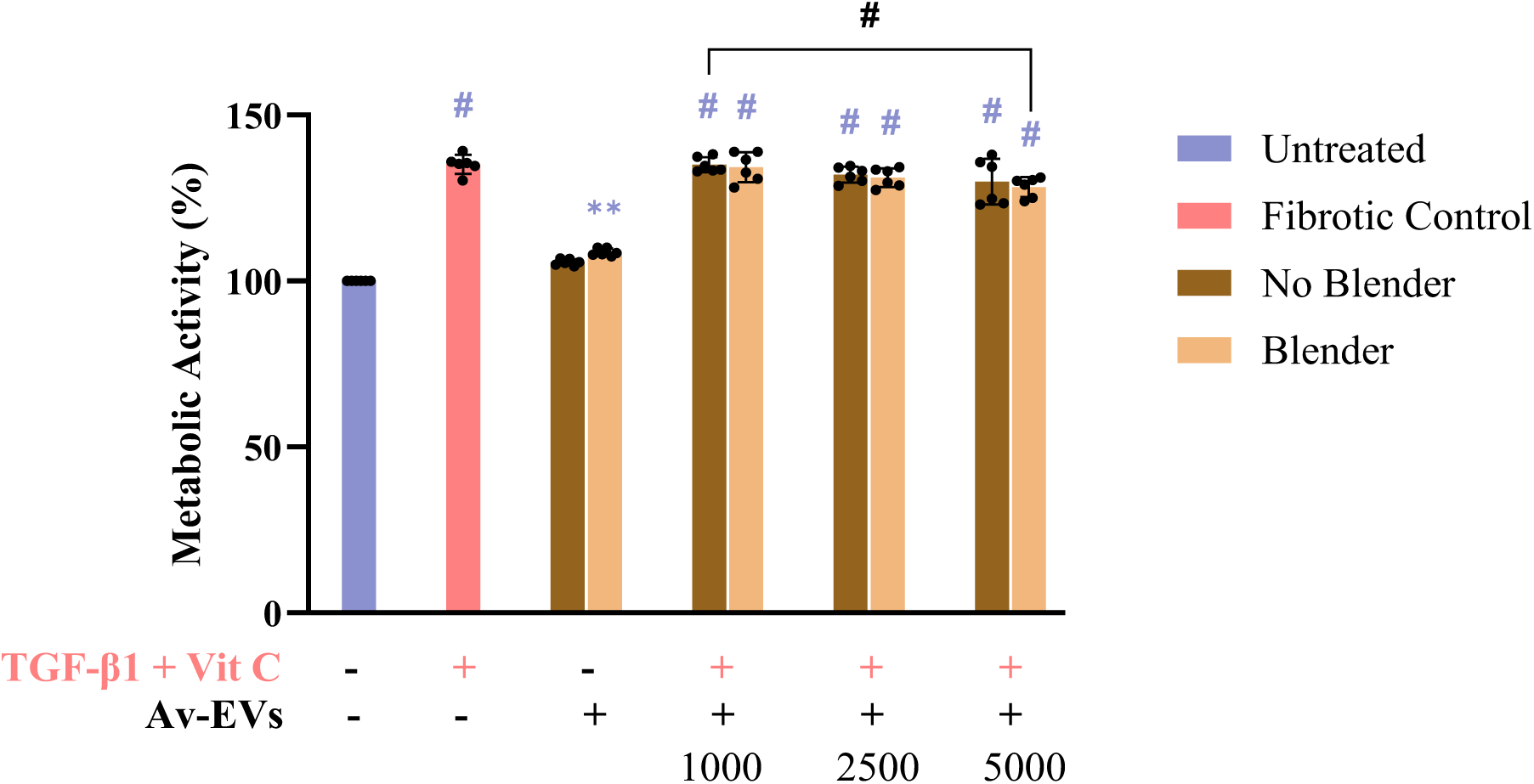
Metabolic activity percentage in TGF-β1 + vitamin C-activated hDFs after 72 h co-treatment with No Blender or Blender Av-EVs at increasing doses (1,000, 2,500, 5,000 EVs/cell). Fluorescent data represents the mean ± SD with n=6 and p < 0.01 **, < 0.0001 #. Statistical differences relative to the untreated control (-TGFꞵ1-VitC/-EVs) are shown in purple, while statistical differences between the isolation methods and doses are shown in black.

However, gene expression analysis and immunofluorescence later revealed that, despite similar metabolic activity across all conditions exposed to TGF-β1 and vitamin C, Av-EV co-treatment significantly suppressed the expression of key fibrotic markers such as COL1A1 and α-SMA, indicating a functional uncoupling between metabolic activity and fibrotic gene expression. This distinction is critical, as elevated metabolic activity alone does not necessarily equate to pathological transdifferentiation into myofibroblasts. Rather, Av-EVs may support a metabolically active yet functionally restrained fibroblast phenotype, in which the cells retain viability and activity but are prevented from progressing into a fully pro-fibrotic myofibroblast state, unlike those in the +TGFꞵ1+VitC/–EVs group.

Collectively, these findings confirm that Av-EVs are non-cytotoxic to both macrophages and hDFs across multiple time points and concentrations, validating their biocompatibility for immunomodulatory applications and supporting their potential use in prolonged treatment regimens aimed at modulating inflammation and preventing fibrosis.

#### 3.2.3 Cellular Uptake of Av-EVs

To confirm cellular uptake, THP-1 and hDFs cells were incubated with the highest dose (D3: 5,000 EVs/cell) of PKH26-labeled Av-EVs and subsequently analyzed by fluorescence microscopy. The results clearly demonstrate that both macrophages and dermal fibroblasts successfully internalized Av-EVs within 24 hours without compromising cell morphology, cytoskeletal organization, or viability, even under inflammatory or fibrotic activation.

**Figure 6A** presents the morphological features of macrophages across different conditions. M0 macrophages displayed a rounded shape, a diffuse cytoplasmic actin network, and clustered nuclei, consistent with a resting, non-polarized phenotype. F-actin was sparsely distributed and mainly localized at the cell periphery, indicating minimal cellular spreading. In contrast, LPS-stimulated macrophages (M1) showed a marked morphological shift, adopting an elongated, stellate shape with prominent stress fibers and lamellipodia, characteristic of M1 polarization. These cytoskeletal changes reflect enhanced motility and phagocytic readiness, driven by pro-inflammatory signaling. Macrophages treated with PKH26-labeled Av-EVs maintained this activated morphology but exhibited distinct red fluorescent signals localized within the perinuclear and cytoplasmic regions, confirming efficient uptake of the vesicles. This internalization, along with preserved cell structure and consistent metabolic activity from viability assays, indicates successful delivery of functional bioactive cargo capable of modulating the expression of both pro-and anti-inflammatory cytokines, as later confirmed in the anti-inflammatory response experiment (see section 3.2.4).

**Figure 6.**
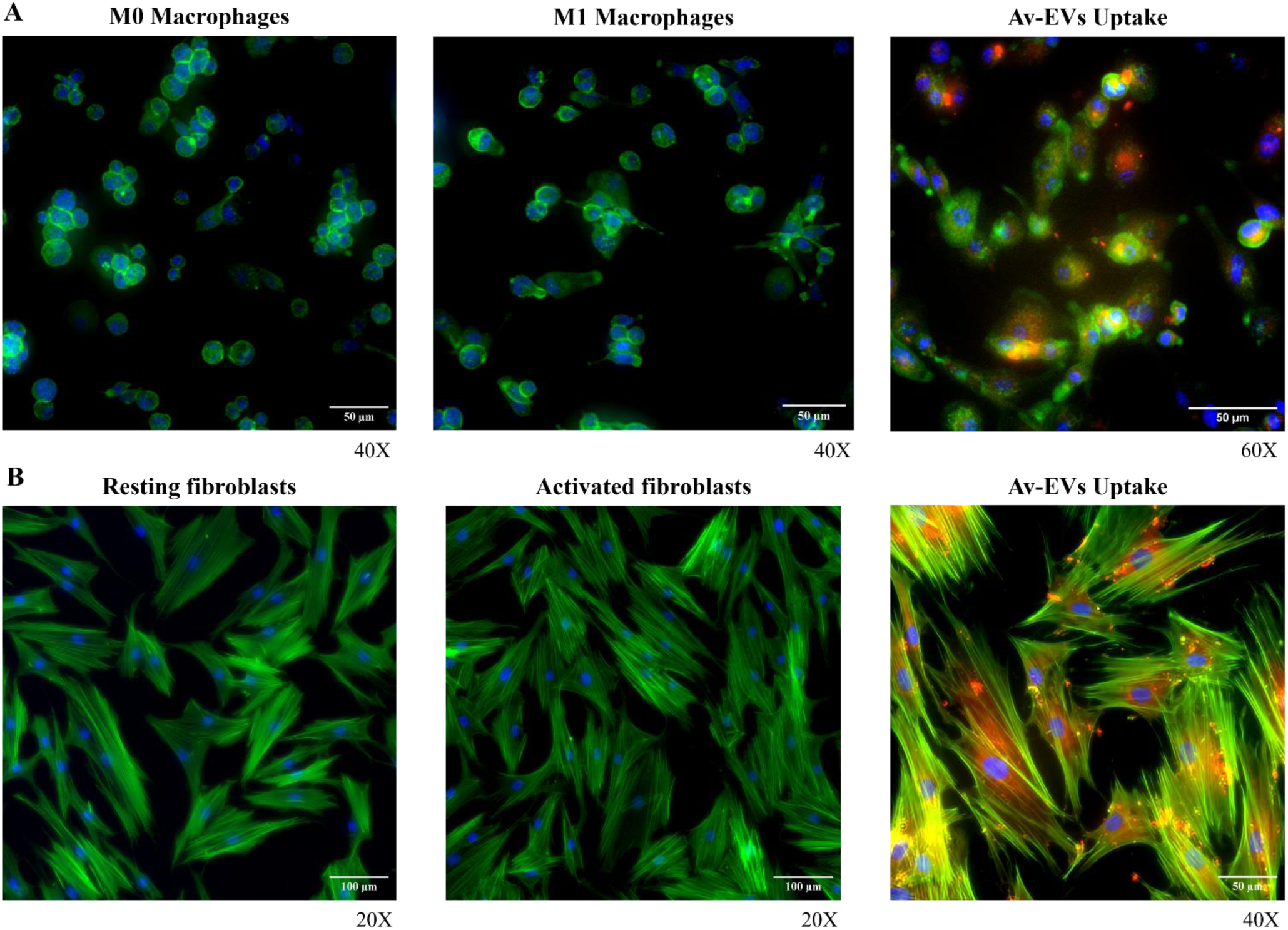
(A) From left to right: M0 macrophages, activation of M0 macrophages with LPS to induce M1 polarization, and cellular uptake of Av-EVs into M1 macrophages. (B) From left to right: resting hDFs, activation of hDFs with TGF-β1 and vitamin C, and cellular uptake of AS-EVs into activated hDFs. PKH26 red fluorescent dye-labeled Av-EVs (red), F-actin (green), and nuclei (blue).

**Figure 6B** shows the progression of fibroblast morphology following activation and Av-EV treatment. Resting fibroblasts exhibited the expected spindle-shaped appearance, with elongated nuclei and well-organized, parallel actin filaments aligned along the cell’s longitudinal axis. These features are typical of quiescent fibroblasts that contribute to ECM maintenance under physiological conditions. After treatment with TGF-β1 and vitamin C, cells adopted a broader, more spread-out shape, with increased density and alignment and intensified F-actin staining, indicating the early stages of fibroblast-to-myofibroblast transdifferentiation. In cells exposed to PKH26-labeled Av-EVs, this activated morphology and cytoskeletal structure was retained, further confirming the non-cytotoxic and non-disruptive nature of Av-EV internalization. Importantly, bright red fluorescent signals were clearly visible, particularly in the perinuclear zones, indicating efficient vesicle uptake and intracellular delivery cargo capable of modulating gene expression of key fibrotic markers such as COL1A1 and α-SMA as later confirmed in the antifibrotic response experiment (see section 3.2.5).

#### 3.2.4 Anti-inflammatory Response (Macrophages)

Two isolation protocols were compared to assess whether mechanical homogenization of *Aloe vera* gel alters the biological efficacy of the resulting vesicles. Both NB and B EVs were purified using identical centrifugation and ultracentrifugation workflows, ensuring comparability. THP-1 monocytes were differentiated into macrophages and further induced with LPS to adopt a proinflammatory M1 polarity, while being exposed to cotreatment with Av-EVs. The experimental conditions were as stated in **Table 1**.

##### Suppression of Pro-inflammatory Cytokines Expression at 24 Hours

RT-qPCR analysis of LPS-induced THP-1-derived macrophages revealed a robust upregulation of pro-inflammatory cytokines, including CXCL10/IP-10, CCL2/MCP-1, IL-1β, TNF-α, and IL-6 compared to untreated controls, consistent with classical M1 macrophage polarization [**Figure 7**]. In contrast, cotreatment with Av-EVs significantly attenuated this pro-inflammatory profile within 24 h, in a dose-and preparation-dependent manner. Notably, Av-EV control groups did not induce cytokine expression, with levels remaining near baseline, confirming that Av-EVs do not activate macrophages under homeostatic conditions.

**Figure 7.**
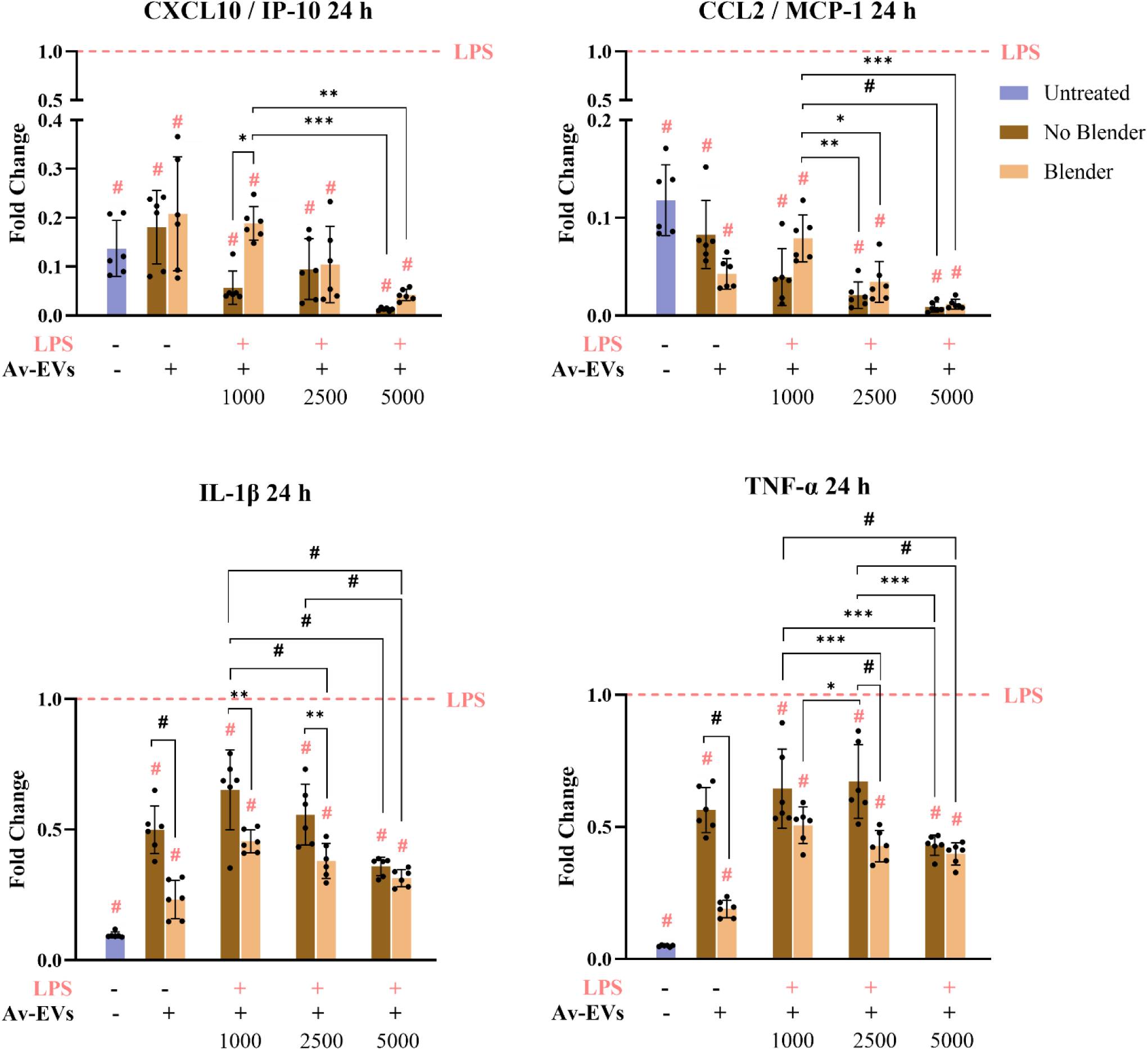
CXCL10/IP-10, CCL2/MCP-1, IL-1β, TNF-α gene expression in LPS-induced macrophages after 24 h of co-treatment with No Blender or Blender Av-EVs at increasing doses (1,000, 2,500, 5,000 EVs/cell). RT-qPCR data represents the mean ± SD with n=6 and p < 0.05 *, < 0.01 **, < 0.001 ***, < 0.0001 #. Statistical differences relative to the inflammatory control (+LPS/-EVs) are shown in pink, while statistical differences between the isolation methods and doses are shown in black.

Among the suppressed genes, CXCL10/IP-10 – a chemokine induced by IFN-γ and LPS that promotes Th1 responses and recruits activated T cells and macrophages – was significantly downregulated. Similarly, CCL2/MCP-1, a critical chemokine for monocyte recruitment and a well-established marker of M1-type polarization, was also strongly inhibited. Both NB and B Av-EVs significantly reduced the expression of CXCL10 and CCL2 in a dose-responsive manner, with near-complete suppression observed at the highest dose (D3, 5000 EVs/cell). Specifically, CXCL10 expression was reduced to a fold change of ≤ 0.017 (NB) and 0.059 (B), while CCL2 was suppressed to ≤ 0.017 (NB) and 0.021 (B). Even at the lowest dose (D1, 1000 EVs/cell), strong inhibitory effects were observed, with CXCL10 expression reduced to ≤ 0.126 (NB) and 0.248 (B), and CCL2 reduced to ≤ 0.094 (NB) and 0.118 (B), where “≤” indicates that the reported value represents the highest observed level within that condition. These data highlight the potent gene-modulatory capacity of both EV preparations, even at lower concentrations. Notably, NB-derived Av-EVs consistently outperformed the B-derived Av-EVs across all doses for both gene targets, with a statistically significant difference at the lowest dose for CXCL10 expression.

In addition, the pro-inflammatory cytokines IL-1β and TNF-α were also evaluated. IL-1β is a canonical early-response cytokine, tightly linked to inflammasome activation, and is a major driver of tissue inflammation and fibrosis. TNF-α, a central cytokine in M1-driven inflammation, initiates and sustains pro-inflammatory cascades in response to endotoxins like LPS. As expected, both IL-1β and TNF-α were significantly upregulated in LPS-stimulated macrophages. Upon cotreatment with Av-EVs, both cytokines were strongly downregulated across all doses, with D3 (5000 EVs/cell) achieving the maximum suppression, regardless of the extraction method. Previous research in *Aloe vera*-EVs also demonstrated this dose-dependent effect.^[35, 36]^ Interestingly, at lower and intermediate doses (D1 and D2), B Av-EVs exhibited superior suppressive effects compared to NB, with D2 (2500 EVs/cell) demonstrating comparable efficacy to D3 in both preparations.

The observed suppression of these cytokines following Av-EV treatment demonstrates that Av-EVs act rapidly and effectively to modulate macrophage inflammatory responses. Upon LPS stimulation, Toll-like receptor 4 (TLR4) activation triggers the MyD88-dependent signaling cascade, leading to phosphorylation and degradation of IκBα, which permits nuclear translocation of NF-κB p65/p50 dimers and subsequent transcription of a wide array of pro-inflammatory genes, including CXCL10, CCL2, IL-1β, TNF-α, and IL-6. The downregulation of IL-1β and TNF-α, in particular, implies that Av-EVs could interfere with canonical TLR4/NF-κB signaling, potentially by inhibiting upstream kinases such as IKK, preserving IκBα stability, and ultimately preventing NF-κB nuclear localization.^[65]^ This inhibitory effect may be attributed to specific phytochemical and molecular cargos within Av-EVs – such as flavonoids and miRNAs – previously shown to antagonize NF-κB signaling and inflammatory gene expression.^[66–68]^

Consistent with these transcriptional findings, proteomic profiling (section 3.3.4) later confirmed a coordinated decreased abundance of TLR, NOD-like receptor, MAPK, PI3K–Akt, and JAK–STAT signaling components in mature Av-EVs, together with enrichment of proteins that are part of the lipid-remodeling and antioxidant-handling pathways (e.g., ether and sphingolipid metabolism). This proteomic pattern supports the notion that mature Av-EVs can suppress the MyD88/TRAF6→NF-κB/MAPK axis at multiple levels while reinforcing redox balance and membrane-dependent immunoregulation – mechanisms fully aligned with the early reduction in CXCL10, CCL2, IL-1β, TNF-α, and IL-6 observed at 24 hours.

##### Modulation of M1-to-M2 Polarization from 24 Hours to 7 Days

IL-6 is a pleiotropic cytokine with complex roles in both pro-inflammatory signaling and tissue regeneration, making its behavior during macrophage activation highly context-dependent. During the early phase of activation (24 hours), IL-6 was significantly upregulated in LPS-treated macrophages [**Figure 8**]. At this point, Av-EV treatment did not result in a substantial modulation of IL-6 expression, regardless of dose or isolation method. This suggests that IL-6 expression during the acute inflammatory response is initially resistant to Av-EV-mediated suppression. After 72 hours, a notable shift occurred. IL-6 expression was significantly downregulated in Av-EV-treated groups compared to the LPS-only condition, with p-values reaching < 0.001 and < 0.0001 at intermediate and high doses for both NB and B EVs. This delayed suppression aligns with the early resolution phase of inflammation, during which IL-6 transitions from a pro-inflammatory effector to a mediator of immune regulation and repair and confirming that IL-6 elevation was part of a controlled resolution process rather than chronic inflammation.

**Figure 8.**
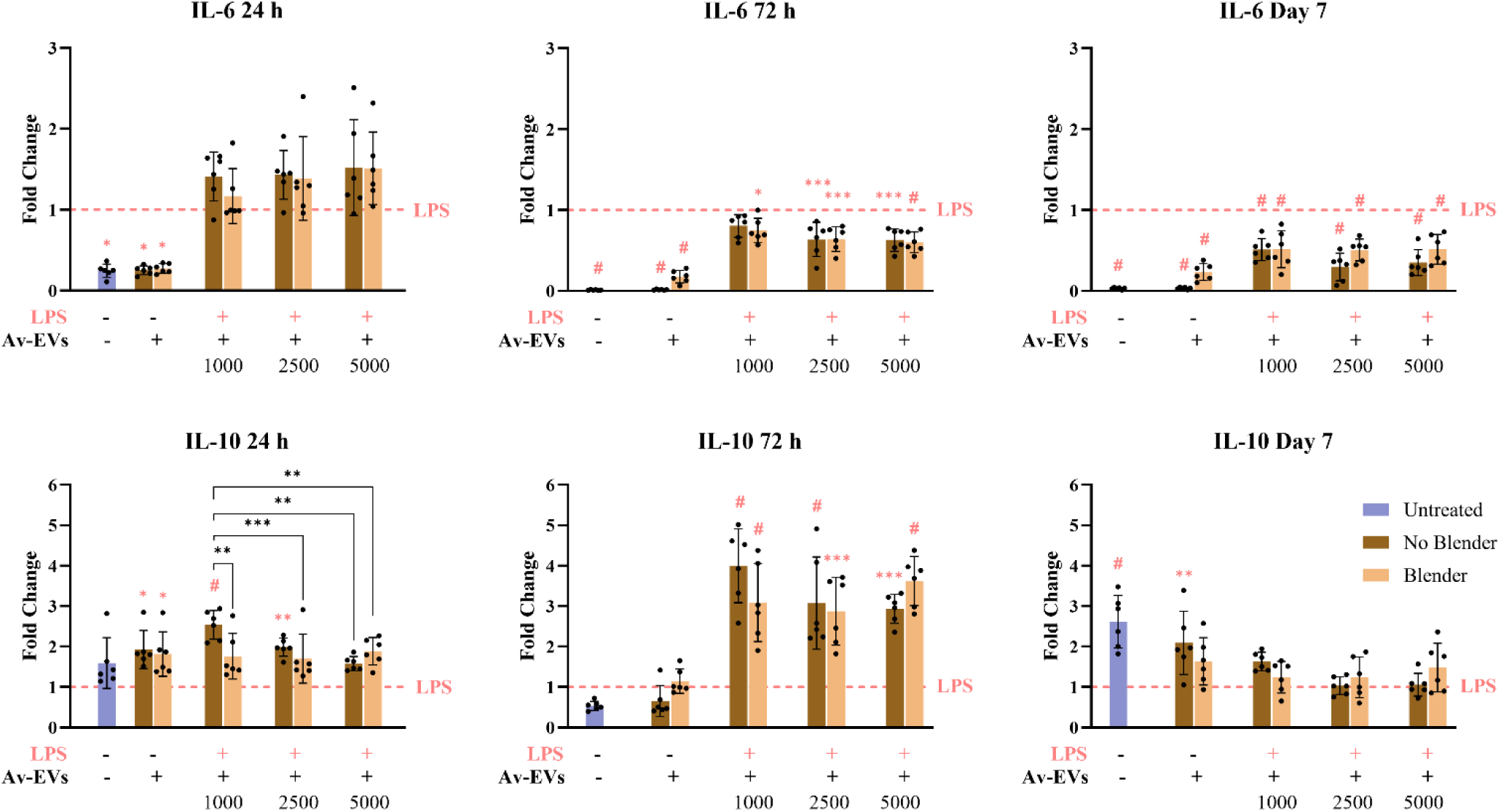
IL-6 and IL-10 gene expression in LPS-induced macrophages after 24 h, 72 h, and 7 days of co-treatment with No Blender or Blender Av-EVs at increasing doses (1,000, 2,500, 5,000 EVs/cell). RT-qPCR data represents the mean ± SD with n=6 and p < 0.05 *, < 0.01 **, < 0.001 ***, < 0.0001 #. Statistical differences relative to the inflammatory control (+LPS/-EVs) are shown in pink, while statistical differences between the isolation methods and doses are shown in black.

Importantly, IL-6 appeared to play a transitional role in initiating macrophage phenotype switching from M1 to M2, a process supported by the concurrent rise in IL-10 expression.^[69–73]^ This pattern is consistent with the activation of STAT3-mediated signaling as described in the literature, where both IL-6 and IL-10 are known to engage the Janus kinase (JAK)/signal transducer and activator of transcription 3 (STAT3) pathway. Upon receptor engagement, these cytokines trigger JAK phosphorylation, which subsequently activates STAT3 through tyrosine phosphorylation. Activated STAT3 then dimerizes and translocates into the nucleus, promoting transcription of anti-inflammatory and reparative genes such as SOCS3, ARG1, and IL-10.^[71, 72]^ In the context of Av-EV treatment, the initial persistence of IL-6, followed by its suppression alongside peak IL-10 expression, suggests that Av-EVs may support a controlled, pro-resolving environment possibly involving STAT3 activity. While the current study did not directly assess STAT3 phosphorylation or nuclear translocation, the observed gene expression kinetics align with previously reported STAT3-dependent resolution mechanisms. By day 7, IL-6 remained consistently suppressed across all Av-EV-treated conditions, regardless of dose or extraction method, indicating the establishment of a stable, anti-inflammatory macrophage state, likely corresponding to M2 polarization and resolution of the inflammatory response.

In parallel, IL-10 expression patterns strongly support this interpretation. At 24 hours, both NB and B Av-EVs induced a dose-dependent upregulation of IL-10, with a particularly strong response at the lowest dose (1000 EVs/cell) – inversely mirroring the IL-6 profile. Notably, NB EVs significantly outperformed B EVs in upregulating IL-10 at early and intermediate time points, suggesting superior early immunomodulatory potency. By 72 hours, IL-10 expression peaked in all EV-treated groups, coinciding with the sharp decline in IL-6 levels and further confirming the M1-to-M2 phenotypic transition. NB EVs again stood out, achieving up to a 5-fold increase in some replicates. By Day 7, IL-10 levels began to decline across all conditions, consistent with the transient nature of IL-10 in initiating the anti-inflammatory cascade and subsequent feedback mechanisms that restore immune homeostasis. This temporal profile underscores IL-10’s role as an early switch that helps reset inflammatory macrophages toward a reparative fate which appeared later in the timeline.^[72]^

##### Transition to an M2 Phenotype and Activation of an Anti-inflammatory Marker in 7 Days

Although IL-10 expression declined by day 7, the combination of (i) sustained suppression of M1-associated cytokines during the critical 72-hour window and (ii) concurrent induction of M2-associated markers, supports the notion of an orchestrated immune transition toward inflammation resolution and tissue repair. These findings suggest that Av-EV treatment facilitates a durable shift toward an anti-inflammatory phenotype, not only by mediating early immunosuppression but also by reinforcing late-stage macrophage reprogramming, essential transition for initiating reparative processes such as wound healing, ECM remodeling, and fibrosis modulation. Aloe-derived vesicles have likewise been reported to modulate macrophage immunity *in vitro* and in clinical bronchoalveolar lavage fluid samples *ex vivo*. The authors assessed macrophage M1-to-M2 reprogramming by flow cytometry of classic surface readouts: CD86 (M1) decreased while CD206 (M2) increased after vesicle uptake, accompanied by IL-10 elevation, shifting the overall M1/M2 balance toward an anti-inflammatory state. Although our study did not quantify CD86 or CD206, these observations outline practical endpoints that support our data and may guide future validation.^[73]^

Consistent with a pro-resolving program, ARG1 – a canonical M2 marker involved in proline synthesis and collagen stabilization –^[74]^ was significantly upregulated in all Av-EV-treated groups compared to the LPS-only condition by day 7 [**Figure 9**]. While variability existed among replicates, several exhibited fold changes between 5 and 10, indicating a robust response. Although differences between EV doses and isolation methods did not reach statistical significance, a clear dose-dependent trend was observed in both types of EVs with a better response in NB Av-EVs with significance even at the lowest dose. The trend aligns with the alamarBlue viability data, where reduced metabolic activity at higher Av-EV concentrations on day 7 may reflect enhanced polarization toward a reparative, quiescent M2 phenotype, rather than cytotoxicity. Additionally, the emergence of ARG1 expression may imply a sequential maturation from M2 into more specialized, tissue-repairing subtypes (e.g., M2a/M2c). However, further research is necessary to elucidate the specific contributions of Av-EV cargo in the modulation of cytokine signaling, receptor engagement, and downstream transcriptional programming involved in M2 macrophage polarization.

**Figure 9.**
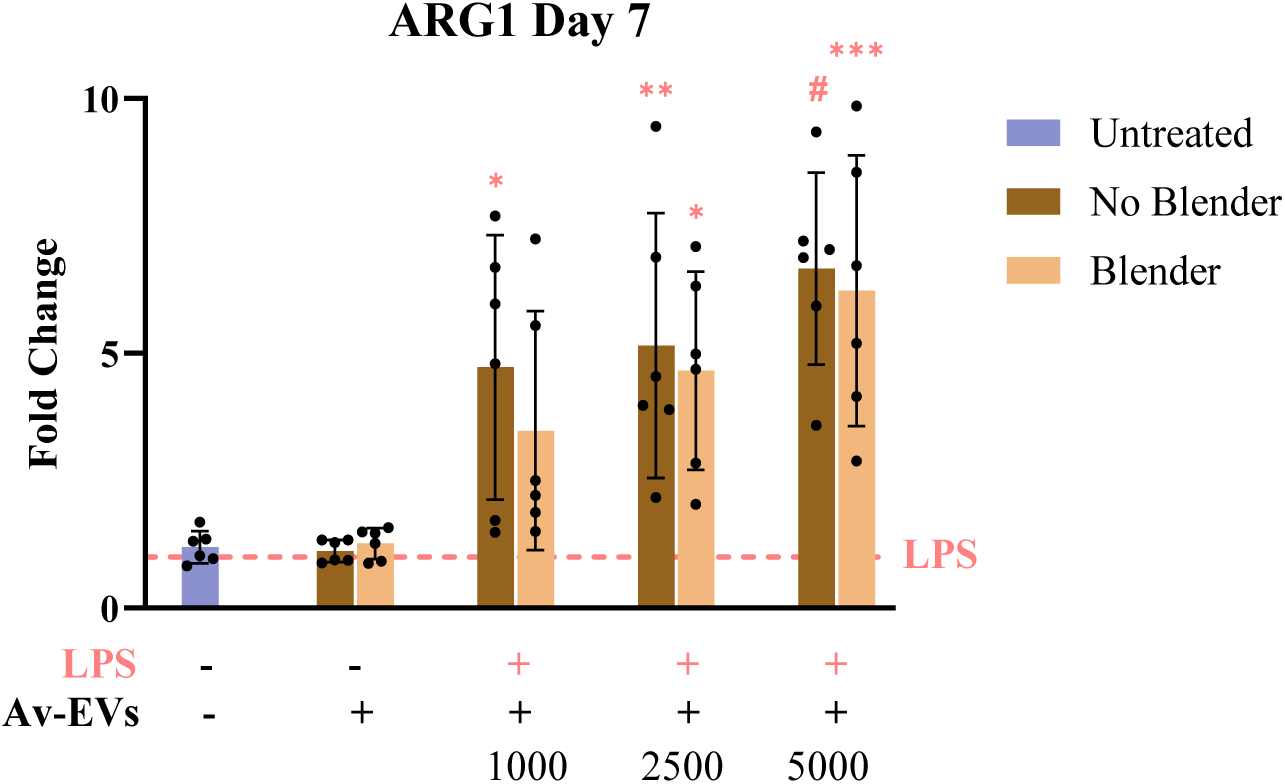
ARG1 gene expression in LPS-induced macrophages after 7 days of co-treatment with No Blender or Blender Av-EVs at increasing doses (1,000, 2,500, 5,000 EVs/cell). RT-qPCR data represents the mean ± SD with n=6 and p < 0.05 *, < 0.01 **, < 0.001 ***, < 0.0001 #. Statistical differences relative to the inflammatory control (+LPS/-EVs) are shown in pink.

It is worth noting that while M2 macrophages are known to facilitate tissue repair, they may contribute to pathological fibrosis through excessive collagen deposition under persistent inflammatory conditions.^[75, 76]^ Favorably, in this study, Av-EV treatment also resulted in significant downregulation of fibrotic markers COL1A1 and α-SMA in fibroblasts (see section 3.2.5), indicating that matrix remodeling occurred without promoting fibrotic tissue accumulation. Moreover, we have already demonstrated that Av-EVs strongly suppressed key pro-inflammatory mediators at earlier time points, suggesting that the inflammatory stimulus had been resolved. In this context, M2 macrophage activity likely operates under homeostatic regulation, supporting tissue regeneration while minimizing the risk of fibrosis. Taken together, these findings highlight the therapeutic potential of NB and B Av-EVs as naturally derived nanocarriers capable of orchestrating both early and late phases of macrophage reprogramming.

#### 3.2.5 Antifibrotic Response (Dermal Fibroblasts)

Myofibroblasts differentiation was induced in hDFs cultures exposed to 10 ng/mL TGF-ꞵ1, a potent pro-fibrotic factor that promotes disorganized and excessive collagen deposition and enhances cell contractility and matrix tension through increased α-SMA stress fiber formation, recapitulating key features of pathological scarring.^[77]^ Vitamin C was included to enhance metabolic activity and collagen synthesis, thereby strengthening the fibrotic phenotype.^[78]^ The study further evaluated the capacity of Av-EVs, isolated either manually (NB) or via shear force homogenization (B), to attenuate this phenotype over a 72-hour period using three dose levels. Fibrotic modulation was assessed by quantifying COL1A1 and α-SMA, two canonical myofibroblast markers. The experimental conditions were as stated in **Table 2**.

**Figure 10** presents fold changes in gene expression for all treatment groups relative to the fibrotic positive control (+TGF-β1+VitC/–EVs), normalized to a fold change of 1. As expected, this condition exhibited a marked upregulation of both genes compared to untreated hDFs and EV-only controls, consistent with a fibrotic phenotype characterized by enhanced ECM deposition and myofibroblast differentiation.

**Figure 10.**
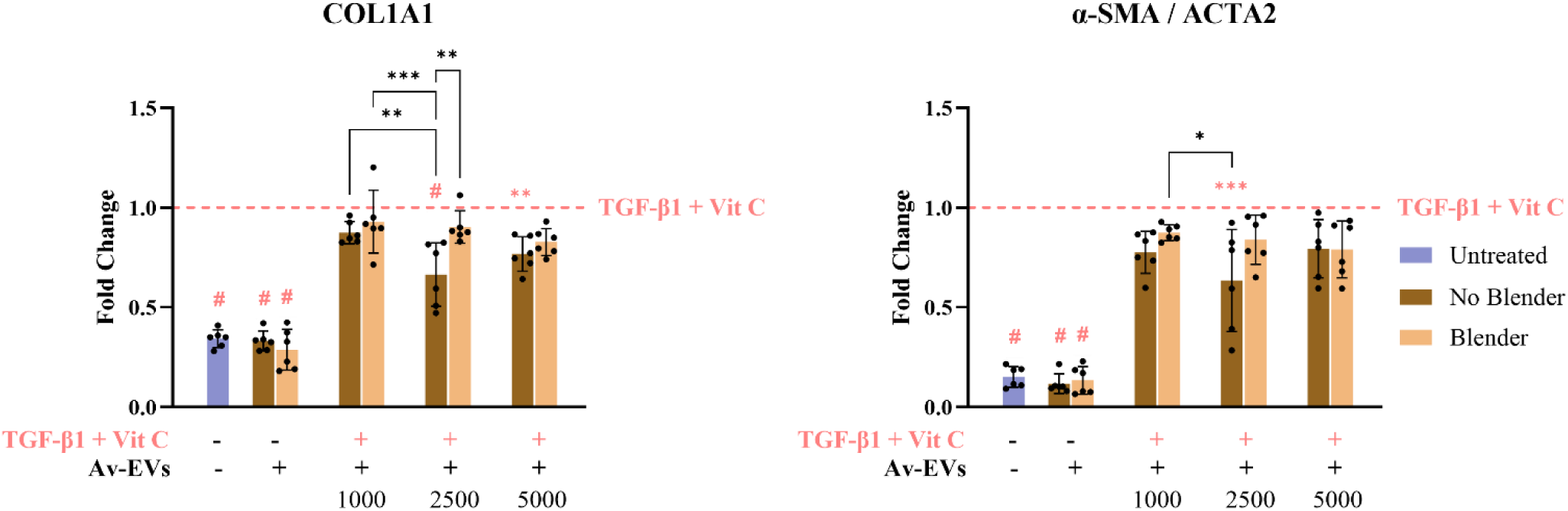
COL1A1 and α-SMA/ACTA2 gene expression in TGF-β1 + vitamin C-activated hDFs after 72 h co-treatment with No Blender or Blender Av-EVs at increasing doses (1,000, 2,500, 5,000 EVs/cell). RT-qPCR data represents the mean ± SD (n=6 measurements; two biological donors with three independent experimental replicates per donor). p < 0.05 *, < 0.01 **, < 0.001 ***, < 0.0001 #. Statistical differences relative to the fibrotic control (+TGFꞵ1+VitC/-EVs) are shown in pink, while statistical differences between the isolation methods and doses are shown in black.

Untreated hDFs displayed low baseline levels of COL1A1 (≤ 0.409) and α-SMA (≤ 0.216), consistent with a non-fibrotic state (p < 0.0001). Similarly, EV-only conditions remained comparable to untreated controls (p < 0.0001), where NB-derived EVs showed consistent levels for both COL1A1 (≤ 0.420) and α-SMA (≤ 0.216), while B-derived EVs exhibited levels of ≤ 0.424 for COL1A1 and ≤ 0.229 for α-SMA. These findings indicate that Av-EVs alone do not promote fibrotic activation in resting fibroblasts and maintain gene expression levels within a physiological range under basal conditions.

Upon fibrotic stimulation, treatment with NB Av-EVs led to a pronounced attenuation of TGF-β1/VitC-induced expression of both markers. This effect was most evident at intermediate (2,500 EVs/cell) and high doses (5,000 EVs/cell) where COL1A1 was significantly reduced (p < 0.0001 and p < 0.01, respectively) and α-SMA expression reached a p < 0.0001 at the intermediate dose. Notably, there was a statistically significant difference between NB and B Av-EVs for COL1A1 at the intermediate dose (p < 0.01), indicating that the manual method of isolation counteracts more effectively TGF-β1-mediated transcriptional activation of collagen genes compared to those EVs isolated via shear force homogenization, ultimately confirming that the isolation method may influence the biological potency of Av-EVs.

To evaluate the robustness of these findings across biological donors, the hDF data were further analyzed using a mixed-effects model (REML). The complete analysis, including donor-specific responses, is provided in the Supplementary Information [Figure 1S]. Although some inter-donor variability was observed, the mixed-effects analysis confirmed the overall conclusions, with the intermediate and highest doses of NB Av-EVs remaining the most effective in reducing fibrotic marker expression.

Overall, these findings demonstrate that Av-EVs have the capacity to limit fibroblast-to-myofibroblast transdifferentiation by the coordinated downregulation of key fibrotic markers such as COL1A1 and α-SMA, leading to limited ECM deposition, and impaired cytoskeletal remodeling and force generation – processes essential for myofibroblast function and fibrotic scar contracture.^[35, 64]^. This coordinated downregulation further suggests that Av-EVs may interfere with the TGF-β1/SMAD signaling axis or its downstream effectors. In the canonical pathway, TGF-β1 binding to its receptor complex (TGF-βRI/II) induces phosphorylation of SMAD2 and SMAD3, which subsequently form a transcriptionally active complex with SMAD4 that translocates to the nucleus to drive expression of pro-fibrotic genes, including COL1A1 and α-SMA (ACTA2). Although direct inhibition of SMAD4 by *Aloe vera* has not been reported, several *Aloe*-derived bioactives – such as acemannan, aloin and flavonoids – have been shown to inhibit SMAD2/3 phosphorylation and nuclear translocation.^[79, 80]^ This may disrupt the formation and activity of the SMAD2/3/4 complex, thereby impairing the transcriptional regulation of downstream signaling.

Consistent with these findings, proteomic enrichment in mature Av-EVs (section 3.3.4) highlighted modules that directly impinge on contractile and matrix programs, including regulation of the actin cytoskeleton, Ca²⁺-handling/channel activities, and lipid-remodeling pathways, all of which are predicted to reduce ERK, p38 MAPK, RhoA/ROCK, and PI3K/Akt/mTOR tone, limit stress-fiber assembly, dampen pro-contractile signaling, and attenuate redox-driven TGF-β amplification.^[21, 65, 81]^ Moreover, proteomic signatures reinforce the presence of non-canonical brakes on TGF-β signaling, evidenced by the relative depletion of translation-associated (ribosome) and proline-supply modules, indicative of decreased collagen biosynthetic/maturation throughput and congruent with the measured reduction in COL1A1 levels.

Taken together, these combined mechanisms support the notion that the bioactive cargo of NB Av-EVs functions as a potent modulator of fibrotic signaling, shifting fibroblasts away from a pro-fibrotic trajectory toward a more quiescent, regenerative phenotype. Furthermore, the dual activity of Av-EVs – immune modulation and fibroblast reprogramming – exert a coordinated regulation of inflammation and fibrosis, two key phases that must be tightly balanced to ensure effective wound healing and tissue regeneration without abnormal scar progression.

#### 3.2.6 Collagen type I and Smooth Muscle Actin Immunofluorescence Analysis

To further validate the antifibrotic activity of Av-EVs at the protein level, collagen I accumulation and α-SMA expression were assessed by immunofluorescence microscopy in hDFs maintained under healthy or TGF-β1/vitamin C-induced fibrotic conditions, with or without Av-EVs co-treatment (2,500 EVs/cell). Collagen I and α-SMA were labeled in green, F-actin in red, and cell nuclei in blue. Fluorescence intensity was quantified from three independent images per condition following background subtraction. As the collagen I and α-SMA analyses were performed using different hDF donors, each staining dataset was interpreted relative to its corresponding untreated and fibrotic controls, allowing treatment-associated changes to be assessed within each immunofluorescence experiment.

As shown in **Figure 11A and C**, untreated hDFs exhibited relatively low collagen I-associated fluorescence, consistent with basal matrix production in resting fibroblasts. In contrast, TGF-β1/vitamin C stimulation produced a marked increase in signal, accompanied by abundant green fluorescence distributed throughout the cells, reflecting enhanced collagen synthesis and matrix accumulation characteristic of the induced fibrotic phenotype. Quantification confirmed an approximately 4.7-fold increase relative to untreated hDFs (p < 0.0001).

**Figure 11.**
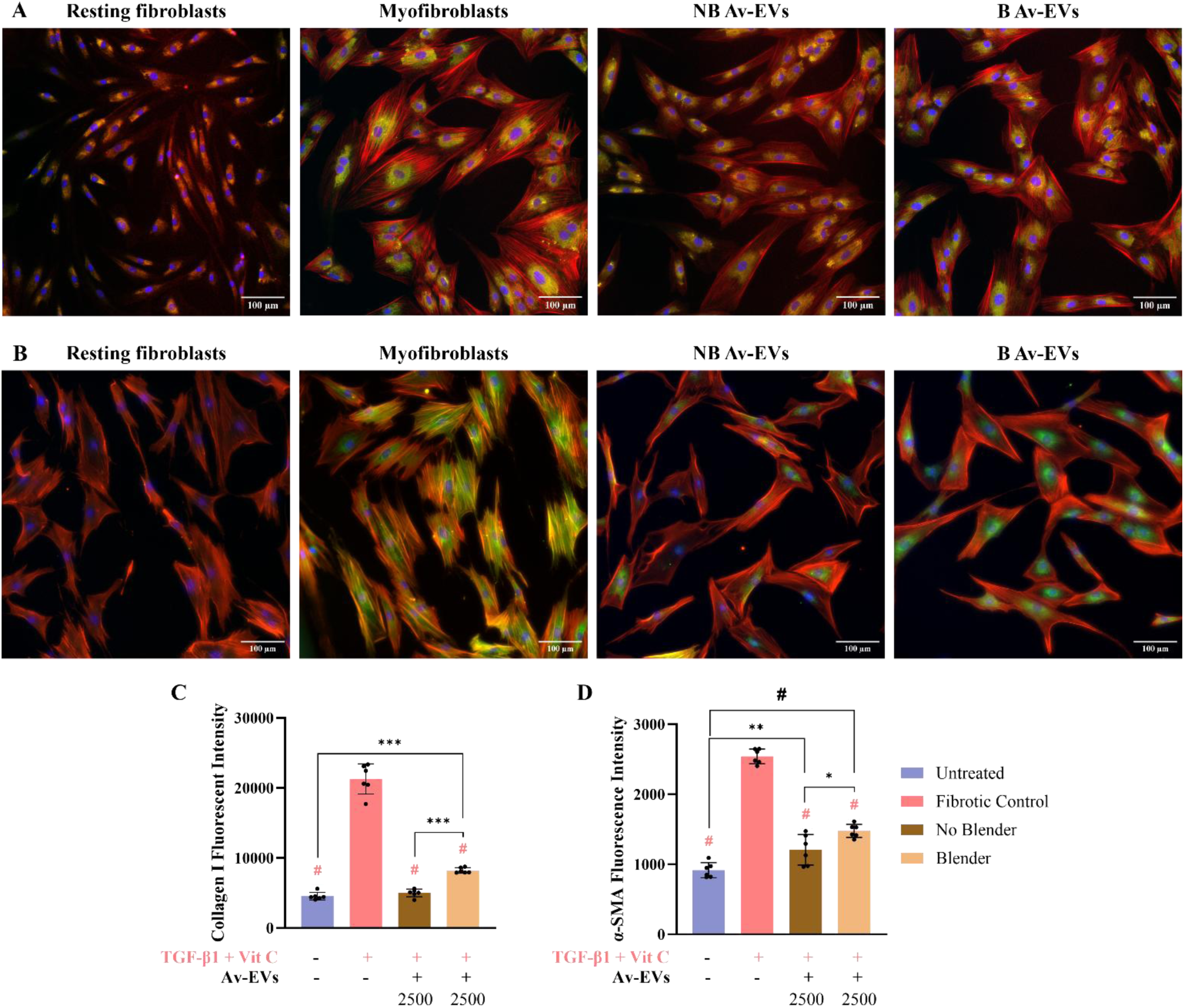
Immunofluorescence analysis of **(A)** collagen I and **(B)** α-SMA expression (green), F-actin (red), and nuclei (blue) in hDFs. From left to right: resting hDFs, myofibroblast differentiation induced with TGF-β1 and vitamin C, and co-treatment with No Blender or Blender Av-EVs at 2,500 EVs/cell for 72 h. Images were captured at 20X magnification under identical exposure settings within each staining experiment. Quantification of fluorescence intensity for **(C)** collagen I and **(D)** α-SMA. Data represent mean ± SD (n=6 measurements obtained from three independent images per condition). p < 0.05 *, < 0.01 **, < 0.0001 #. Statistical differences relative to the fibrotic control (+TGFꞵ1+VitC/-EVs) are shown in pink, while statistical differences between the isolation methods are shown in black.

Co-treatment with either Av-EV preparation significantly attenuated this response compared with the fibrotic control (p < 0.0001) whose levels were raised to 19,484.8 – 21,539.8. B Av-EVs reduced fluorescence intensity to 7,905.2 – 8,773.2, representing a ∼61% reduction. Nonetheless, the signal remained significantly elevated compared with untreated hDFs (p < 0.001). A stronger response was observed with NB-derived EVs, which reduced fluorescence to 4,024 – 5,646, corresponding to a ∼76% reduction relative to fibrotic cells and restoring collagen I levels to values not significantly different from untreated ones. Thus, although both EV isolation methods effectively limited TGF-β1/vitamin C-induced matrix accumulation, NB Av-EVs produced the greatest normalization under these experimental conditions.

A complementary response was observed for α-SMA (**Figure 11B and D**). Untreated hDFs displayed the characteristic morphology of resting fibroblasts, with elongated spindle-shaped cells and organized F-actin filaments, together with low α-SMA intensities (817.2 to 1,090.2). Fibrotic stimulation induced extensive cytoskeletal remodeling and pronounced colocalization of α-SMA with F-actin, resulting in the formation of prominent contractile stress fibers – a hallmark of differentiated myofibroblasts. Quantification revealed a 2.5-fold rise over untreated cells (p < 0.0001).

Similar to collagen I, both Av-EV treatments significantly attenuated α-SMA expression relative to the fibrotic control (p < 0.0001). NB Av-EV reduced α-SMA levels to 966.8 – 1,474.8, effectively halving the fibrotic effect. Although the cells retained a spread morphology, they exhibited reduced colocalization and loss of organized stress fibers, suggesting diminished contractility and a shift toward a less fibrogenic state. B-derived EVs also decreased α-SMA levels (1,353.3 – 1,610.5), although the effect was less pronounced, resulting in a significant difference between the two isolation methods (p < 0.05). Notably, when compared to untreated cells, NB EV-treated hDFs showed only a modest difference (p < 0.01), whereas B EV-treated cells maintained a stronger divergence (p < 0.0001), further indicating the superior efficacy of NB vesicles in restoring α-SMA expression towards non-fibrotic baseline.

Taken together, the immunofluorescence findings provide complementary protein-level evidence supporting the antifibrotic response observed by RT-qPCR in section 3.2.5. While gene expression measures changes in COL1A1 and ACTA2 transcription, immunofluorescence captures the subsequent accumulation and spatial organization of the proteins. Despite interrogating different levels of the fibrotic response, both approaches yielded directionally consistent findings: TGF-β1/vitamin C enhanced fibrotic gene expression together with matrix accumulation and acquisition of the contractile myofibroblast phenotype, whereas Av-EV treatment attenuated these responses. Notably, NB-derived vesicles consistently produced the strongest normalization toward the non-fibrotic state, further supporting the greater antifibrotic bioactivity of Av-EVs obtained through manual isolation and suggesting that this method may better preserve functionally relevant vesicle components. By simultaneously limiting collagen I accumulation and α-SMA-associated cytoskeletal remodeling, Av-EVs may reduce both excessive ECM deposition and myofibroblast contractility – two central features underlying tissue stiffening and scar contracture – and, therefore, represent a promising plant-derived nanotherapeutic strategy for the modulation of hypertrophic scars and keloids.

### 3.3 Analysis of Cell Responses to NB Av-EVs from Young *Aloe vera* Leaves

All results discussed thus far were obtained using EVs isolated – either manually or mechanically – from mature *Aloe vera* leaves sourced from a supermarket supplied by a large-scale distributor in the United States. These leaves exhibited consistent morphological characteristics (∼67 cm × ∼9 cm × ∼3 cm) and yielded approximately 400 mL per leaf. Given their robust and reproducible performance in reducing pro-inflammatory and fibrotic markers – particularly for NB-derived EVs – we extended our investigation to assess EVs derived from young *Aloe vera* leaves, manually harvested from live potted plants obtained from a local plant shop. These younger leaves were notably smaller (∼45 cm × ∼5 cm × ∼1.5 cm) and produced a reduced gel yield of ∼100 mL per leaf. NTA established that final suspension samples yielded concentrations ranging from 1.2 to 2.8 × 10¹⁰ particles/mL, for a total of 1.2 to 2.8 × 10^12^ particles per leaf. This large disparity in concentration is the first indication that even leaves of the same plant, at different stages of growth, produce and release different amounts of EVs.

This comparative analysis between NB Av-EVs derived from mature and young *Aloe vera* leaves is not only scientifically relevant but also holds important translational implications. Since *Aloe vera* plants require several years to reach full maturity – a factor that directly affects cultivation time, production scalability, and manufacturing costs, understanding whether younger plants can yield vesicles with comparable therapeutic efficacy is critical for future commercialization efforts. Evaluating how plant age and extraction method influence EV composition and bioactivity may help streamline production strategies and inform selection criteria for cost-effective, large-scale Av-EV manufacturing.

#### 3.3.1 Anti-Inflammatory Response (Macrophages)

Co-treatment with NB Av-EVs from young leaves and LPS resulted in attenuation of M1-associated pro-inflammatory cytokines such as CXCL10/IP-10, CCL2/MCP-1, IL-1β, TNF, and IL-6 as shown in **Figure 12**; however, compared to previous results at 24 h (section 3.2.4), it is important to highlight that NB mature leaf-derived EVs consistently outperformed young EVs across all measured markers.

**Figure 12.**
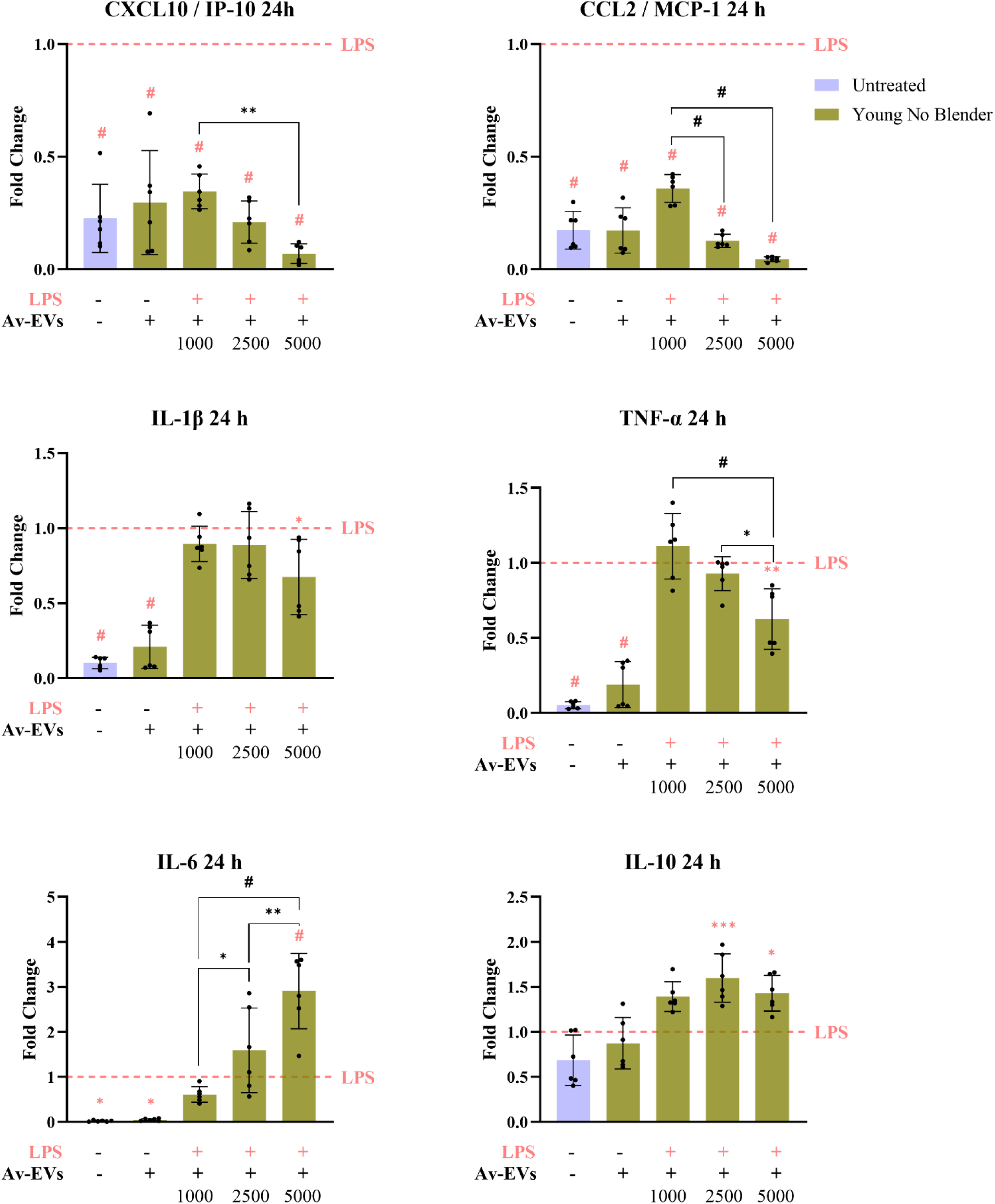
CXCL10/IP-10, CCL2/MCP-1, IL-1β, TNF-α, IL-6 and IL-10 gene expression in LPS-induced macrophages after 24 h of co-treatment with No Blender Av-EVs isolated from young Aloe vera leaves at increasing doses (1,000, 2,500, 5,000 EVs/cell). RT-qPCR data represents the mean ± SD with n=6 and p value < 0.05 *, < 0.01 **, < 0.001 ***, < 0.0001 #. Statistical differences relative to the inflammatory control (+LPS/-EVs) are shown in pink, while statistical differences between doses are shown in black.

CXCL10 and CCL2 were significantly downregulated in a strong dose-dependent manner. Yet, NB-mature EVs achieved near-complete suppression at the lowest dose (fold change values of ≤ 0.126 for CXCL10, and ≤ 0.094 for CCL2), while young EVs required the highest dose to reach comparable levels (≤ 0.021 for CXCL10, and ≤ 0.056 for CCL2), highlighting a marked difference in potency.

While IL-1β and TNF-α exhibited a slightly, dose-dependent downregulation in response to young EVs, their suppression remained consistently inferior to that achieved with mature EVs – with even the lowest dose of mature EVs outperforming the highest dose of young counterparts by achieving statistical significance of p < 0.0001 for both genes vs. p < 0.05 for IL-1β and p < 0.01 for TNF-α, underscoring the limited anti-inflammatory efficacy of young EVs. Interestingly, both expressions were significantly lower in the young EV-only control groups compared to the mature EV control groups, suggesting distinct immune responses by vesicles from plants at different stages of development. Nevertheless, young EVs were unable to restore cytokine expression to baseline levels in the treated cells, in contrast to the superior immunomodulatory capacity of mature EVs.

Surprisingly, IL-6 expression exhibited a clear dose-dependent increase in cells treated with young Av-EVs, reaching fold changes of up to 4 at the highest concentration (p < 0.0001). This response stands in contrast to the stable IL-6 expression observed in mature EV-treated groups at 24 h. Findings that imply that young EVs may not only lack the capacity to suppress IL-6-mediated inflammatory signaling, but at elevated doses, may in fact intensify the inflammatory response, underlying a potential pro-inflammatory liability associated with immature vesicle cargo.

In parallel, IL-10 expression was modestly elevated in cells treated with young Av-EVs, displaying no clear dose-dependent trend and a significant difference of p < 0.001 for the medium and p < 0.05 for the highest dose compared to the LPS-only control, similar response obtained with the mature counterparts at 24 h. However, IL-10 levels in the young EV-only control group were markedly lower than those in the mature EV-only control (fold change values of ≤ 1.314 vs. ≤ 2.848), suggesting that the baseline immunomodulatory potential of vesicles from immature plants may be intrinsically limited.

#### 3.3.2 Antifibrotic Response (Dermal Fibroblasts)

When evaluating fibrotic gene expression (COL1A1 and α-SMA), NB young EVs did not reproduce the modulatory effects on fibroblast-to-myofibroblast transdifferentiation previously observed with NB mature EVs in the same hDF donor (donor 1, lot number 80405992) [**Figure 13**]. Although a dose-dependent pattern was observed, none of the tested young EV concentrations reduced expression below the fibrotic control for either target. Notably, the lowest dose (1,000 EVs/cell) significantly increased COL1A1 expression (p < 0.01). These findings indicate that, under the experimental conditions tested, NB young Av-EVs do not exert a measurable antifibrotic effect, in contrast to NB mature EVs, which effectively attenuated COL1A1 expression at medium and high doses, and α-SMA even at the lowest dose – central mechanisms in pathological scar formation.

**Figure 13.**
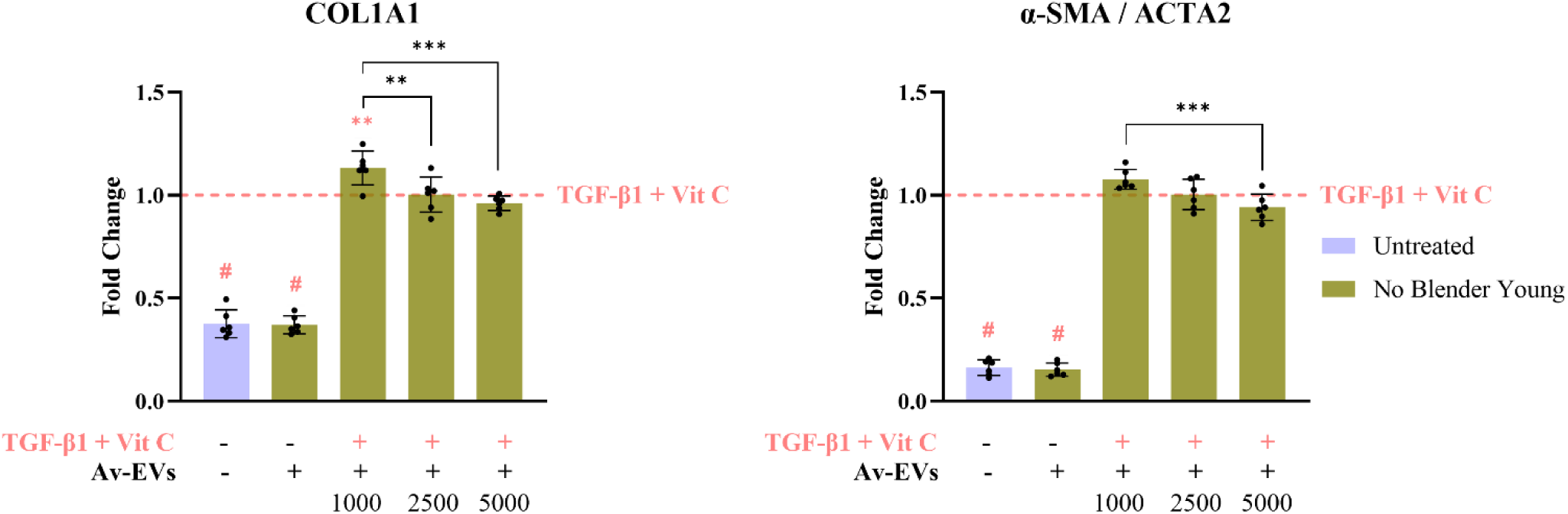
COL1A1 and α-SMA/ACTA2 gene expression in TGF-β1 + vitamin C-activated hDFs after 72 h co-treatment with No Blender Av-EVs isolated from young Aloe vera leaves at increasing doses (1,000, 2,500, 5,000 EVs/cell). RT-qPCR data represents the mean ± SD with n=6 and p value < 0.05 *, < 0.01 **, < 0.001 ***, < 0.0001 #. Statistical differences relative to the fibrotic control (+TGFꞵ1+VitC/-EVs) are shown in pink, while statistical differences between doses are shown in black.

Collectively, this comparative study clearly drew attention to EVs derived from mature *Aloe vera* leaves possess markedly superior anti-inflammatory and antifibrotic properties compared to those from younger plants. Mature Av-EVs consistently outperformed their younger counterparts, achieving stronger downregulation of inflammatory mediators and fibrotic markers. In contrast, young Av-EVs not only exhibited limited efficacy, but in some cases induced paradoxical pro-inflammatory effects, such as increased IL-6 expression at higher doses. These findings align with previous work on orange-derived EVs, where the maturity of the fruit significantly influenced vesicle characteristics, including particle concentration and diameter, surface potential, lipid and protein content, and their ability to encapsulate and deliver bioactive compounds. Specifically, vesicles from ripe orange juice demonstrated superior curcumin delivery performance – enhancing its solubility, stability, bioaccessibility, and antioxidant activity – compared to those from unripe or overripe sources.^[82]^ Given curcumin’s well-established anti-inflammatory properties, this further supports the concept that plant maturity plays a crucial role in optimizing the therapeutic potential of EV-based delivery systems.

#### 3.3.3 Biological Considerations

The disparity in the anti-inflammatory and antifibrotic efficacy observed between mature and young Av-EVs is likely rooted in biological differences in vesicle cargo and surface marker profiles, which are inherently influenced by the plant’s age, developmental stage, and growing conditions.^[83]^ Mature leaves from supermarket-grade *Aloe vera barbadensis miller* plants are typically cultivated under controlled agronomic environments optimized for commercial use. These conditions favor the accumulation of polysaccharides (e.g., acemannan), sterols, flavonoids, and other secondary metabolites that are well-documented for their anti-inflammatory, antioxidant, and immunomodulatory properties.^[84, 85]^ In contrast, younger nursery-grown plants, especially those intended for ornamental purposes, are less metabolically mature. Their smaller biomass, lower gel yield, and underdeveloped biosynthetic pathways likely contribute to a less complex and less bioactive EV cargo as confirmed with the proteomics (section 3.3.4). This includes potential deficits in regulatory lipids, miRNAs, and proteins involved in modulating key inflammatory and fibrotic signaling cascades such as NF-κB and STAT3/STAT6 in macrophages, and TGF-β1/SMAD in fibroblasts. These molecular deficiencies may explain the reduced potency of young Av-EVs in downregulating pro-inflammatory cytokines or reversing myofibroblast-associated gene expression.

These findings highlight the critical role of plant maturity, cultivation purpose, and bioactive composition in determining the therapeutic potential of Aloe-derived bioproducts. In particular, the biochemical and functional properties of Av-EVs can vary substantially depending on factors such as *Aloe* species, plant age, geographical origin, climate, soil type, and extraction methodology.^[86–88]^ Importantly, the optimal harvest time for *Aloe* leaves is typically after three years of growth, when bioactive compound content peaks.^[84]^ Furthermore, post-harvest handling, storage, and transport conditions play a pivotal role in preserving bioactivity, since exposure to excessive light, heat, or humidity can degrade key molecules before vesicle isolation.^[30, 84, 86, 88]^ To address these limitations, the variables mentioned must be carefully controlled to ensure the reproducibility and potency of EV-based therapeutics derived from *Aloe* plant sources, streamlining both clinical translation and regulatory approval.

#### 3.3.4 Differentially Expressed Proteins (DEPs) in Av-EVs based on Plan Maturity

After confirming plant-specific EV markers (see section 3.1), a comparison of Av-EV protein cargo stratified by leaf age – mature (M1–M3) versus young (Y1–Y3) – was sought to provide mechanistic context for the performance gap between preparations.

Across all samples, 5,376 proteins (38,890 peptides) were identified. Label-free quantification followed by PCA revealed tight intragroup clustering with clear separation by leaf age [**Figure 14A**]. Statistical testing yielded 583 significantly differentially expressed proteins (DEPs), with 309 increased (red) and 274 decreased (blue) in mature EVs [**Figure 14B**]. This marker landscape suggests that leaf age is a key determinant of Av-EV protein cargo and supports the notion that mature Av-EVs are produced and trafficked through more robust secretory, cytoskeletal and stress-adaptive pathways than young Av-EVs; thus, providing a coherent explanation for their superior bioactivity in macrophage repolarization and suppression of fibroblast activation.

**Figure 14.**
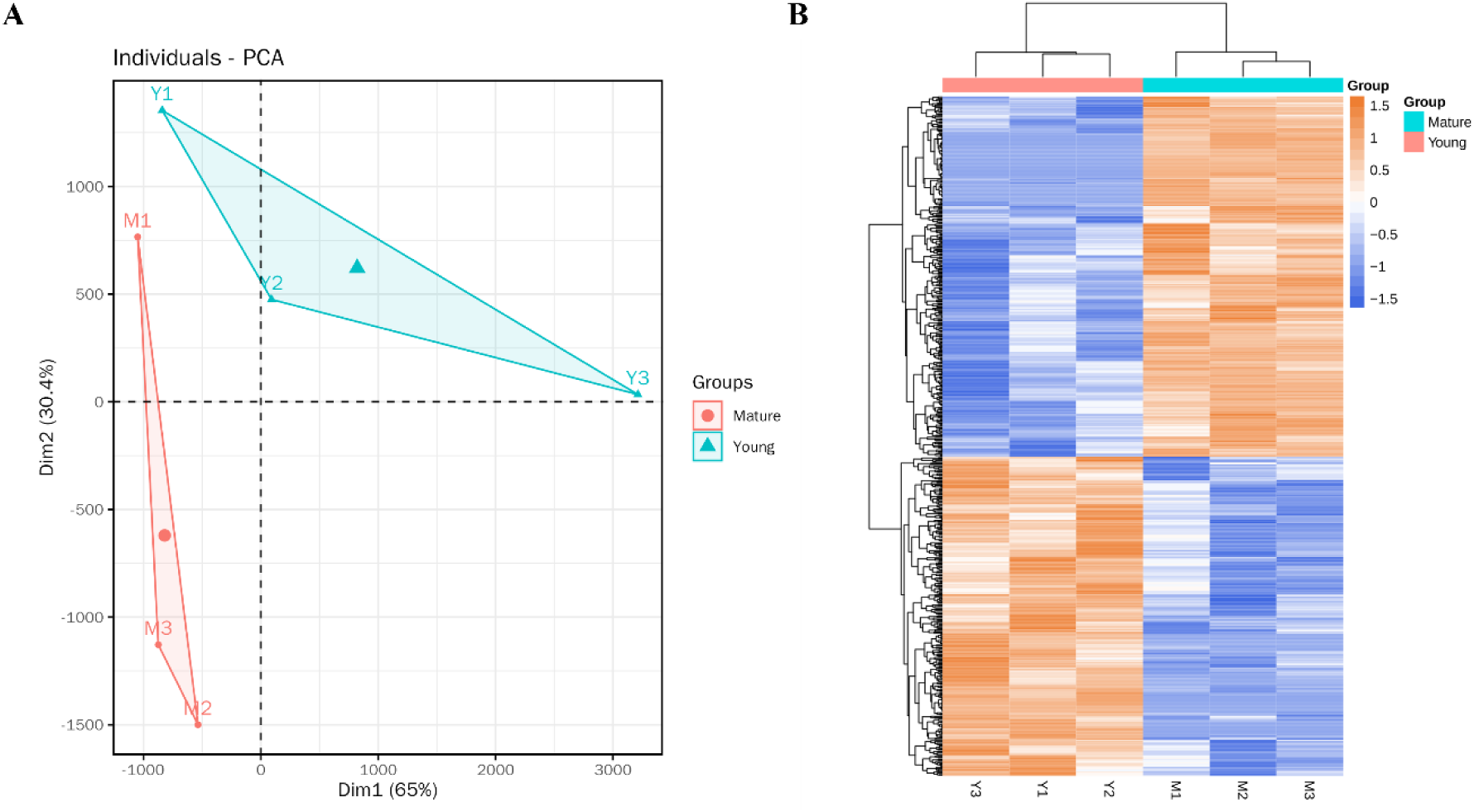
Proteomic cargo comparison of Av-EVs isolated from mature (M1-M3) vs. young (Y1-Y3) leaves. (A) PCA. (B) Heatmap of DEPs.

##### GO/KEGG Enrichment Related with the EV Biogenesis, Trafficking, and Uptake Sequence

GO/KEGG enrichment analysis revealed that the plant-specific marker families described above were among the DEPs, distinguishing EVs derived from mature versus young leaves. Indeed, higher representation in mature Av-EVs of these terms help to support vesicle biogenesis and stabilize cargo during isolation and after uptake (HSP70/90);^[89, 90]^ strengthen actin/clathrin-dependent budding and trafficking across the cell wall;^[91–95]^ enhance targeting, docking, and fusion – and thus cargo delivery – via Rab GTPases and syntaxin/SNARE (soluble N-ethylmaleimide–sensitive factor attachment protein receptor) complexes;^[96–98]^ ^[99–101]^ and provide Ca²⁺/osmotic and redox-responsive protection and tuning of downstream signaling (annexins).^[102–105]^ Taken together, this upregulated marker profile offers a mechanistic basis for the superior functional performance of mature Av-EVs.

##### GO/KEGG Directionality in the Context of Anti-inflammatory and Antifibrotic Functions

To relate proteomics to antifibrotic and anti-inflammatory functions, we categorized each GO term and KEGG pathway based on the differential abundance of the majority of its corresponding IDs. Terms whose constituent proteins were mostly increased in mature Av-EVs are discussed as “upregulated”; those whose IDs were mostly decreased in mature are discussed as “downregulated”. The most representative GO molecular-function terms are summarized in **Figure 15**, and the KEGG pathways in **Figure 16**.

**Figure 15.**
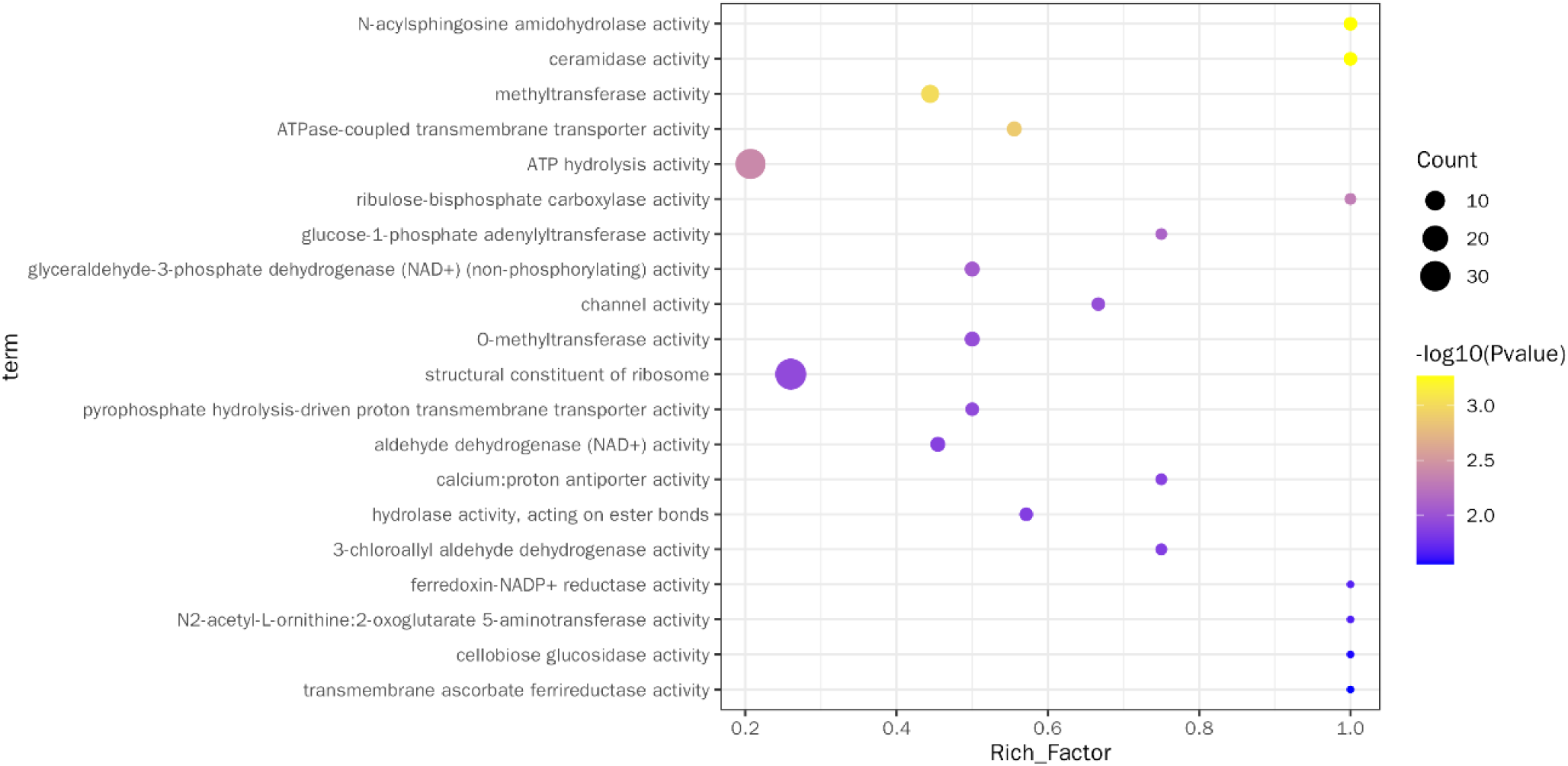
GO molecular function enrichment for proteins in No Blender Av-EVs (isolated from mature and young Aloe vera leaves).

**Figure 16.**
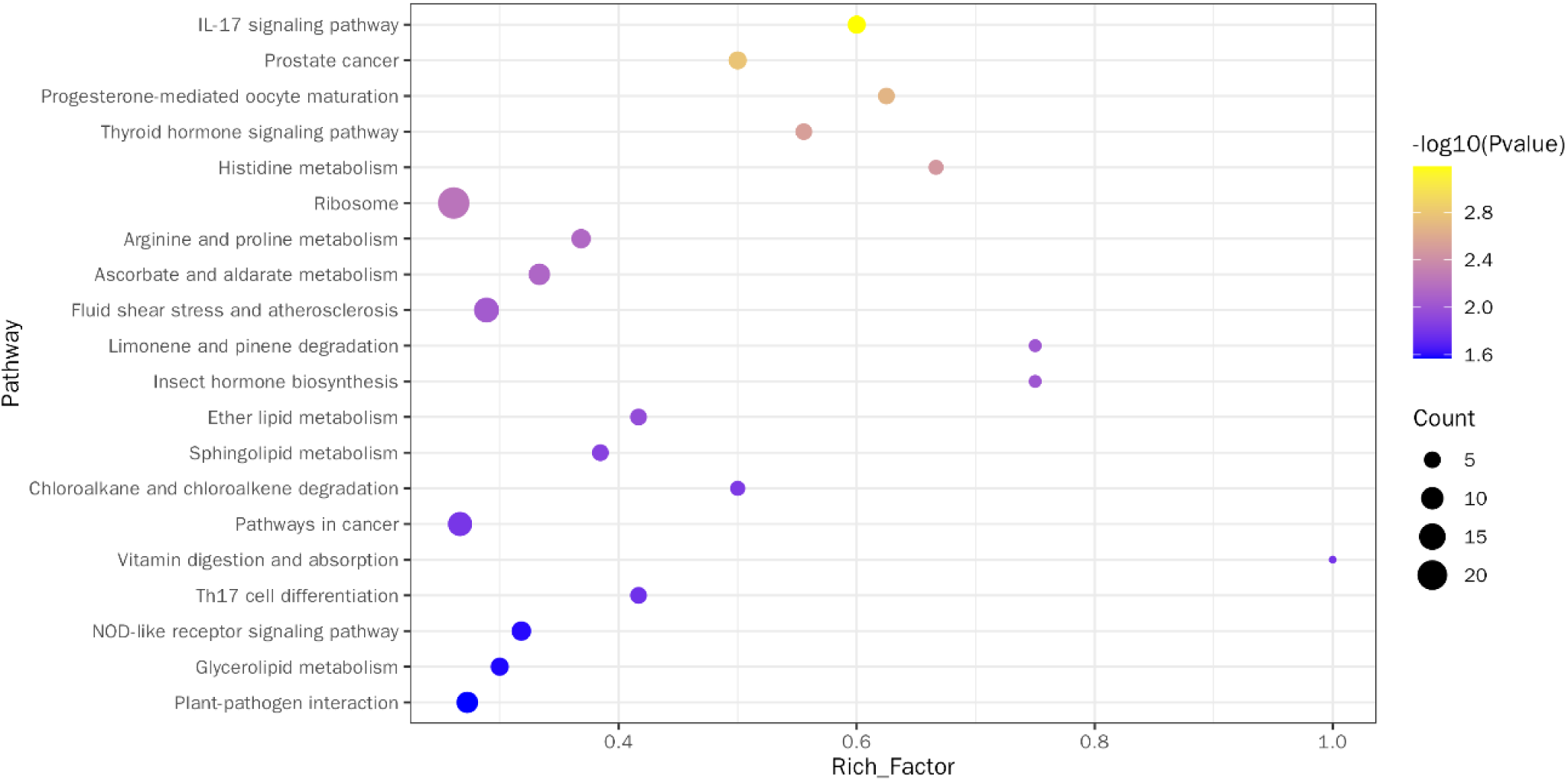
KEGG pathway enrichment for proteins in No Blender Av-EVs (isolated from mature and young Aloe vera leaves).

###### GO Enrichment Terms Upregulated in Mature Av-EVs

**- Ceramidase axis (N-acylsphingosine amidohydrolase / ceramidase activity).** Ceramides are bioactive lipids that promote inflammation and apoptosis, whereas sphingosine-1-phosphate (S1P) supports cell survival and tissue repair. Enrichment of ceramidase activity in mature EVs is consistent with a shift from ceramide → sphingosine → S1P. Functionally, lower ceramide and appropriately tuned S1P would be expected to lessen RhoA/ROCK signaling and weakening stress-fiber formation, while in immune cells, this balance typically reduces NF-κB/NLRP3 activation and favors M2-like, IL-10/ARG1-positive phenotypes. ^[106–109]^
**- Calcium handling (channel activity; calcium:proton antiporter activity; pyrophosphate-driven proton pumps).** Together these terms point to tighter cytosolic Ca²⁺ control. Limiting Ca²⁺ influx/redistribution would be expected to blunt Ca²⁺-dependent myosin light-chain kinase activity (MLCK) and myocardin-related transcription factor–serum response factor (MRTF–SRF) pathways, reducing α-SMA assembly and contractile tone in fibroblasts, and to dampen Ca²⁺-dependent steps of NF-κB/NLRP3 activation in macrophages – aligning with lower IL-1β/TNF and stabilized IL-6.^[110–112]^
**- Membrane remodeling (ATPase-coupled transmembrane transporter activity).** This term points to ABC transporters and P4-ATPase flippases that reset lipid asymmetry and lipid-raft architecture. In fibroblasts, such remodeling can reduce integrin clustering and focal adhesion kinase (FAK) activation, which in turn influences EV uptake, adhesion dynamics, and downstream cytoskeletal tension in recipient cells.^[113]^ In macrophages, exposure of phosphatidylserine and raft reorganization can promote IL-10 production and temper TLR/NF-κB signaling.
**- Reactive-aldehyde detox (aldehyde dehydrogenase, NAD⁺ activity).** Greater capacity to clear 4-HNE and related aldehydes would lower ROS-propagated NF-κB/TGF-β throughput, a common feed-forward driver of both myofibroblast differentiation and pro-inflammatory cytokine release.^[114, 115]^

###### GO Enrichment Terms Downregulated in Mature Av-EVs

**- Proline supply (N2-acetyl-L-ornithine:2-oxoglutarate 5-aminotransferase activity).** This enzyme lies on the arginine→ornithine→proline axis, specifically in the acetylated branch of *de novo* arginine biosynthesis. Its depletion means that fewer mature-EV proteins map to this specific biosynthetic step than in young EVs. In other words, mature EVs are not enriched for an upstream enzyme that builds arginine from scratch, but rather skew toward arginine and proline metabolism as evidenced in the KEGG enrichment pathways, where it coupled to a more regulated proline supply in fibroblasts, moderating collagen synthesis and aligning immune resolution with controlled matrix deposition.^[116–119]^ The ARG1 increase observed in macrophages (see section 3.2.4), therefore, likely reflects recipient-cell induction by mature-EV cues that dampen NF-κB signaling and enhance the pathways that bias macrophages toward an M2 phenotype, rather than direct enzyme delivery.
**- Protein-synthesis capacity (structural constituent of ribosome).** A young-skewed ribosomal signature suggests a higher translational drive that could support increased ECM production and cytokine synthesis; its attenuation in mature EVs is directionally consistent with less matrix output and a calmer cytokine milieu.^[120–122]^

###### KEGG Enrichment Terms Upregulated in Mature Av-EVs

**- Ether lipid metabolism.** Plasmalogen-rich membranes bolster antioxidant buffering and stabilize lipid microdomains, thereby attenuating TGF-β–sensitized actomyosin stress-fiber assembly and blunting TLR–NF-κB signaling.^[123–125]^
**- Sphingolipid metabolism.** Re-tuning the ceramide → sphingosine → S1P pools away from ceramide is predicted to reduce inflammasome/NF-κB activation and soften cytoskeletal tension, aligning with lower α-SMA and reparative macrophage phenotypes as described in the ceramidase GO enrichment term.^[106–109]^
**- Regulation of actin cytoskeleton.** Enrichment of actin-regulatory cargo is coherent with direct constraints on α-SMA assembly and myofibroblast contractility.^[91–93]^
**- Arginine and proline metabolism.** The enrichment suggests tuning of the arginine→ornithine→proline axis where mature EVs may facilitate controlled proline flux that supports tissue repair without excessive collagen deposition.^[116–119]^
**- IL-17 signaling pathway.** Several proteins mapping to nodes in IL-17 pathway function as modulators or brakes. Thus, their enrichment is compatible with a capacity to tune the IL-17 axis downward in recipient cells, helping to reduce the IL-17↔IL-6↔TGF-β1 feed-forward loop that sustains myofibroblast differentiation.^[126, 127]^

###### KEGG Enrichment Terms Downregulated in Mature Av-EVs

**- Fluid shear stress and atherosclerosis.** Lower representation of this composite mechanotransduction/ROS/NF-κB module is consistent with less cytoskeletal tension and lower MAPK/NF-κB tone.^[128, 129]^
**- NOD-like receptor signaling.** Reduced NOD/NLR inputs predict less inflammasome activation and lower IL-1β maturation, matching the cytokine profile observed in the anti-inflammatory response.^[130, 131]^
**- Ribosome.** Depletion of translation machinery is coherent with lower ECM protein output and attenuated cytokine synthesis as also evidenced by the downregulation of the structural constituent of ribosome GO term.^[120–122]^
**- Ascorbate and aldarate metabolism.** A lower metabolism in mature EVs could limit effective endoplasmic reticulum ascorbate available for collagen prolyl/lysyl hydroxylases, thereby curbing collagen maturation.^[132, 133]^
**- Toll-like receptor, MAPK, PI3K–Akt, and JAK–STAT signaling.** Across these pathways, reduced representation in mature EVs would be expected to lower MyD88/TRAF6→NF-κB/MAPK throughput, temper AP-1–driven cytokines and reduce STAT-linked inflammatory transcription – all consistent with lower CXCL10/CCL2/IL-1β/TNF/IL-6 and increase of IL-10/ARG1.^[134–138]^

Viewed together, the enrichment landscape offers a unified mechanistic narrative. From a fibrosis-centric perspective, mature Av-EVs are enriched for modules that reduce contractility (actin regulation, Ca²⁺ control), reprogram membranes (ether/sphingolipid remodeling), and detoxify aldehydes, while being relatively depleted in proline-supply and ribosomal terms. This directional pattern bridges proteomics to phenotype by explaining the reductions in COL1A1 and α-SMA we observed. In parallel, an immune-centric reading shows that terms increased in mature EVs converge on lipid signaling, Ca²⁺ control, redox detox, and tunable IL-17–pathway nodes, whereas terms decreased in mature EVs include NOD/TLR/MAPK/JAK–STAT modules. Together, that tilt rationalizes the lower chemokines/cytokines, stabilized IL-6, and higher IL-10/ARG1.

In sum, the proteomic signature of mature Av-EVs indicates that they are produced and trafficked through more robust secretory, cytoskeletal, and stress-adaptive pathways than vesicles from young leaves. This profile offers a coherent mechanistic link between EV cargo and redox control, immune modulation, and matrix regulation consistent with the age-dependent bioactivity. For instance, mature Av-EVs uniquely reduced α-SMA and COL1A1 in TGF-β1/vitamin C–stimulated fibroblasts and attenuated pro-inflammatory cytokines in M1 macrophages at markedly lower doses (1,000 vs 5,000 EVs per cell). Together, these findings identify leaf maturity as a decisive variable of Av-EV composition and function, and they motivate standardization of source maturity alongside targeted validation of candidate proteins to ensure reproducible therapeutic performance.

## 4. Conclusion

This study presents the first comprehensive characterization of *Aloe vera* gel-derived extracellular vesicles (Av-EVs), including the first proteomic atlas for this EV class, with emphasis on how isolation method and plant maturity shape their physicochemical and bioactive profiles, and thereby their capacity to modulate key pathological mechanisms implicated in abnormal scar formation: chronic inflammation, oxidative stress, and myofibroblast differentiation – three interlinked drivers of fibrotic remodeling.

Av-EVs were successfully isolated by both manual and shear-force-based methods and compared across mature and young *Aloe vera* leaves. Notably, Av-EVs manually extracted from mature leaves yielded superior vesicle bioactivity *in vitro* as evidenced in the attenuation of pro-inflammatory cytokine expression in M1-polarized macrophages (CXCL10, CCL2, IL-1β, TNF-α, IL-6) alongside the promotion of an M2-like phenotype (IL-10, ARG1), time-dependent inhibition of free radical levels, and suppression of myofibroblast differentiation in activated human dermal fibroblasts (COL1A1, α-SMA). These results suggest that gentle extraction may have an advantage in retaining and preserving the bioactive components require to break the inflammation–oxidative stress–fibrosis cycle and restore tissue homeostasis.

Proteomic profiling established a plant-typical EV signature and mapped age-dependent mechanisms. Across samples we identified 5,376 proteins (38,890 peptides); PCA cleanly separated groups by leaf age, and 583 DEPs were detected (309 up, 274 down in mature EVs). Marker families repeatedly reported in plant EVs – HSP70/HSP90, actin-cytoskeleton machinery, syntaxin/SNARE, small GTPases, clathrin heavy chain, annexins, and cell-wall remodeling – were all present, confirming EV identity. KEGG/GO directionality then linked cargo to function: mature EVs were enriched in regulation of actin cytoskeleton and Ca²⁺-handling/channel activities (reduced contractility), ether/sphingolipid/ metabolism (membrane remodeling and antioxidant buffering), ceramidase activity (S1P shift to support cell survival and tissue repair), and aldehyde dehydrogenase (NAD⁺) (ROS detox). Conversely, mature EVs showed lower representation of ribosome-associated translation capacity, proline-supply nodes, and innate/inflammatory modules (NOD/TLR/MAPK/PI3K–Akt/JAK–STAT). Together with our gene-expression data, this maturity-biased proteome supports a model in which mature Av-EVs arise from more robust secretory, cytoskeletal, and stress-adaptive programs and deliver cargo that dampens fibroblast contractility and inflammatory throughput – molecular features absent from young counterparts. Immunoblotting further supported Av-EV identity through detection of the EV-associated marker SYP121/PEN1 and absence of a discrete calnexin signal, providing complementary protein-level evidence of a plant EV-enriched preparation with limited detectable ER-associated contamination.

While limited to *in vitro* contexts, these findings underscore the importance of standardizing the isolation protocol and source material for reproducible outcomes. Most importantly, they position Av-EVs as natural, scalable, cell-free nanotherapeutics that warrant further investigation in underlying mechanistic studies, *in vivo* validation, and advanced delivery systems to enable clinical translation for preventing and treating fibrosis-related skin disorders.

## Acknowledgments

The authors gratefully acknowledge the members of the Abhyankar Lab Biological Microsystems at Rochester Institute of Technology for their technical assistance with fluorescence microscopy, especially Ph.D. candidate Poorya Esmaili and Dr. Indranil "Neil" Joshi.

## Data Availability Statement

Mass spectrometry–based proteomics data comparing NB Av-EVs isolated from mature and young *Aloe vera* leaves have been deposited in the ProteomeXchange Consortium via the PRIDE partner repository under the dataset identifier PXD069742. All other datasets generated during this study are available through the Figshare repository DOI:10.6084/m9.figshare.29367212.

## Supplementary Methods

### Donor-Resolved Analysis of hDF Gene Expression in Response to Av-EVs Isolated from Mature *Aloe vera* Leaves

Gene expression experiments evaluating hDF responses to Av-EVs isolated from mature *Aloe vera* leaves included two independent biological donors. For each donor, all treatment conditions were evaluated in three independent experimental replicates, with each replicate performed on a separate day and plate using an independently prepared Av-EV isolation sample. Cell culture, fibrotic induction, Av-EV treatment, RNA isolation, RT-qPCR, and relative gene expression analysis were performed as described in the corresponding Methods sections of the main manuscript (2.2.2, 2.3.4, and 2.4.1).

To specifically evaluate inter-donor variability, the donor-resolved data were additionally analyzed using a mixed-effects model with restricted maximum likelihood (REML), specifying treatment as a fixed effect and donor as a random effect, with independent experimental replicates nested within donor. This analysis complemented the one-way ANOVA presented in the main manuscript. Donor-specific responses and the corresponding statistical comparisons are presented in **Figure 1S**.

## Supplementary Results

### Donor-Specific Antifibrotic Responses

Donor-resolved analysis revealed differences in the magnitude of the antifibrotic response between the two hDF donors; however, the overall treatment pattern remained consistent with the findings presented in the main manuscript.

Relative to the fibrotic control (+TGF-β1 + VitC/–EVs), untreated hDFs displayed low baseline levels of COL1A1 (≤ 0.409 for donor 1 and ≤ 0.351 for donor 2) and α-SMA (≤ 0.216 for donor 1 and ≤ 0.116 for donor 2), where “≤” indicates that the reported value represents the highest observed level within that condition, consistent with a non-fibrotic state (p < 0.0001). Similarly, EV-only conditions remained comparable to untreated cells and were also significantly lower than the fibrotic control (p < 0.0001), indicating that Av-EVs did not induce fibrotic activation in resting fibroblasts. Under these basal conditions, NB-derived EVs showed consistent levels between donors for both COL1A1 and α-SMA, with no significant inter-donor differences. In contrast, B-derived EVs exhibited modest donor-dependent variability, with higher COL1A1 and α-SMA levels in donor 1 compared to donor 2 (p < 0.01 and p < 0.05, respectively), indicating greater donor-dependent variability in the response to B-derived EVs.

Upon fibrotic stimulation, treatment with NB Av-EVs led to a pronounced attenuation of TGF-β1/VitC-induced expression of both COL1A1 and α-SMA. Donor 1 (●, solid bars) exhibited significant reductions in COL1A1 at the intermediate and high doses (p < 0.0001 and p < 0.01, respectively) and in α-SMA/ACTA2 at both doses (p < 0.0001). Donor 2 (▲, patterned bars) showed a more moderate response, with significant reductions in both markers at the highest dose (p < 0.05). These differences resulted in significant inter-donor variability for some treatment conditions, reflecting the expected biological heterogeneity of primary human fibroblasts.

B-derived Av-EVs showed a more limited response. No significant reduction in COL1A1 was detected across doses in either donor, whereas α-SMA/ACTA2 was significantly reduced in donor 2 at the intermediate and high doses (p < 0.01 and p < 0.0001, respectively). Direct comparison of the isolation methods within donor 1 further favored NB-derived Av-EVs, particularly at the intermediate dose for COL1A1 (p < 0.0001) and at the intermediate and high doses for α-SMA/ACTA2 (p < 0.0001 and p < 0.01, respectively).

Overall, although donor-dependent differences were evident, the donor-resolved analysis supported the principal conclusion of the main manuscript: NB-derived Av-EVs, particularly at the intermediate and high doses, produced the strongest and most consistent attenuation of the fibrotic response. These findings further support an influence of the isolation method on the biological activity of Av-EVs while highlighting the intrinsic variability of primary hDF responses.

**Figure 1S.**
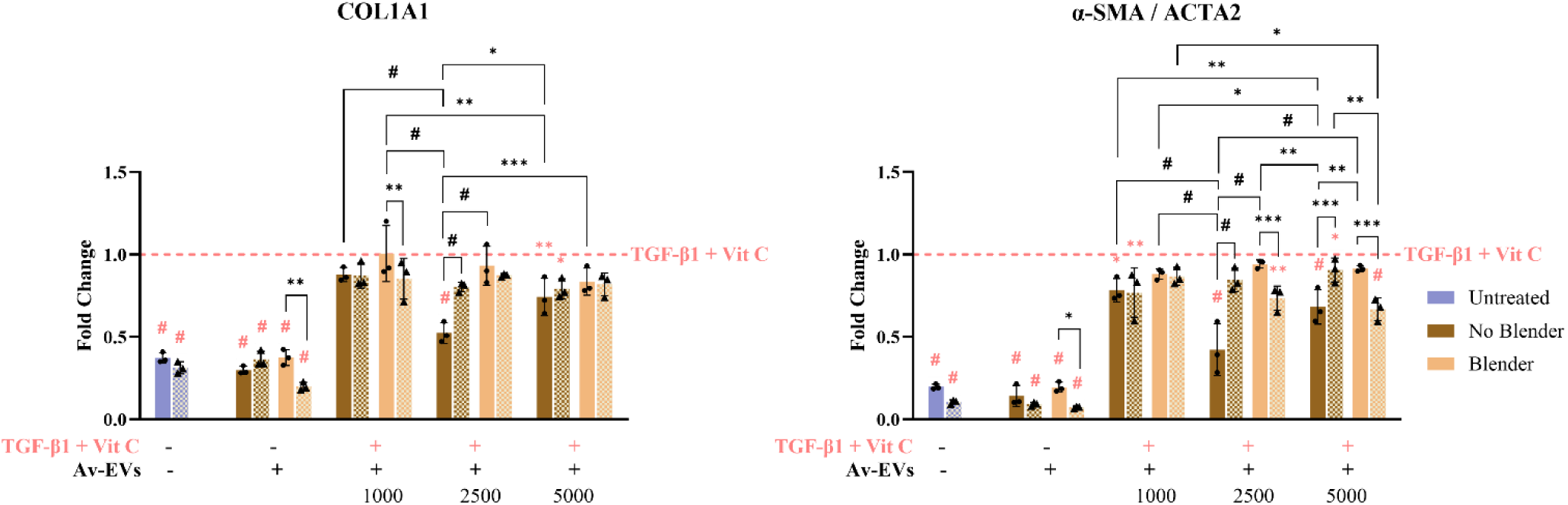
Donor-resolved analysis of COL1A1 and α-SMA/ACTA2 gene expression in TGF-β1 + vitamin C-activated hDFs after 72 h co-treatment with No Blender or Blender Av-EVs at increasing doses (1,000, 2,500, 5,000 EVs/cell). hDFs from two independent biological donors are shown separately: donor 1 (●, solid bars; lot number 80405992) and donor 2 (▲, patterned bars; lot number 80124232), with three independent experimental replicates per donor for each treatment condition (n=3 per donor). Each experimental replicate was performed on a separate day and plate using an independently prepared Av-EV isolation sample. Data are presented as mean ± SD. Donor-resolved data were analyzed using a mixed-effects model (REML), with treatment specified as a fixed effect and donor as a random effect, with experimental replicates nested within donor. Statistical significance is indicated as *p < 0.05, **p < 0.01, ***p < 0.001, and #p < 0.0001. Comparisons relative to the fibrotic control (+TGF-β1 + VitC/–EVs) are shown in pink, whereas comparisons between isolation methods and doses are shown in black.

## References

[1] M. Xue and C. Jackson, "Extracellular Matrix Reorganization During Wound Healing and Its Impact on Abnormal Scarring," Advances in Wound Care, vol. 4, no. 3, pp. 119–136, 2015, doi: 10.1089/wound.2013.0485.

[2] A. Shroff, A. Mamalis, and J. Jagdeo, "Oxidative Stress and Skin Fibrosis," Current Pathobiology Reports, vol. 2, no. 4, pp. 257–267, 2014, doi: 10.1007/s40139-014-0062-y.

[3] M. Mittal, M. R. Siddiqui, K. Tran, S. P. Reddy, and A. B. Malik, "Reactive Oxygen Species in Inflammation and Tissue Injury," Antioxid Redox Signaling, vol. 20, no. 7, pp. 1126–1167, 2014, doi: 10.1089/ars.2012.5149.

[4] S. Antar, N. Ashour, M. Marawan, and A. Al-Karmalawy, "Fibrosis: Types, Effects, Markers, Mechanisms for Disease Progression, and Its Relation with Oxidative Stress, Immunity, and Inflammation," International Journal of Moleculas Sciences, vol. 24, no. 4, p. 4004, 2023, doi: 10.3390/ijms24044004.

[5] S. Piera-Velasquez and S. A. Jimenez, "Oxidative Stress Induced by Reactive Oxygen Species (ROS) and NADPH Oxidase 4 (NOX4) in the Pathogenesis of the Fibrotic Process in Systemic Sclerosis: A Promising Therapeutic Target," Journal of Clinical Medicine, vol. 10, no. 20, p. 4791, 2021, doi: 10.3390/jcm10204791.

[6] R. Kumar, A. K. Singh, A. Gupta, A. Bishayee, and A. K. Pandey, "Therapeutic potential of Aloe vera—A miracle gift of nature," Phytomedicine, vol. 60, p. 152996, 2019, doi: 10.1016/j.phymed.2019.152996.

[7] D. Sánchez-Machado, J. López-Cervantes, R. Sendón, and A. Sanches-Silva, "Aloe vera: Ancient knowledge with new frontiers," Trends in Food Science & Technology, vol. 61, pp. 94–102, 2017, doi: 10.1016/j.tifs.2016.12.005.

[8] M. H. Radha and N. P. Laxmipriya, "Evaluation of biological properties and clinical effectiveness of Aloe vera: A systematic review," Journal of Traditional and Complementary Medicine, vol. 5, no. 1, pp. 21–26, 2015, doi: 10.1016/j.jtcme.2014.10.006.

[9] R. Pereira, A. Carvalho, D. Vaz, D. Gil, A. Mendes, and P. Bártolo, "Development of novel alginate based hydrogel films for wound healing applications," International Journal of Biological Macromolecules, vol. 52, pp. 221–230, 2013, doi: 10.1016/j.ijbiomac.2012.09.031.

[10] M. Sánchez, E. González-Burgos, I. Iglesias, and M. P. Gómez-Serranillos, "Pharmacological Update Properties of Aloe Vera and its Major Active Constituents," Molecules, vol. 25, no. 6, p. 1324, 2020, doi: 10.3390/molecules25061324.

[11] J. H. Hamman, "Composition and Applications of Aloe vera Leaf Gel," Molecules, vol. 13, no. 8, pp. 1599–1616, 2008, doi: 10.3390/molecules13081599.

[12] W. Alawad, M. Othman, J. Hsaian, and L. Alsabek, "Effect of aloe vera hydrogel on the scarring and healing of free gingival graft: Randomized controlled trial," Dentistry 3000, vol. 10, no. 1, 2022, doi: 10.5195/d3000.2022.225.

[13] N. Zhang et al., "Modulating cationicity of chitosan hydrogel to prevent hypertrophic scar formation during wound healing," International Journal of Biological Macromolecules, vol. 154, pp. 835–843, 2020, doi: 10.1016/j.ijbiomac.2020.03.161.

[14] P. Danish, Q. Ali, M. M. Hafeez, and A. Malik, "Antifungal and Antibacterial Activity of Aloe vera Plant Extract," Biological and Clinical Sciences Research Journal, vol. 4, 2020, doi: 10.54112/bcsrj.v2020i1.4.

[15] D. Hekmatpou, F. Mehrabi, K. Rahzani, and A. Aminiyan, "The Effect of Aloe Vera Clinical Trials on Prevention and Healing of Skin Wound: A Systematic Review," Iranian Journal of Medical Sciences, vol. 44, no. 1, pp. 1–9, 2019.

[16] S. Singh, A. Gupta, and B. Gupta, "Scar free healing mediated by the release of aloe vera and manuka honey from dextran bionanocomposite wound dressings," International Journal of Biological Macromolecules, vol. 120, pp. 1581–1590, 2018, doi: 10.1016/j.ijbiomac.2018.09.124.

[17] M. Hormozi, R. Assaei, and M. B. Boroujeni, "The effect of aloe vera on the expression of wound healing factors (TGFβ1 and bFGF) in mouse embryonic fibroblast cell: In vitro study," Biomedicine & Pharmacotherapy, vol. 88, pp. 610–616, 2017, doi: 10.1016/j.biopha.2017.01.095.

[18] A. Oryan, A. Mohammadalipour, A. Moshiri, and M. R. Tabandeh, "Topical Application of Aloe vera Accelerated Wound Healing, Modeling, and Remodeling," Annals of Plastic Surgery, vol. 77, no. 1, pp. 37–46, 2016, doi: 10.1097/SAP.0000000000000239.

[19] M. Avijgan, A. Kamran, and A. Abedini, "Effectiveness of Aloe Vera Gel in Chronic Ulcers in Comparison with Conventional Treatments," Iranian Journal of Medical Sciences, vol. 41, 2016.

[20] F. Eghdampour, F. Jahdie, M. Kheyrkhah, M. Taghizadeh, S. Naghizadeh, and H. Hagani, "The Impact of Aloe vera and Calendula on Perineal Healing after Episiotomy in Primiparous Women: A Randomized Clinical Trial," Journal of Caring Sciences, vol. 2, no. 4, pp. 279–286, 2013, doi: 10.5681/jcs.2013.033.

[21] M. M. Budai, A. Varga, S. Milesz, J. Tozser, and S. Benko, "Aloe vera downregulates LPS-induced inflammatory cytokine production and expression of NLRP3 inflammasome in human macrophages," no. 56, pp. 471–479, 2013, doi: 10.1016/j.molimm.2013.05.005.

[22] F. Nejatzadeh-Barandozi, "Antibacterial activities and antioxidant capacity of Aloe vera," Organic and Medical Chemistry Letters, vol. 3, no. 5, 2013, doi: 10.1186/2191-2858-3-5.

[23] D. Saini, Saini, M., "Evaluation of radioprotective efficacy and possible mechanism of action of Aloe gel," Environmental Toxicology and Pharmacology, vol. 31, no. 3, pp. 427–435, 2011, doi: 10.1016/j.etap.2011.02.004.

[24] N. Yun, C.-H. Lee, and S. M. Lee, "Protective effect of Aloe vera on polymicrobial sepsis in mice," Food and Chemical Toxicology, vol. 47, pp. 1341–1348, doi: 10.1016/j.fct.2009.03.013.

[25] F. Habeeb et al., "The inner gel component of Aloe vera suppresses bacterial-induced pro-inflammatory cytokines from human immune cells," Methods, vol. 42, no. 4, pp. 388–393, 2007, doi: 10.1016/j.ymeth.2007.03.005.

[26] K. Eamlamnam, S. Patumraj, N. Visedopas, and D. Thong-Ngam, "Effects of Aloe vera and sucralfate on gastric microcirculatory changes, cytokine levels and gastric ulcer healing in rats," World Journal of Gastroenterology, vol. 12, no. 13, pp. 2034–2039, 2006, doi: 10.3748/wjg.v12.i13.2034.

[27] R. Prabjone, D. Thong-Ngam, N. Wisedopas, T. Chatsuwan, and S. Patumraj, "Anti-inflammatory effects of Aloe vera on leukocyte–endothelium interaction in the gastric microcirculation of Helicobacter pylori-infected rats," Clinical Hemorheology and Microcirculation, vol. 35, no. 3, pp. 359–366, 2006.

[28] R. J. Langmead and D. S. Makins, "Anti-inflammatory effects of aloe vera gel in human colorectal mucosa in vitro," Alimentary Pharmacology and Therapeutics, vol. 19, no. 5, pp. 521–527, 2004, doi: 10.1111/j.1365-2036.2004.01874.x.

[29] S. W. Woo, J.-X. Nan, S. H. Lee, E.-J. Park, Y. Z. Zhao, and D. H. Sohn, "Aloe Emodin Suppresses Myofibroblastic Differentiation of Rat Hepatic Stellate Cells in Primary Culture," Pharmacology & Toxicology, vol. 90, no. 4, pp. 193–198, 2002, doi: 10.1034/j.1600-0773.2002.900404.x.

[30] F. A. Alzahrani, M. I. Khan, N. Kameli, E. Alsahafi, and Y. M. Riza, "Plant-Derived Extracellular Vesicles and Their Exciting Potential as the Future of Next-Generation Drug Delivery," Biomolecules, vol. 13, no. 5, p. 839, 2023, doi: 10.3390/biom13050839.

[31] M. Kim, H. Jang, W. Kim, D. Kim, and J. H. Park, "Therapeutic Applications of Plant-Derived Extracellular Vesicles as Antioxidants for Oxidative Stress-Related Diseases," Antioxidants, vol. 12, no. 6, p. 1286, 2023, doi: 10.3390/antiox12061286.

[32] X. Liu, Lou, K., Zhang, Y., Li, C., Wei, S., Feng, S., "Unlocking the Medicinal Potential of Plant-Derived Extracellular Vesicles: current Progress and Future Perspectives," International Journal of Nanomedicine, vol. 19, pp. 4877–4892, 2024, doi: 10.2147/IJN.S463145.

[33] E. Calzoni, Bertoldi, A., Cusumano, G., Buratta, S., Urbanelli, L., Emiliani, C., "Plant-Derived Extracellular Vesicles: Natural Nanocarriers for Biotechnological Drugs," Processes, vol. 12, no. 12, p. 2938, 2024, doi: 10.3390/pr12122938.

[34] L. Bahmani, Ullah, M., "Different Sourced Extracellular Vesicles and Their Potential Applications in Clinical Treatments," Cells, vol. 11, no. 13, p. 1989, 2022, doi: 10.3390/cells11131989.

[35] O. Ramírez et al., "Aloe vera peel-derived nanovesicles display anti-inflammatory properties and prevent myofibroblast differentiation," Phytomedicine, vol. 122, p. 155108, 2024, doi: 10.1016/j.phymed.2023.155108.

[36] M. Kim and J. H. Park, "Isolation of Aloe saponaria-Derived Extracellular Vesicles and Investigation of Their Potential for Chronic Wound Healing," Pharmaceutics, vol. 14, p. 1905, 2022, doi: https://doi.org/10.3390/.

[37] M. K. Kim, Y. C. Choi, S. H. Cho, J. K. Choi, and Y. W. Cho, "The Antioxidant Effect of Small Extracellular Vesicles Derived from Aloe vera Peels for Wound Healing," Tissue Engineering and Regenerative Medicine, vol. 18, no. 4, pp. 561–571, 2021, doi: 10.1007/s13770-021-00367-8.

[38] L. Zeng et al., "Aloe derived nanovesicle as a functional carrier for indocyanine green encapsulation and phototherapy," Journal of Nanobiotechnology, vol. 439, 2021, doi: 10.1186/s12951-021-01195-7.

[39] M. C. Ceballos-Santa, Sanchez-Ortiz, I., Gaborski, T.R., Wuertz-Kozak, K., "A Step-by-Step Protocol for the Isolation of Aloe vera–Derived Extracellular Vesicles via Manual and Shear-Force Homogenization," Bio-protocol, vol. 16, no. 8, 2026.

[40] P. Choudhri, M. Rani, R. Sangwan, R. Kumar, A. Kumar, and V. Chhokar, "*De novo* sequencing, assembly and characterisation of *Aloe vera* transcriptome and analysis of expression profiles of genes related to saponin and anthraquinone metabolism," BMC Genomics, vol. 19, no. 427, 2018, doi: 10.1186/s12864-018-4819-2.

[41] C. Hughes, S. Moggridge, T. Müller, P. Sorensen, G. Morin, and J. Krijgsveld, "Single-pot, solid-phase-enhanced sample preparation for proteomics experiments," Protocols, vol. 14, pp. 68–85, 2018, doi: 10.1038/s41596-018-0082-x.

[42] B. Haas et al., "*De novo* transcript sequence reconstruction from RNA-seq using the Trinity platform for reference generation and analysis," Protocols, vol. 8, pp. 1494–1512, 2013, doi: 10.1038/nprot.2013.084.

[43] TheGalaxyCommunity, "The Galaxy platform for accessible, reproducible, and collaborative data analyses: 2024 update," Nucleic Acids Research, vol. 52, no. W1, pp. W83-W94, 2024, doi: 10.1093/nar/gkae410.

[44] Y. Zhang, Lan, M., Chen, Y., "Minimal Information for Studies of Extracellular Vesicles (MISEV): Ten-Year Evolution (2014–2023)," Pharmaceutics, vol. 16, no. 11, p. 1394, 2024, doi: 10.3390/pharmaceutics16111394.

[45] Z. Sun, Zheng, Y., Wang, T., Zhang, J., Li, J., Wu, Z., Zhang, F., Gao, T., Yu, L., Xu, X., Qian, H., Tan, Y., "Aloe Vera Gel and Rind-Derived Nanoparticles Mitigate Skin Photoaging via Activation of Nrf2/ARE Pathway," International Journal of Nanomedicine, vol. 20, pp. 4051–4067, 2025, doi: 10.2147/IJN.S510352.

[46] V. Syromiatnikova, Prokopeva, A., Gomzikova, M., "Methods of the Large-Scale Production of Extracellular Vesicles," International Journal of Molecular Sciences, vol. 23, no. 18, p. 10522, 2022, doi: 10.3390/ijms231810522.

[47] M. Nemati et al., "Plant-derived extracellular vesicles: a novel nanomedicine approach with advantages and challenges," Cell Communication and Signaling, vol. 20, no. 69, 2022, doi: 10.1186/s12964-022-00889-1.

[48] Y. Zhu et al., "Applications of plant-derived extracellular vesicles in medicine," MedComm, vol. 5, no. 10, p. e741, 2024, doi: 10.1002/mco2.741.

[49] J. Zimmerman, Verboonen, B., Hanson, A., Arballo, L., Brusslan, J., "Arabidopsis apoplast TET8 positively correlates to leaf senescence, and tet3tet8 double mutants are delayed in leaf senescence," Plant Direct, vol. 8, no. 9, p. e70006, 2024.

[50] Q. Cai, Qiao, L., Wang, M., He, B., Lin, F., Palmquist, J., Huang, S., Jin, H., "Plants send small RNAs in extracellular vesicles to fungal pathogen to silence virulence genes," Science, vol. 360, no. 6393, pp. 1126–1129, 2018.

[51] F. Wang, Muto, A., Velde, J., Neyt, P., Himanen, K., Vandepoele, K., Lijsebettens, M., "Functional Analysis of the Arabidopsis TETRASPANIN Gene Family in Plant Growth and Development," Plant Physiology, vol. 169, no. 3, pp. 2200–2214, 2015.

[52] Y. Yang, Wu, J., Xia, J., Wan, Y., Xu, J., Zhang, L., Liu, D., Chen, L., Tang, F., Ao, H., Peng, C., "Can aloin develop to medicines or healthcare products?," Biomedicine & Pharmacotherapy, vol. 153, p. 113421, 2022, doi: 10.1016/j.biopha.2022.113421.

[53] Y. He, Xi, J., Fang, J., Zhang, B., Cai, W., "Aloe-emodin alleviates doxorubicin-induced cardiotoxicity via inhibition of ferroptosis," Free Radical Biology and Medicine, vol. 206, pp. 13–21, 2023, doi: 10.1016/j.freeradbiomed.2023.06.025.

[54] B. Salehi, Albayrak, S., Antolak, H., Kręgiel, D., Pawlikowska, E., Sharifi-Rad, M., Uprety, Y., Fokou, P., Yousef, Z., Zakaria, Z., Varoni, E., Sharopov, F., Martins, N., Iriti, M., Sharifi-Rad, J., "Aloe Genus Plants: From Farm to Food Applications and Phytopharmacotherapy," International Journal of Molecular Sciences, vol. 19, no. 9, p. 2843, 2018, doi: 10.3390/ijms19092843.

[55] M. Bingqin, Weiyun, Q., Zhenguo, L., Liangzhi, W., "Effect of barbaloin on the NOX4/ROS/p38 MAPK signaling pathway and podocyte function in rats with diabetic nephropathy," Chinese Journal of Comparative Medicine, vol. 30, no. 9, pp. 1–7, 2020, doi: 10.3969/j.issn.1671-7856.2020.09.001.

[56] M. Kumar, Sharma, S., Kumar, J., Barik, S., Mazumder, S., "Mitochondrial electron transport chain in macrophage reprogramming: Potential role in antibacterial immune response," Current Research in Immunology, vol. 5, p. 100077, 2024, doi: 10.1016/j.crimmu.2024.100077.

[57] S. Pérez, Rius-Pérez, S., "Macrophage Polarization and Reprogramming in Acute Inflammation: A Redox Perspective," Antioxidants, vol. 11, no. 7, p. 1394, 2022, doi: 10.3390/antiox11071394.

[58] A. Viola, Munari, F., Sánchez-Rodríguez, R., Scolaro, T., Castegna, A., "The Metabolic Signature of Macrophage Responses," Frontiers in Immunology, vol. 10, 2019, doi: 10.3389/fimmu.2019.01462.

[59] R. Orihuela, McPherson, C., Harry, G., "Microglial M1/M2 polarization and metabolic states," British Journal of Pharmacology, vol. 173, no. 4, pp. 649–665, 2015, doi: 10.1111/bph.13139.

[60] D. Lavogina, Lust, H., Tahk, M-J., Laasfeld, T., Vellama, H., Nasirova, N., Vardja, M., Eskla, K-L., Salumets, A., Rinken, A., Jaal, J., "Revisiting the Resazurin-Based Sensing of Cellular Viability: Widening the Application Horizon," Biosensors, vol. 12, no. 4, p. 196, 2022, doi: 10.3390/bios12040196.

[61] C. Riendeau, Kornfeld, H., "THP-1 Cell Apoptosis in Response to Mycobacterial Infection," Infection and Immunity, vol. 71, no. 1, pp. 254–259, 2003, doi: 10.1128/IAI.71.1.254-259.2003.

[62] D. Korns, Frasch, S., Fernandez-Boyanapalli, R., Henson, P., Bratton, D., "Modulation of macrophage efferocytosis in inflammation," vol. 2, 2011, doi: 10.3389/fimmu.2011.00057.

[63] S. Wang, Liang, Y., Dai, C., "Metabolic Regulation of Fibroblast Activation and Proliferation during Organ Fibrosis," Kidney Diseases, vol. 8, no. 2, pp. 115–125, 2022, doi: 10.1159/000522417.

[64] K. Bernard, Logsdon, N., Ravi, S., Xie, N., Persons, B.., Rangarajan, S., Zmijewski, J., Mitra, K., Liu, G., Darley-Usmar, V., Thannickal, V., "Metabolic Reprogramming Is Required for Myofibroblast Contractility and Differentiation," Journal of Biological Chemistry, vol. 290, no. 42, pp. 25427–25438, 2015, doi: 10.1074/jbc.M115.646984.

[65] F. Wang, Liu, J., An, Q., Wang, Y., Yang, Y., Huo, T., Yang, S., Ju, R., Quan, Q., "Aloe Extracts Inhibit Skin Inflammatory Responses by Regulating NF-κB, ERK, and JNK Signaling Pathways in an LPS-Induced RAW264.7 Macrophages Model," Clinical, Cosmetic and Investigational Dermatology, vol. 16, pp. 267–278, 2023, doi: 10.2147/CCID.S391741.

[66] J. Yadav, Verma, A., Pathak, P., Dwivedi, A., Singh, A., Kumar, P., Khalilullah, H., Jaremko, M., Emwas, A-H., Patel, D., "Phytoconstituents as modulators of NF-κB signalling: Investigating therapeutic potential for diabetic wound healing," Biomedicine & Pharmacotherapy, vol. 177, p. 117058, 2024, doi: 10.1016/j.biopha.2024.117058.

[67] R. Eskandani, Kazempour, M., Farahzadi, R., Sanaat, Z., Eskandani, M., Adibkia, K., Vandghanooni, S., Mokhtarzadeh, A., "Engineered nanoparticles as emerging gene/drug delivery systems targeting the nuclear factor-κB protein and related signaling pathways in cancer," Biomedicine & Pharmacotherapy, vol. 156, p. 113932, 2022, doi: 10.1016/j.biopha.2022.113932.

[68] H. Svitina, Hamman, J., Gouws, C., "Molecular mechanisms and associated cell signalling pathways underlying the anticancer properties of phytochemical compounds from Aloe species (Review)," Experimental and Therapeutic Medicine, vol. 22, no. 2, p. 852, 2021, doi: 10.3892/etm.2021.10284.

[69] H. Sapan, Paturusi, I., Jusuf, I., Patellongi, I., Massi, M., Pusponegoro, A., Arief, S., Labeda, I., Islam, A., Rendy, L., Hatta, M., "Pattern of cytokine (IL-6 and IL-10) level as inflammation and anti-inflammation mediator of multiple organ dysfunction syndrome (MODS) in polytrauma," International Journal of Burns and Trauma, vol. 6, no. 2, pp. 37–43, 2016.

[70] L. Luckett-Chastain, Calhoun, K., Schartz, T., Gallucci, R., "IL-6 influences the balance between M1 and M2 macrophages in a mouse model of irritant contact dermatitis," The Journal of Immunology, vol. 196, no. 1, 2016, doi: 10.4049/jimmunol.196.Supp.196.17.

[71] C. Niemand, Nimmesgern, A., Haan, S., Fischer, P., Schaper, F., Rossaint, R., Heinrich, P., Müller-Newen, G., "Activation of STAT3 by IL-6 and IL-10 in Primary Human Macrophages Is Differentially Modulated by Suppressor of Cytokine Signaling 3," The Journal of Immunology, vol. 170, no. 6, pp. 3263–3272, 2003, doi: 10.4049/jimmunol.170.6.3263.

[72] X.-L. Fu, Duan, W., Su, C-Y., Mao, F-Y., Lv, Y-P., Teng, Y-S., Yu, P-W., Zhuang, Y., Zhao, Y-L., "Interleukin 6 induces M2 macrophage differentiation by STAT3 activation that correlates with gastric cancer progression," Cancer Immunoly, Immunotherapy, vol. 12, pp. 1597–1608, 2017, doi: 10.1007/s00262-017-2052-5.

[73] H. Zhou et al., "Aloe-derived vesicles enable macrophage reprogramming to regulate the inflammatory immune environment," Frontiers in Bioengineering Biotechnology, vol. 11, p. 1339941, 2023, doi: 10.3389/fbioe.2023.1339941.

[74] S. Gordon, Martinez, F., "Alternative Activation of Macrophages: Mechanism and Functions," Immunity, vol. 32, no. 5, pp. 593–604, 2010, doi: 10.1016/j.immuni.2010.05.007.

[75] Y. Jiang, Cai, R., Huang, Y., Zhu, L., Xiao, L., Wang, C., Wang, L., "Macrophages in organ fibrosis: from pathogenesis to therapeutic targets," Cell Death Discovery, vol. 10, no. 487, 2024, doi: 10.1038/s41420-024-02247-1.

[76] T. Wynn, Vannella, K., "Macrophages in tissue repair, regeneration, and fibrosis," Immunity, vol. 44, no. 3, pp. 450–462, 2016, doi: 10.1016/j.immuni.2016.02.015.

[77] K. Kim, Sheppard, D., Chapman, H., "TGF-β1 Signaling and Tissue Fibrosis," Cold Spring Harbor Perspectives in Biology, vol. 10, no. 4, p. a022293, 2018, doi: 10.1101/cshperspect.a022293.

[78] S. Dikici, "Ascorbic Acid Enhances the Metabolic Activity, Growth and Collagen Production of Human Dermal Fibroblasts Growing in Three-dimensional (3D) Culture," Journal of Science, vol. 36, no. 4, pp. 1625–1637, 2023, doi: 10.35378/gujs.1040277.

[79] H. Li, Yang, L., Zhang, Y., Gao, Z., "Kaempferol inhibits fibroblast collagen synthesis, proliferation and activation in hypertrophic scar via targeting TGF-β receptor type I," Biomedicine & Pharmacotherapy, vol. 83, no. 8, pp. 967–974, 2016, doi: 10.1016/j.biopha.2016.08.011.

[80] E. Jo, Park, S., Choi, Y., Jeon, W-K., Kim, B-C., "Kaempferol Suppresses Transforming Growth Factor-β1–Induced Epithelial-to-Mesenchymal Transition and Migration of A549 Lung Cancer Cells by Inhibiting Akt1-Mediated Phosphorylation of Smad3 at Threonine-179," Neoplasia, vol. 17, no. 7, pp. 525–537, 2015, doi: 10.1016/j.neo.2015.06.004.

[81] F. Dou, Liu, Y., Liu, L., Wang, J., Sun, T., Mu, F., Guo, Q., Guo, C., Jia, N., Liu, W., Yi, D., Wen, A., "Aloe-Emodin Ameliorates Renal Fibrosis Via Inhibiting PI3K/Akt/mTOR Signaling Pathway In Vivo and In Vitro," Rejuvenation Research, vol. 22, no. 3, 2018, doi: 10.1089/rej.2018.2104.

[82] H. Y. Liu, C., Wu, X., Peng, S., Zhou, L., McClements, D., Liu, W., "Influence of the maturity on the characteristics of orange-derived extracellular vesicles and their delivery performance for curcumin," Food Chemistry, vol. 485, p. 144518, 2025, doi: 10.1016/j.foodchem.2025.144518.

[83] A. Eliades. "Deep Green Permaculture." https://deepgreenpermaculture.com/2019/04/16/identifying-and-growing-edible-aloe-vera/#:~:text=A%20more%20definite%20way%20to,vera%20barbadensis%20Miller%20behind%20it. (accessed 05/05, 2025).

[84] M. Hęś, Dziedzic, K., Górecka, D., Jędrusek-Golińska, A., Gujska, E., "Aloe vera (L.) Webb.: Natural Sources of Antioxidants – A Review," vol. 74, pp. 255–265, 2019.

[85] A. Martínez-Sánchez, López-Cañavate, M., Guirao-Martínez, J., Roca, M., Aguayo, E., "Aloe vera Flowers, a Byproduct with Great Potential and Wide Application, Depending on Maturity Stage," Food, vol. 9, no. 11, p. 1542, 2020, doi: 10.3390/foods9111542.

[86] E. Mensah, Adadi, P., Asase, R., Kelvin, O., Mozhdehi, F., Amoah, I., Agyei, D., "Aloe vera and its byproducts as sources of valuable bioactive compounds: Extraction, biological activities, and applications in various food industries," PharmaNutrition, vol. 31, p. 100436, 2025, doi: 10.1016/j.phanu.2025.100436.

[87] S. Sonam, Tiwari, A., "Phytochemical Evaluation of Different Aloe Species," International Journal of Chemical and Pharmaceutical Analysis, vol. 3, no. 4, 2016, doi: 10.21276/ijcpa.

[88] S. Kumar, Yadav, A., Yadav, M., Yadav, J., "Effect of climate change on phytochemical diversity, total phenolic content and in vitro antioxidant activity of Aloe vera (L.) Burm.f.," BMC Research Notes, vol. 10, no. 60, 2017, doi: 10.1186/s13104-017-2385-3.

[89] O. Genest, S. Wickner, and S. Doyle, "HSP90 and HSP70 chaperones: collaborators in protein remodeling," Journal of Biological Chemistry, vol. 294, no. 6, pp. 2109–2120, 2019, doi: 10.1074/jbc.REV118.002806.

[90] J. Höhfeld, D. Cyr, and C. Patterson, "From the cradle to the grave: molecular chaperones that may choose between folding and degradation," EMBO Reports, vol. 2, no. 10, pp. 885–890, 2001, doi: 10.1093/embo-reports/kve206.

[91] T. Svitkina "The Actin Cytoskeleton and Actin-Based Motility," Cold Spring Harb Perspectives in Biology, vol. 10, no. 1, p. a018267, 2018, doi: 10.1101/cshperspect.a018267.

[92] M. Skruzny, T. Brach, R. Ciuffa, S. Rybina, M. Wachsmuth, and M. Kaksonen "Molecular basis for coupling the plasma membrane to the actin cytoskeleton during clathrin-mediated endocytosis," Cell Biology, vol. 109, no. 38, pp. E2533–E2542, 2012, doi: 10.1073/pnas.1207011109.

[93] E. Coudrier and C. Almeida "Myosin 1 controls membrane shape by coupling F-Actin to membrane," vol. 1, no. 5, pp. 230–235, 2011, doi: 10.4161/bioa.18406.

[94] M. Mettlen, P. Chen, S. Srinivasan, G. Danuser, and S. Schmid "Regulation of Clathrin-Mediated Endocytosis," Annual Review of Biochemistry vol. 16, no. 87, pp. 871–896, 2018, doi: 10.1146/annurev-biochem-062917-012644.

[95] P. McPherson, B. Ritter, and B. Wendland, "Clathrin-Mediated Endocytosis," in Madame Curie Bioscience Database [Internet]: Landes Bioscience, 2000-2013.

[96] L. Blanc and M. Vidal, "New insights into the function of Rab GTPases in the context of exosomal secretion," Small GTPases, vol. 9, no. 1-2, pp. 95–106, 2017, doi: 10.1080/21541248.2016.1264352.

[97] V. Hyenne, M. Labouesse, and J. Goetz "The Small GTPase Ral orchestrates MVB biogenesis and exosome secretion," Small GTPases, vol. 9, no. 6, pp. 445–451, 2016, doi: 10.1080/21541248.2016.1251378.

[98] S. Krylova and D. Feng, "The Machinery of Exosomes: Biogenesis, Release, and Uptake," International Journal of Molecular Science, vol. 24, no. 2, p. 1337, 2023, doi: 10.3390/ijms24021337.

[99] R. Prekeris, B. Yang, V. Oorschot, J. Klumperman, and R. Scheller, "Differential Roles of Syntaxin 7 and Syntaxin 8 in Endosomal Trafficking," Molecular Biology of the Cell, vol. 10, no. 11, 2017, doi: 10.1091/mbc.10.11.3891.

[100] F. Teng, Y. Wang, and B. Tang, "The syntaxins," Genome Biology, vol. 2, no. 11, 2001, doi: 10.1186/gb-2001-2-11-reviews3012.

[101] L. Mulcahy, R. Pink, and D. Carter "Routes and mechanisms of extracellular vesicle uptake," Journal of Extracellular Vesicles, vol. 3, 2014, doi: 10.3402/jev.v3.24641.

[102] D. Konopka-Postupolska and G. Clark, "Annexins as Overlooked Regulators of Membrane Trafficking in Plant Cells," International Journal of Molecular Science, vol. 18, no. 4, p. 863, 2017, doi: 10.3390/ijms18040863.

[103] V. Gerke et al., "Annexins—a family of proteins with distinctive tastes for cell signaling and membrane dynamics," Nature Communications, vol. 15, p. 1574, 2024, doi: 10.1038/s41467-024-45954-0.

[104] L. Gadipudi et al., "Annexin A1 treatment prevents the evolution to fibrosis of experimental nonalcoholic steatohepatitis," Clinical Science (London), vol. 136, no. 9, pp. 643–656, 2022, doi: 10.1042/CS20211122.

[105] J. Fan, N. Luo, G. Liu, X. Xu, S. Li, and X. Lv, "Mechanism of annexin A1/N-formylpeptide receptor regulation of macrophage function to inhibit hepatic stellate cell activation through Wnt/β-catenin pathway," World Journal of Gastroenterology, vol. 29, no. 22, pp. 3422–3439, 2023, doi: 10.3748/wjg.v29.i22.3422.

[106] P. Tu et al., "Mechanical Stretch Promotes Macrophage Polarization and Inflammation via the RhoA-ROCK/NF-*κ* B Pathway," BioMed Research International, vol. 2022, p. 6871269, 2022, doi: 10.1155/2022/6871269.

[107] N. Manickam, M. Patel, K. Griendling, Y. Gorin, and J. Barnes, "RhoA/Rho kinase mediates TGF-β1-induced kidney myofibroblast activation through Poldip2/Nox4-derived reactive oxygen species," American Journal of Physiology-Renal Physiology, vol. 307, no. 2, pp. F159–71, 2014, doi: 10.1152/ajprenal.00546.2013.

[108] D. Xu, H. Kishi, H. Kawamichi, K. Kajiya, Y. Takada, and S. Kobayashi, "Sphingosylphosphorylcholine induces stress fiber formation via activation of Fyn-RhoA-ROCK signaling pathway in fibroblasts," Cell Signaling, vol. 24, no. 1, pp. 282–289, 2012, doi: 10.1016/j.cellsig.2011.09.013.

[109] K. Katoh, K. Yumiko, and Y. Noda, "Rho-associated kinase-dependent contraction of stress fibres and the organization of focal adhesions," Journal of the Royal Society Interface, vol. 8, no. 56, pp. 305–311, 2010, doi: 10.1098/rsif.2010.0419.

[110] A. Rinne and F. Pluteanu, "Ca 2+ Signaling in Cardiovascular Fibroblasts," Biomolecules vol. 14, no. 11, p. 1365, 2024, doi: 10.3390/biom14111365.

[111] L. Johnson, E. Rodansky, A. Haak, S. Larsen, R. Neubig, and P. Higgins, "Novel Rho/MRTF/SRF inhibitors block matrix-stiffness and TGF-β-induced fibrogenesis in human colonic myofibroblasts," Inflammatory Bowel Disease, vol. 20, no. 1, pp. 154–165, 2014, doi: 10.1097/01.MIB.0000437615.98881.31.

[112] E. Small, "The actin-MRTF-SRF gene regulatory axis and myofibroblast differentiation," Journal of Cardiovascular Translational Research, vol. 5, no. 6, pp. 794–804, 2012, doi: 10.1007/s12265-012-9397-0.

[113] J. Andersen, A. Vestergaard, S. Mikkelsen, L. Mogensen, M. Chalat, and R. Molday, "P4-ATPases as Phospholipid Flippases-Structure, Function, and Enigmas," Frontier in Physiology, vol. 7, p. 275, 2016, doi: 10.3389/fphys.2016.00275.

[114] J. Gao, Y. Hao, X. Piao, and X. Gu, "Aldehyde Dehydrogenase 2 as a Therapeutic Target in Oxidative Stress-Related Diseases: Post-Translational Modifications Deserve More Attention," International Journal of Molecular Sciences, vol. 23, no. 5, p. 2682, 2022, doi: 10.3390/ijms23052682.

[115] M. Breitzig, C. Bhimineni, R. Lockey, and N. Kolliputi, "4-Hydroxy-2-nonenal: a critical target in oxidative stress?," American Journal of Physiology-Cell Physiology, vol. 311, no. 4, pp. C537-C543, 2016, doi: 10.1152/ajpcell.00101.2016.

[116] H. Zheng, L. Zhang, C. Wang, Y. Wang, and C. Zeng, "Metabolic dysregulation in pulmonary fibrosis: insights into amino acid contributions and therapeutic potential," C ell Death Discovery, vol. 11, no. 411, 2025, doi: 10.1038/s41420-025-02715-2.

[117] J. Lee et al., "Role of lung ornithine aminotransferase in idiopathic pulmonary fibrosis: regulation of mitochondrial ROS generation and TGF-β1 activity," E xperimental & Molecular Medicine, vol. 56, pp. 478–490, 2024, doi: 10.1038/s12276-024-01170-w.

[118] E. Karna, L. Szoka, T. Huynh, and J. Palka, "Proline-dependent regulation of collagen metabolism," Cellular and Molecular Life Sciences, vol. 77, no. 10, pp. 1911–1918, 2020, doi: 10.1007/s00018-019-03363-3.

[119] J. Albina, J. Abate, and B. Mastrofrancesco, "Role of Ornithine as a Proline Precursor in Healing Wounds," Journal of Surgical Research, vol. 55, no. 1, pp. 97–102, 1993, doi: 10.1006/jsre.1993.1114.

[120] C. Chang et al., "Ribosome Biogenesis Serves as a Therapeutic Target for Treating Endometriosis and the Associated Complications," Biomedicines, vol. 10, no. 1, p. 185, 2022, doi: 10.3390/biomedicines10010185.

[121] S. Zhou, X. Yin, M. Mayr, M. Noor, P. Hylands, and Q. Xu, "Proteomic landscape of TGF-β1-induced fibrogenesis in renal fibroblasts," S cientific Reports, vol. 10, no. 19054, 2020, doi: 10.1038/s41598-020-75989-4.

[122] J. Wu et al., "Glutamyl-Prolyl-tRNA Synthetase Regulates Proline-Rich Pro-Fibrotic Protein Synthesis During Cardiac Fibrosis," vol. 127, no. 6, pp. 827–846, 2021, doi: 10.1161/CIRCRESAHA.119.315999.

[123] M. Jové et al., "Ether Lipid-Mediated Antioxidant Defense in Alzheimer’s Disease," Antioxidants, vol. 12, no. 2, p. 293, 2023, doi: 10.3390/antiox12020293.

[124] M. Youssef, A. Ibrahim, K. Akashi, and M. Hossain, "PUFA-Plasmalogens Attenuate the LPS-Induced Nitric Oxide Production by Inhibiting the NF-kB, p38 MAPK and JNK Pathways in Microglial Cells," Neuroscience, vol. 397, pp. 18-30, 2019, doi: 10.1016/j.neuroscience.2018.11.030.

[125] F. Ali, M. Hossain, S. Sejimo, and K. Akashi, "Plasmalogens Inhibit Endocytosis of Toll-like Receptor 4 to Attenuate the Inflammatory Signal in Microglial Cells," Molecular Neurobiology, vol. 56, no. 5, pp. 3404–3419, 2019, doi: 10.1007/s12035-018-1307-2.

[126] M. Sisto and S. Lisi, "Targeting Interleukin-17 as a Novel Treatment Option for Fibrotic Diseases," Journal of Clinical Medicine, vol. 13, no. 1, p. 164, 2023, doi: 10.3390/jcm13010164.

[127] M. Wilson et al., "Bleomycin and IL-1beta-mediated pulmonary fibrosis is IL-17A dependent," Journal of Experimental Medicine, vol. 207, no. 3, pp. 535–552, 2010, doi: 10.1084/jem.20092121.

[128] S. Chatterjee, "Endothelial Mechanotransduction, Redox Signaling and the Regulation of Vascular Inflammatory Pathways," Frontiers in Physiology, vol. 9, p. 524, 2018, doi: 10.3389/fphys.2018.00524.

[129] H. Hsieh, C. Liu, B. Huang, A. Tseng, and D. Wang, "Shear-induced endothelial mechanotransduction: the interplay between reactive oxygen species (ROS) and nitric oxide (NO) and the pathophysiological implications," Journal of Biomedical Science, vol. 21, no. 1, p. 3, 2014, doi: 10.1186/1423-0127-21-3.

[130] D. Zheng, T. Liwinski, and E. Elinav, "Inflammasome activation and regulation: toward a better understanding of complex mechanisms," vol. 6, no. 36, 2020, doi: 10.1038/s41421-020-0167-x.

[131] P. Broz and V. Dixit, "Inflammasomes: mechanism of assembly, regulation and signalling," Nature Reviews Immunology, vol. 16, pp. 407–420, 2016, doi: 10.1038/nri.2016.58.

[132] H. Asard, R. Barbaro, P. Trost, and A. Bérczi, "Cytochromes b561: ascorbate-mediated trans-membrane electron transport," Antioxidant Redox Signaling vol. 19, no. 9, pp. 1026–1035, 2013, doi: 10.1089/ars.2012.5065.

[133] M. Manresa et al., "Hydroxylase inhibition regulates inflammation-induced intestinal fibrosis through the suppression of ERK-mediated TGF-β1 signaling," American Journal of Physiology Gastrointestinal and Liver Physiology, vol. 311, no. 6, pp. G1076-G1090, 2016, doi: 10.1152/ajpgi.00229.2016.

[134] G. Zhang and S. Ghosh, "Toll-like receptor–mediated NF-κB activation: a phylogenetically conserved paradigm in innate immunity," Journal of Clinical Investigation, vol. 107, no. 1, pp. 13–19, 2001, doi: 10.1172/JCI11837.

[135] A. Bachstetter and L. Eldik, "The p38 MAP Kinase Family as Regulators of Proinflammatory Cytokine Production in Degenerative Diseases of the CNS," Aging and Disease, vol. 1, no. 3, pp. 199–211, 2010.

[136] T. Fukao and S. Koyasu, "PI3K and negative regulation of TLR signaling," Trends in Immunology, vol. 24, no. 7, pp. 358–363, 2003, doi: 10.1016/s1471-4906(03)00139-x.

[137] T. Antoniv and L. Ivashkiv, "Interleukin-10-induced gene expression and suppressive function are selectively modulated by the PI3K-Akt-GSK3 pathway," Immunology, vol. 132, no. 4, pp. 567–577, 2011, doi: 10.1111/j.1365-2567.2010.03402.x.

[138] T. Liu, Y. Li, M. Xu, H. Huang, and Y. Luo, "PRMT2 silencing regulates macrophage polarization through activation of STAT1 or inhibition of STAT6," BMC Immunology, vol. 25, no. 1, 2024, doi: 10.1186/s12865-023-00593-w.

